# Hierarchically ordered multi-timescale structural dynamics of the intrinsically disordered p53 transactivation domain

**DOI:** 10.1101/2025.08.01.668138

**Authors:** Dániel Szöllősi, Supriya Pratihar, Dwaipayan Mukhopadhyay, Ashok Kumar Rout, G. Jithender Reddy, Niklas Ebersberger, Stefan Becker, Gábor Nagy, Sarah Rauscher, Donghan Lee, Reinhard Klement, Christian Griesinger, Helmut Grubmüller

## Abstract

Intrinsically disordered proteins (IDPs) exhibit pronounced structural dynamics, which is crucial for their functional versatility. Yet their dynamics slower than nanoseconds remain largely elusive. We combined high-power relaxation dispersion nuclear magnetic resonance spectroscopy with molecular dynamics simulations to characterize these kinetics and the underlying structural interconversions of a prototypical IDP, the N-terminal transactivation domain of the tumor suppressor p53 (p53-TAD). We find a complex hierarchy of structural dynamics on timescales covering over seven orders of magnitude, ranging from fast nanoseconds backbone re-orientations, via sub-microsecond helix-formation dynamics involving many structural sub-states and transition times, to transient tertiary structure formation slower than 25 microseconds. These rich structural dynamics of p53-TAD, and likely those of other IDPs, parallel the timescale hierarchy of the conformational dynamics of folded proteins.

**One-Sentence summary:** A hierarchical energy landscape governs kinetics and structural dynamics of the disordered p53 transactivation domain.

## Introduction

Folded proteins exhibit complex structural dynamics across a wide range of timescales, covering picoseconds to milliseconds or even longer which often implements or modulates their function (*1–4*). These dynamics are governed by a hierarchically structured (*5, 2, 6*) and funnel-shaped free energy landscape (*7*), characterized by a hierarchy of energy barrier “tiers” (*8*). Detailed knowledge of the distribution of these energy barriers, the resulting kinetics, and the underlying structural dynamics are therefore key to our understanding of how proteins fold and how they perform their remarkably broad range of biochemical functions on the molecular level. Intrinsically disordered peptides and proteins (IDPs), in contrast, lack such a stable native structure (*9, 10*). Rather, they rapidly interconvert between many different structures, with only transiently formed secondary structure elements (*11*) and long-range contacts (*12*). Operating at the limits of Anfinsen’s dogma (*13*), these fast reconfiguration dynamics (*14, 15*) lead to the general notion that the underlying free energy landscape is rather shallow and unstructured (*16*).

IDPs pose considerable experimental and computational challenges, and hence information on their structural dynamics and specifically their kinetics is still sparse. For example, single-molecule spectroscopy experiments revealed fast (ns) reconfiguration dynamics of several IDPs (*14, 15, 17*), but no atomistic details of the underlying structural dynamics. Kinetic information at fast (ps to ns) timescales is also provided by nuclear magnetic resonance (NMR) relaxation measurements (*18, 19*), whereas NMR relaxation dispersion (RD) measurements typically probe kinetics at several tens of μs or slower (*20*). Atomistic molecular dynamics (MD) simulations provide more direct structural insights, but typically only quantify sub-μs structural dynamics (*21–24*). Hence, the structural dynamics and kinetics of IDPs particularly on timescales between 100 ns and 10 μs are largely uncharted territory.

Here we combined high-power NMR RD measurements and milliseconds atomistic MD simulations to gain access to these 100 ns to 10 μs kinetics and structural dynamics of the intrinsically disordered N-terminal transactivation domain of the tumor protein p53 (p53-TAD, Fig. 1A). Also known as the “guardian of the genome”, p53 regulates the cellular response to genomic damages and thus prevents cancer formation (*25*). As one of the most important signaling hubs, p53-TAD adopts different structures upon specific binding to a stunningly large number of proteins (*26, 27*). X-ray structures of the p53/MDM2 complex (*26*), NMR measurements (*11*), and MD simulations (*21*) suggest that the TAD in solution forms transient helical structures (“helix 1” and “helix 2” in Fig. 1A) (*28*). These helices cooperatively engage in binding, can adopt well-folded conformations upon interaction with binding partners (*26, 19*), and affect the autoinhibition of p53 (*29, 30*). Further, their stability in the unbound state modulates the lifetime of the bound complex (*31, 32*). The structural dynamics of this prototypical IDP might therefore regulate and accelerate the promiscuous yet selective binding of p53-TAD via conformational selection (*28, 33, 34*), yet much of its kinetics and the underlying structural dynamics are elusive.

**Fig. 1.**
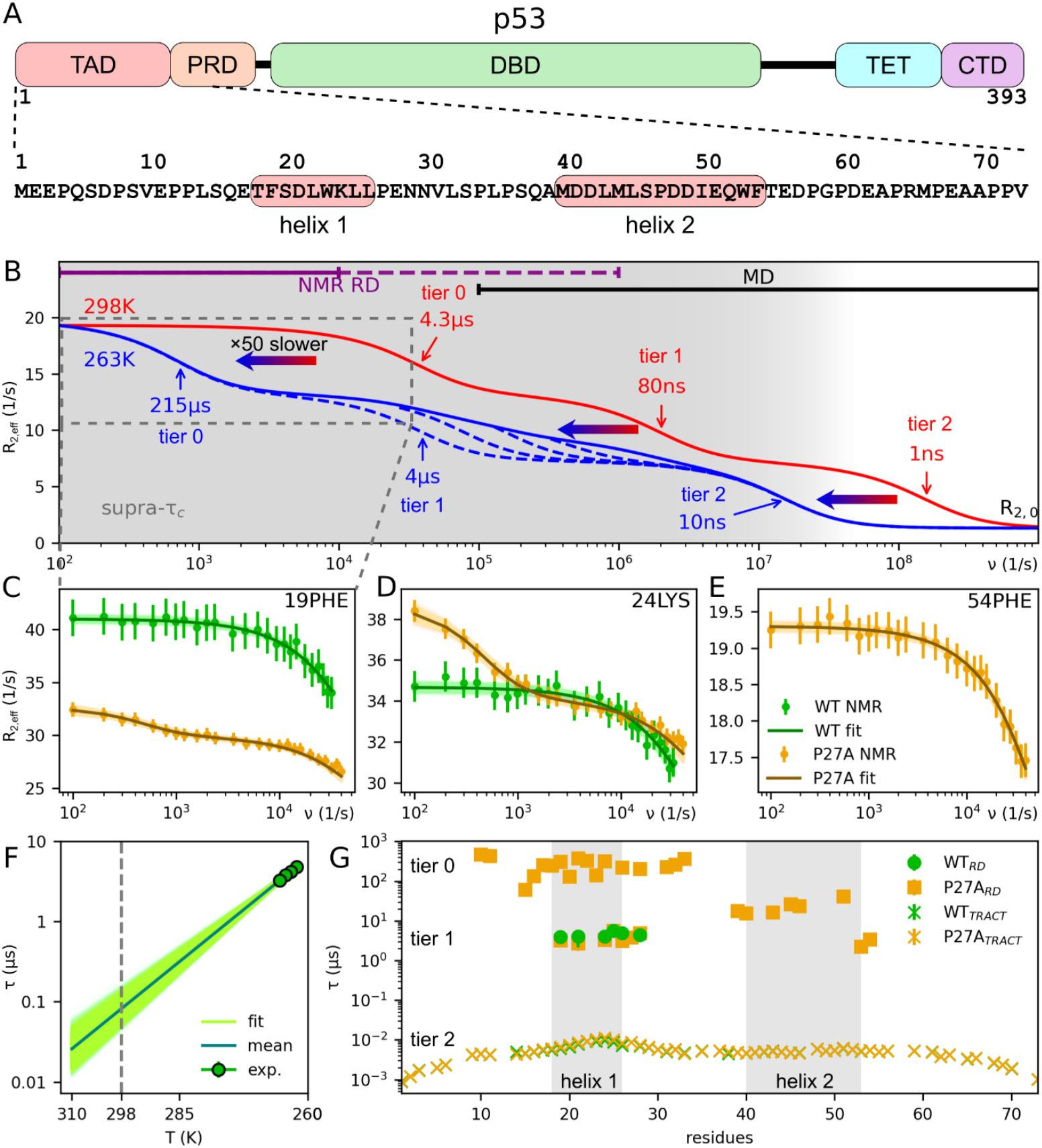
Hierarchical dynamics of p53-TAD across multiple timescales revealed by NMR relaxation dispersion and MD simulations. (**A**) Domain organization of full-length p53 protein and sequence of the p53-TAD with highlighted helical segments. (**B**) Schematic representation of RD profiles revealing three distinct tiers of structural dynamics: slow dynamics (215 μs and 4.3 μs, tier 0), fast dynamics (4 μs and 80 ns, tier 1) and molecular tumbling (10 ns and 1 ns, tier 2) at 263 K (blue) and 298 K (red), respectively. Thin arrows indicate timescales from NMR relaxation experiments scaled by (2π)^−1^ for easier interpretation in terms of the well-known Lorentzian function; at the two different temperatures (blue: 263 K, red: 298 K), similar dynamical processes occur at timescales differing by an Arrhenius factor of ca. 50 (horizontal thick arrows, see **F**). Timescales accessible to high-power NMR RD measurement and MD simulations are indicated as purple and black bars; supercooling renders faster timescales accessible to NMR RD (dashed purple bar). The timescale window of the supra-τ_c_ dynamics is indicated in gray. (**C, D** and **E**) Example amide proton (^1^H_N_) CPMG RD profiles measured at 263 K for three different residues (dots) and fitted CPMG curves (lines) including error bars indicating standard deviation. (**F**) Temperature dependence of relaxation times scales using supercooled NMR ^1^H_N_ *R*_1ρ_ measurements between 262 K and 265 K (green dots, error within symbol size) and Arrhenius extrapolation to 298 K (green line, uncertainty light green). (**G**) Timescales obtained by fitting RD data for all measured residues as shown in C to E. WT RD profiles (green) show a single timescale; several P27A mutant RD profiles show two separate timescales (orange squares); one of which (4 μs) is similar to WT; all residue level tumbling timescales (crosses) are similar for WT and P27A. Error bars are within symbol size; gray shaded areas indicate helix 1 and helix 2.

### NMR reveals rich multi-timescale dynamics of p53-TAD

Figure 1B provides a schematic overview of the timescales and dynamics probed by our combined approach. To access so-called supra-τ_c_ (*35*) dynamics of p53-TAD (residues 1-73) in the low μs range, we have utilized recent advances in high-power RD NMR measurements, in particular, amide proton (^1^H_N_) E-CPMG (CPMG: Carr-Purcell-Meiboom-Gill) (*36*) performed at 1.2 GHz (the part of the red curve indicated by the purple bar). To detect even faster protein motions, we also measured under supercooled conditions at 263 K (*37*) (dashed purple bar) and found dynamics around 4 μs, e.g., for residues 19, 24, and 54 (green curves in Fig. 1C to E, all measured residues shown in fig. S1 and the extracted timescales in table S1). At room temperature (298 K) these supra-τ_c_ dynamics is extrapolated to take place between 66 ns and 100 ns (Fig. 1B, middle red arrow).

Notably, these supra-τ_c_ dynamics are much slower than the nanoseconds reconfiguration dynamics previously observed for IDPs, e.g., by room temperature single-molecule fluorescence resonance energy transfer (FRET) measurements (*14*). They are therefore governed by markedly higher energy barriers, which we will collectively refer to as ‘tier 1’. Incidentally, loop-closure dynamics of p53-TAD measured by photoinduced electron transfer fluorescence correlation spectroscopy (PET-FCS) (*38*) occur at similar timescales. We also observed fast reorientation dynamics (Fig. 1B, right red arrow), governed by lower energy barriers, which we will refer to as ‘tier 2’. Figure 1G shows that these two timescales are measured for many residues (green crosses and circles), suggesting collective structural dynamics.

### Tier 1: Sub-microseconds supra-τ_c_ dynamics of p53-TAD

Which structural motions cause the unexpectedly slow tier 1 dynamics of this IDP? As sketched in Fig. 2A, these dynamics might arise either from the folding and unfolding of transient secondary structure elements (e.g., the helices involving residues 18-26 or 40-55) (*28*) or, alternatively, from the collapse and subsequent dissolving of a hydrophobic patch (*39*) – or due to other unknown structural dynamics.

**Fig. 2.**
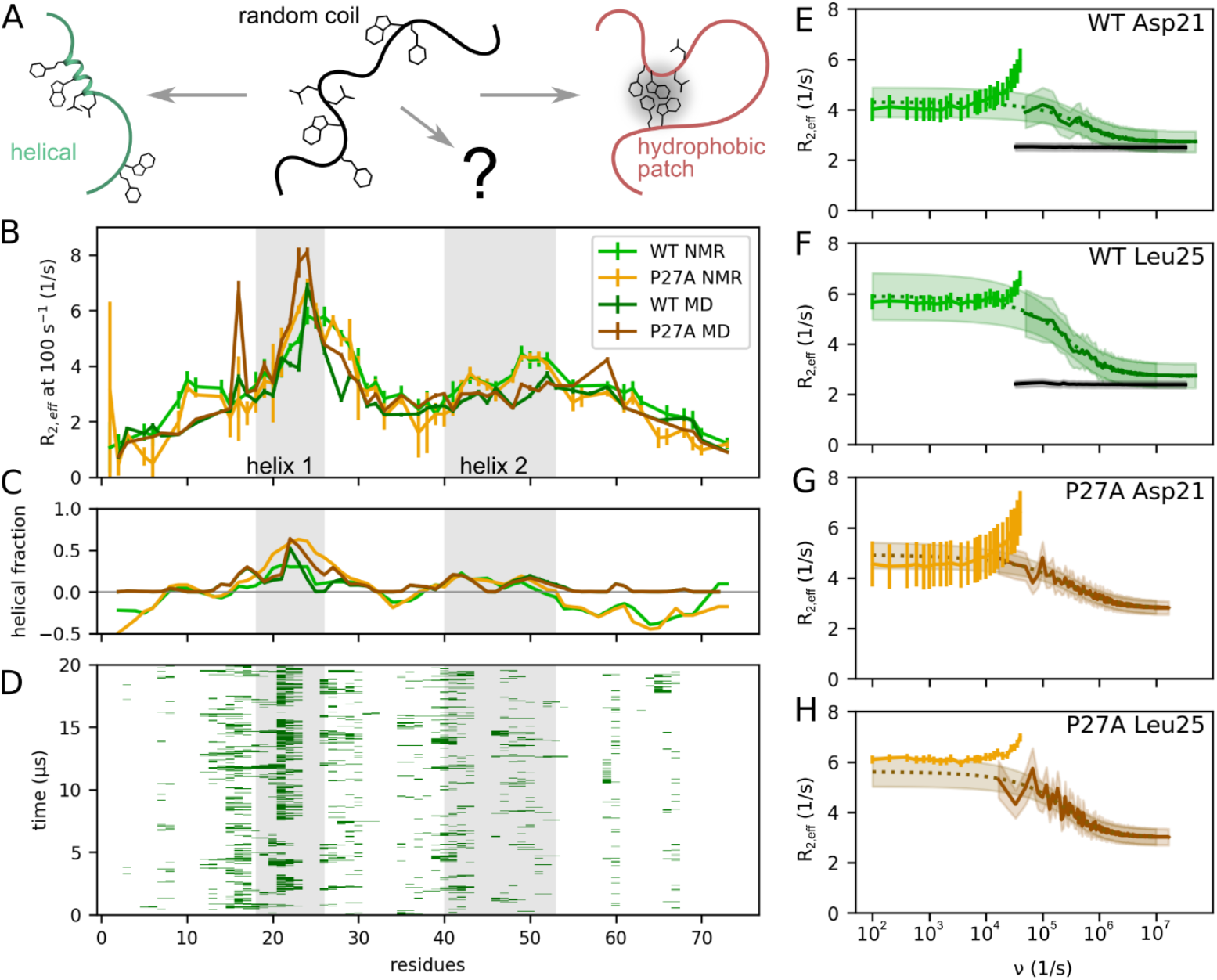
Tier 1 supra-τ_c_ dynamics of p53-TAD as observed by NMR RD and MD simulations. **(A)**Schematic representation of two plausible structural origins of the observed supra-τ_c_ (tier 1) dynamics: folding/unfolding of transient helices and collapse/dissolution of hydrophobic clusters. **(B)** Comparison of amplitudes of NMR RD profiles at 100 s^−1^ CPMG frequency (solid lines) for each measured residue with those amplitudes (dashed) extrapolated from MD simulations (darker colors) for WT (green) and P27A mutant (orange). Experimental and statistical uncertainties are shown as error bars. (**C**) Comparison of the helix fraction of each residue estimated from measured NMR chemical shifts and SSP with those derived from the MD simulations using dihedral angles (colors as in B); helices 1 and 2 are indicated by gray-shading. Negative values indicate a preference for extended conformations. (**D**) Dynamics of helix formation; green bars indicate instances during a representative 20 μs MD simulation at which each residue and its neighbors form a short helical segment. (**E** to **H**) Four examples of measured RD profiles (lighter solid lines) and those calculated from the MD simulations (darker solid) and extrapolated to the NMR frequency range (dotted). As a control, black lines show RD profiles calculated from WT simulations for which unfolding of the α-helix was suppressed. Uncertainties of the measured profiles (vertical bars) were estimated from repeated measurements; for the calculated RD profiles, standard deviations around the mean are indicated as shaded areas.

Evidence for helix-formation dynamics comes from partial helicity seen in our NMR chemical shift measurements, which agree with previously observed (*11*) helical populations between 15% and 30% for the residues within helix 1 (Fig. 2C), and 30 to 60% for the well-described Pro27→Ala mutant (P27A) (*34*). This mutant also shows a markedly increased affinity to MDM2 (*40*), probably because of its higher helix propensity (*31*).

The time scale of the tier 1 supra-τ_c_ dynamics is the same for WT and P27A mutant observed from NMR E-CPMG RD profiles (Fig. 1C, D, G) at 1.2 GHz, yet, a very slow 215 µs kinetics points to an additional process different from tier 1, which is prevalent in the mutant but not in the WT (Fig. 1, C to E, G, orange). This notion is further independently supported by TRACT (TROSY for rotational correlation times) measurements (crosses in Fig. 1G), as explained in the Supplementary Materials, SM (C,F). We will refer to these new dynamics as “tier 0”, the structural origin of which is elusive at this stage.

### Unbiased atomistic simulations validated against NMR

To reveal the structural motions that cause the observed dynamics on all three tiers identified by NMR, we have carried out extensive MD simulations of both p53-TAD WT and P27A mutant in explicit solvent at 298 K. Contrary to previous studies (*41, 42*), these simulations are unbiased and have not been fitted to any p53 measurement, which allowed assessing their accuracy by calculating RD profiles from the MD simulations. Thanks to the total MD trajectory length of 2.4 ms, these calculated RD profiles cover slow timescales up to 10 μs (sketched in Fig. 1B, black bar), thus enabling direct comparison with the measured ones. For a discussion of possible uncertainties and control simulations, see SM (H).

Figure 2, E to H shows measured full RD profiles (solid, lighter colors) and those calculated from the MD simulations (solid, darker colors) for two selected residues of both the WT (Fig. 2, E and F) and the mutant (Fig. 2, G and H; fig. S4-5 show RD profiles for all residues). Note that high-power artefacts cause an unphysical increase of the measured RD profiles at higher frequencies. To enable a most direct comparison, we show Bayesian fits (dotted lines) of a stretched CPMG model (*43*) to the calculated RD profiles, which also did not use the measured RD profiles, see also SM (O, P). This model generalizes the established two-state model (*44*) to many states and, hence, represents a superposition of many relaxation rates. Very good agreement is seen, e.g., for residue Asp21 (Fig. 2E and G), with deviations typical for all residues (Fig. 2B) of ca. 15% exemplified for Leu25 (Fig. 2H), as discussed in SM (H). Further, the helical fractions for all residues calculated from the MD simulations (Fig. 2C, darker colors) also agree well with those derived from the NMR chemical shifts using SSP (*45*) (lighter colors). The MD results also agree with small-angle X-ray scattering, dynamic light scattering, size exclusion chromatography, FRET, Photoinduced Electron Transfer Fluorescence Correlation Spectroscopy (PET-FCS) and Paramagnetic Relaxation Enhancement (PRE) measurements (figs. S12 to S15).

### MD simulations reveal tier 1 helix-formation dynamics with multiple relaxation rates

We conclude that the structural dynamics of p53-TAD are sufficiently accurately described by the simulations to allow identification of the structural motions that give rise to the observed tier 1 dynamics. To this end, Fig. 2D shows the secondary structure dynamics particularly within the helical regions (gray), revealing sub-μs folding and unfolding of short helical structure elements (green bars). These dynamics are indeed the dominant cause of the observed NMR RD dynamics, as evidenced by control simulations, for which the α-helical regions have been forced to remain in helical geometry. Indeed, the RD profiles calculated from these simulations (black curves in Fig. 2, E and F) do not show any tier 1 dynamics. These results establish sub-μs helix folding dynamics as the structural determinant of tier 1, on top of to the known and much faster tier 2 structural reorganization dynamics (*11, 14*) (cf. Fig. 1B).

As can be seen in Fig. 2D, these secondary structure folding dynamics are quite complex and are neither described by a fully cooperative two-state model, nor are the partial folding steps fully uncorrelated. Rather, it seems to involve transitions between many different partially helical conformations and, accordingly, also many timescales.

To characterize these partially cooperative tier 1 dynamics in more detail, we have used the WT MD trajectories to derive a Markov model of the helix 1 conformational dynamics (Fig. 3, SM (R)). Figures 3, A and B show the MD ensemble after dimension reduction via time-lagged independent component analysis (TICA) (*46*); each dot represents a structure snapshot, characterized by the most relevant independent components IC1 and IC2, which describe collective motions. In Fig. 3A, each of the obtained seven Markov states (colors and red numbers) represents a particular conformation of the helix, characterized in Fig. 3B by the number of intra-helical hydrogen-bonds (colors) ranging from 10 (fully folded) to zero (unfolded). The conformations corresponding to these seven Markov states (top of Fig. 3C, hydrogen-bonds as red dots) also show this sequence of decreasing helical structure. Our Markov state analysis yields free energy estimates for each state (bottom of Fig. 3C, thick bars) and transition rates between these states, ranging from 0.2 to 24.1 µs (arrows). These form a ‘folding funnel’ very similar to the one characteristic for folded proteins (*7*). Resolved by residue, the whole spectrum of timescales of the p53 dynamics (Fig. 3D, blue bars) densely covers a range between a few and several hundred ns.

**Fig. 3.**
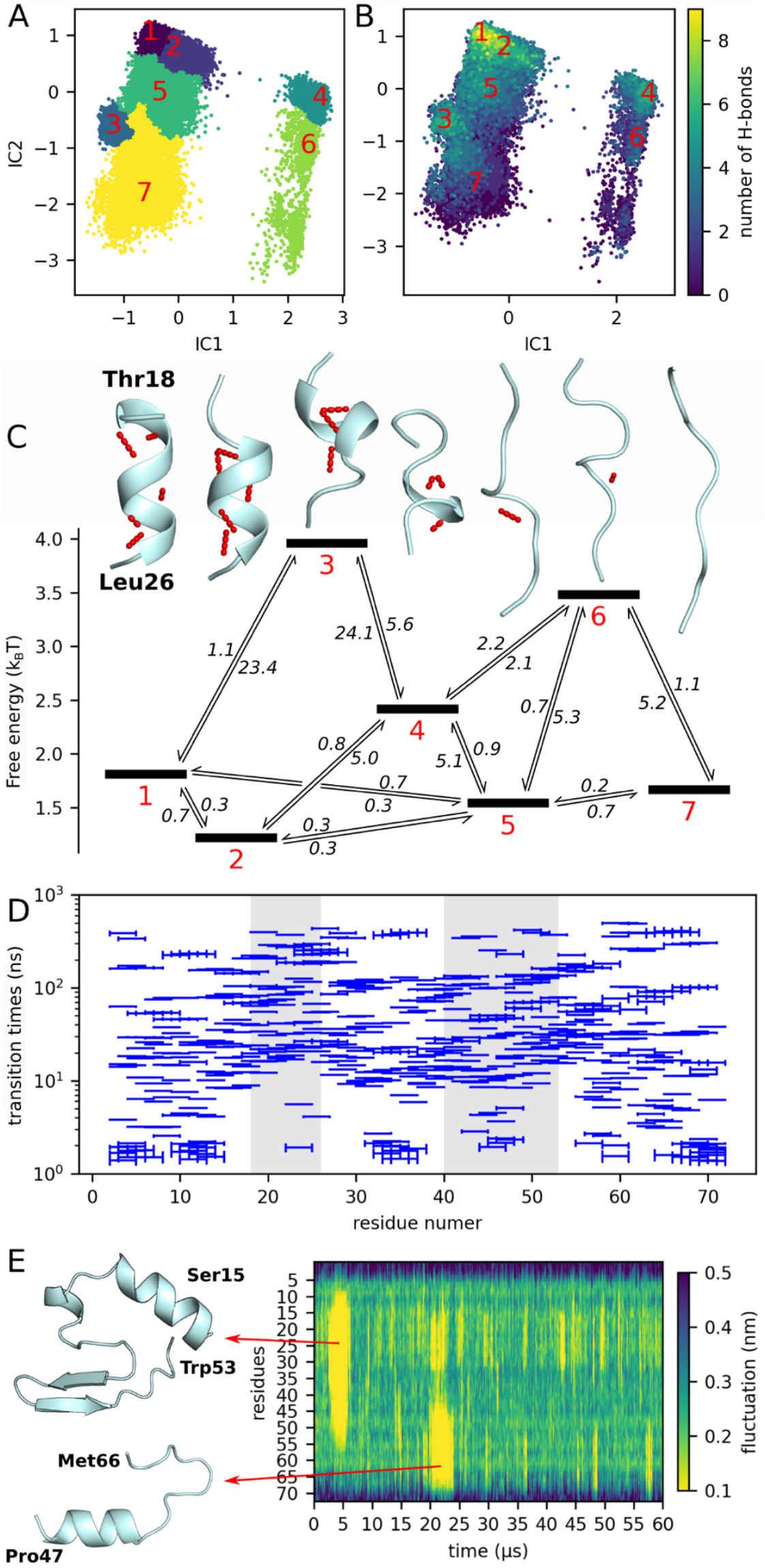
Markov state model reveals multi-timescale folding/unfolding dynamics of helix 1 underlying tier 1 motions of p53-TAD. (**A**) Projection of the conformational ensemble of helix 1 (residues 18-26) onto the TICA space defined by the two collective coordinates IC 1 and IC 2 that contribute most to the intra-helical dynamics using WT trajectories. Colors and red numbers indicate seven Markov states. (**B**) The same projection as in (A), but colored by the number of intra-helical hydrogen bonds. (**C**) Representative structures of the seven Markov states (intra-helical hydrogen bonds shown as red dots, top) and their free energies (black bars, below); mean first passage times of transitions between the states (arrows) are shown in μs. For a similar analysis (A to C) of helix 2, see fig. S16. (**D**) Per-residue timescale spectra (blue bars) calculated from MD trajectories, obtained using four residue (one helix turn) segments across the p53-TAD sequence (statistical uncertainties shown as error bars). Gray shaded areas indicate helix 1 and helix 2 residues. (**E**) Example of C_α_-C_α_ distance fluctuations analysis. The trajectory visited multiple transient tertiary structures from which two are shown. For every residue, distance fluctuations, ranging from 0.1 (yellow, rigid) to 0.5 nm (dark blue, flexible), were determined and averaged over a 0.1 μs sliding time window.

Taken together, the IDP undergoes structural dynamics with transient and stepwise secondary structure formation that are far more complex than a simple two-state model. This finding also explains why a single two-state RD profile (*44*) is unable to properly describe the E-CPMG RD profiles calculated from our MD simulations; instead, a stretched CMPG model that represents a superposition of many CPMG curves at different frequencies (*43*) (indicated by the blue dashed lines in Fig. 1B), fits the calculated profiles very well (Fig. 2 and figs. S4 and S5). Notably, the stretching factor of γ= 0.6 obtained from this fit also yields a timescale spectrum covering a similar range (fig. S10).

### Tier 0: Multi-µs dynamics due to transient formation of tertiary structure elements resembling folding intermediates of natively folded proteins

Which structural motions give rise to the slow 215 µs (tier 0) kinetics (cf. Fig. 1B left side of the blue curve)? To answer this question, we systematically searched for long-lived stable structures in our MD trajectories. Indeed, a residue-resolved analysis of C_α_-C_α_ distance fluctuations (Fig. 3E) revealed sporadic multi-µs collective excursions of pronounced stability, which are both larger and longer lived than the transient helices (Fig. 2D) of tier 1. We identified a total of 18 such long-lived events in the 0.6 ms WT simulations and 75 for P27A, three of which are shown in the summary Fig. 4 (right side). They represent metastable tertiary structure elements larger than simple α-helices, which persist up to 5 μs. For the P27A mutant, the simulations suggest that the removal of the sterically restrictive proline increases the conformational flexibility of p53-TAD, which enables the formation of larger and longer-lived tertiary structures. The fact that over four times more long-lived structure formation events are seen in the P27A simulations explains the much more pronounced low-frequency signal seen in the P27A CPMG RD profiles.

**Fig. 4.**
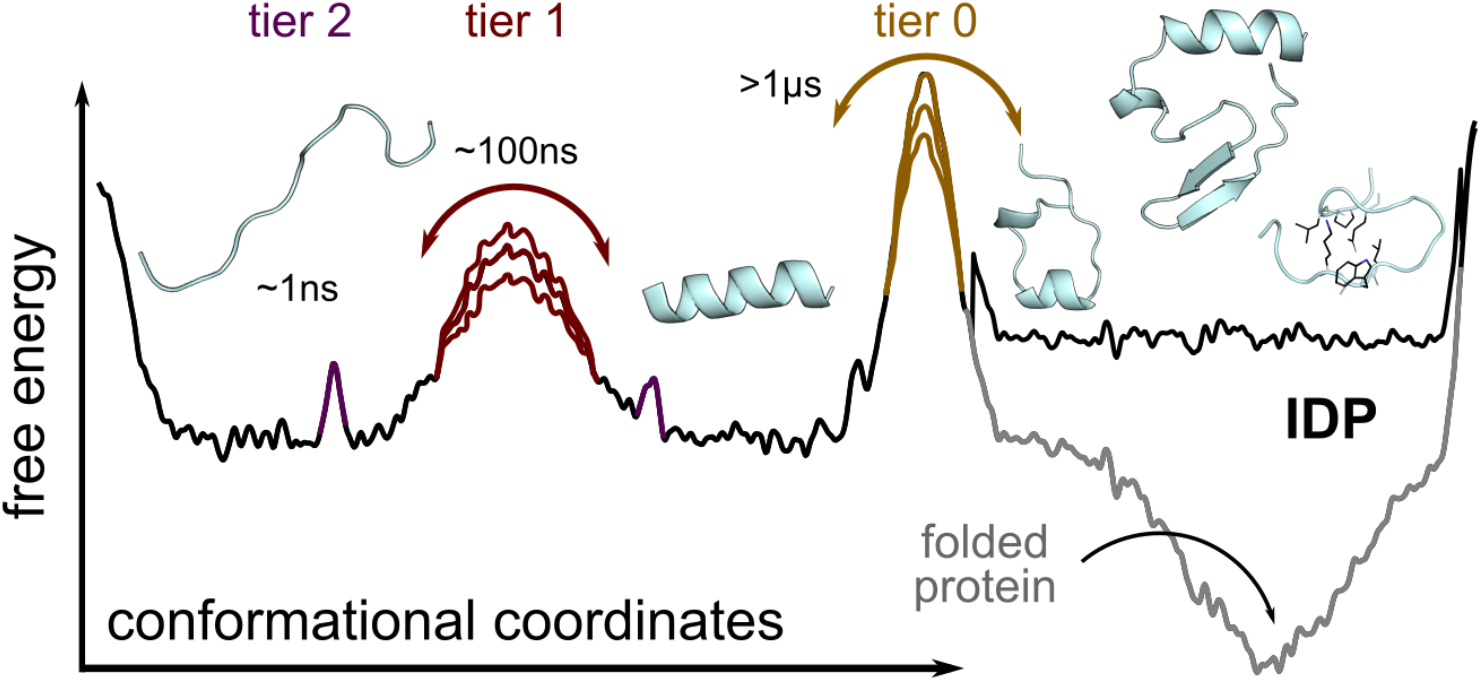
Sketch of the p53-TAD IDP hierarchical free energy landscape emerging from our combined NMR/MD study. Schematic free energy landscape (black curve) summarizing the three distinct tiers of p53-TAD dynamics emerging from our combined NMR/MD study. A hierarchy of free energy barriers (colored) similar to that known for folded proteins (*5*) governs fast (ns) reorientation and reconfiguration dynamics (tier 2 in purple), slower (sub-µs) supra-τ_c_ dynamics arising from transient helix formation (tier 1 in red), and very slow (multi-µs to ms) formation of transient tertiary structure elements (tier 0 in beige). The indicated variation of barrier heights is also similar to that observed for folded proteins and gives rise to a correspondingly broadened spectrum of relaxation times, which explains the observed stretched RD profiles. Contrary to folded proteins (gray curve), the transient tertiary structure elements of IDPs (tier 0) are only metastable and difficult to access experimentally. Structures shown as ribbon plots within the respective energy wells indicate typical intermediates seen in the MD ensemble.

Some of these structures are also stabilized through hydrophobic contacts (rightmost example in Fig. 4); apparently, hydrophobic collapses contribute to tier 0 dynamics, rather than to the faster tier 1. Notably, these long-lived tertiary structures frequently form in the presence of (and involve) one of the two much shorter-lived α-helices (e.g., the largest structure shown in Fig. 4), thereby stabilizing these helices over much longer timescales. This observation suggests that, at least for this particular IDP, the helices form “nucleation seeds” promoting subsequent tertiary structure formation attempts, similar to primary folding intermediates seen for natively folded proteins (*47, 48*).

### Tier 2: A stretched spectral density function reflects complex multistate dynamics also at fast nanosecond timescales

Contributing to the NMR RD profiles are also faster, ns and ps motions, which have been probed by regular relaxation experiments (*20*). These motions are characterized by the spectral density function (SDF) *J*(*v*), which is the Fourier transform of the correlation function of the dipolar couplings and the chemical shift anisotropies (*49*). Despite its central role, the full *J*(*v*) is inaccessible to experiments; rather, combinations of multiple NMR measurements such as *η*_xy_, τ_c_, *R*_1_, *R*_2_, and NOE at different conditions only allows to determine isolated points of the *J*(*v*) at specific frequencies (*9*).

From our validated MD trajectories, we calculated the full *J*(*v*) of p53-TAD (see SM (M)). Direct comparison with the above five NMR observables derived from multiple NMR measurements shows good agreement (figs. S6, S7, S8), facilitating structural interpretation of the parts of the *J*(*v*) that are inaccessible to experiment. Notably, a simple *J*(*v*) assuming a single state model, calculated from the measured τ_c_ values (fig. S7, gray lines) generally deviate more from the measured *J*(*v*) points at *v* = 96.3 MHz than the *J*(*v*) calculated from our simulations, providing independent evidence for the presence of multi-state dynamics involving a wide spectrum of timescales also within this fast tier 2 regime.

Because the orientational dynamics within secondary structure elements are expected to be slower than those of disordered regions, residues that are involved in the formation of transient helices should show correspondingly slower correlation times. Indeed, whereas fast sub-ns dynamics are seen for the terminal regions, the local reorientation dynamics of residues with substantial helical population slow down to 1 to 2 ns (fig. S6). These timescales agree well with previous NMR measurements (*11*) as well as with single-molecule FRET experiments on the Csp*Tm* cold-shock protein (*15*). Two regions show particularly slow dynamics and coincide with helix 1 and helix 2 (gray shaded Fig. 2, fig. S6), underscoring the slower rigid body motion of these transient helices within the faster rearranging disordered phase.

Our simulations also provide information on the fast (tier 2) conformational dynamics of p53 and enables comparison to polymer models (*17, 42, 50*). In particular, p53 is highly and rather homogeneously flexible with a persistence length of 2-3 amino acids (fig. S17A,B). Its scaling exponent of *v* = 0.66 is compatible with a self-avoiding chain model (*51*), which agrees with previous measurements of polymers in good solvents (*52*) and is above the critical Θ-point, at which chain-chain and chain-solvent interaction balance at the thermodynamic phase boundary (*53*).

Exponential fits to the orientation autocorrelation function (fig. S17D) show even higher flexibilities at short contour lengths, in line with single-molecule FRET experiments (*15*). An unexpected additional component is seen with a considerably longer persistence length of over 5 nm, pointing to so far unresolved partial long-range order, likely due to self-crowding governed by the slower tier 1 and tier 0 structural dynamics. These findings are supported by similar fast components of the time-lagged structure RMSD decay as well as time autocorrelation functions of the radius of gyration and end-to-end distance (10 ns, 22 ns, and 18.5 ns, respectively, figs. S17E,F). The latter two also reveal additional, much slower dynamics of 405 ns and 350 ns, respectively, likely arising from tier 1 dynamics. Overall, the structural dynamics of p53 resemble those of a highly flexible heteropolymer with fast reorganization dynamics (*14*) at short contour lengths and additional slower dynamics at larger length scales that give rise to pronounced long range correlations.

## Discussion

High-power RD NMR measurements (E-CPMG) (*36*) performed at 1.2 GHz combined with milliseconds atomistic MD simulations revealed a previously unappreciated hierarchy of complex structural dynamics of the intrinsically disordered transactivation domain of the central gene regulation protein p53 (p53-TAD). Covering a dynamic range of over seven orders of magnitude, from ps to sub-ms, we found three tiers of quite diverse structural dynamics (Fig. 4). This finding follows independently and consistently from our NMR measurements as well as from our MD simulations and challenges the prevailing view that the lack of stable folded structures of IDPs implies an unstructured and shallow free energy landscape that gives rise only to rather simple and fast tier 2 reorganization dynamics (*14*).

Rather, our findings suggest that the classic Frauenfelder stretched exponential description of folded protein dynamics, which arises from an underlying multi-tier hierarchical free energy landscape (*5*), also applies to the largely unstructured p53 IDP. Notably, we have obtained similar results for Measles Virus NTAIL (fig. S20), an unrelated IDP, which suggests that these unexpectedly complex dynamics may be a general feature of many other IDPs.

In particular, transient secondary structure formation dynamics are seen on a broad range of supra-τ_c_ (10s to 100s of ns) timescales between several ns and sub-µs, forming a second dynamics level, tier 1. In our simulations, these serve as dynamic precursors for the tier 0 formation of metastable and quite diverse tertiary structures at different positions along the TAD sequence. We speculate that these may be a key factor in the ability of p53 to fold into a broad range of different structures upon binding to a similarly broad range of different partners. Indeed, a structure similarity search in the Protein Data Bank (PDB) (*54*) revealed that motifs of these tertiary structures seen in our MD simulations also occur in proteins (SM (Z)). Because such rich conformational selection dynamics has the potential to accelerate binding, this third dynamics tier, emerging from an unexpectedly complex free energy landscape (Fig. 4), may thus be not only a feature of p53-TAD, but also a key to understanding the binding promiscuity of many other IDPs in general.

Our results show that the structural dynamics of IDPs are not limited to fast and essentially random reorganizations, but are organized across a broad hierarchy of timescales, similar to the conformational dynamics of natively folded proteins. This complexity also mirrors the (un-)folding dynamics of folded proteins, where both MD simulations (*55*) and Φ-value measurements (*56*) have revealed the presence of transiently stable secondary and tertiary structure elements along unfolding pathways, similar to those seen here for p53-TAD in equilibrium. Such transient structures are not merely biochemical curiosities, but may be relevant for binding to interaction partners and as potential drug targets. By shifting the balance of conformational ensembles that IDPs explore, such interventions could selectively modulate their activity.

## Acknowledgments

We thank Claudia Schwiegk for protein sample production, Lars Bock, Eliane Briand, Nicolai Kozlowski, Bert de Groot, and Florian Leidner for discussions and suggestions, and Petra Kellers for help editing the manuscript.

## Author contributions

Conceptualization: D.S., S.R., H.G., C.G., S.P.

Formal analysis: D.S., S.R., H.G., S.P., D.M.

Investigation: D.S., G.N., S.R., R.K., S.P., D.M., G.J.R., N.E., A.K.R., D.L., S.B.

Methodology: D.S., S.R., H.G., S.P., D.M., G.J.R., A.K.R., D.L., S.B.

Project administration: H.G., C.G. Resources: H.G., C.G., S.B. Software: D.S.

Supervision: H.G, C.G.

Writing – original draft: D.S., H.G.

Writing – review & editing: D.S., S.R., H.G., C.G., D.M., S.P.

## Data and materials availability

The program code used for all calculations, additional sample calculations and sample data, as well as Movies S1, S2, and S3, have been deposited at https://github.com/dszollosi/p53_TAD_dynamics. All NMR spectra have been deposited at https://edmond.mpg.de/previewurl.xhtml?token=bbf1f9d4-4eb2-455e-921f-41e767868656, DOI:10.17617/3.JWVPWJ. All molecular dynamics simulation trajectories are available from H.G. upon reasonable request; these data have not been deposited due to large data size (>1TByte).

## Supplementary Materials

The PDF file includes:

Materials, Methods, Supplementary Results

Figs. S1 to S21

Tables S1 to S3

References (57–133)

Movies S1 to S3

