## Supplemental Material for "Hierarchically ordered multi-timescale structural dynamics of the intrinsically disordered p53 transactivation domain"

References in Supplementary Materials: #57 - #133

**Note:** For the convenience of the reader, the information provided here is arranged according to the sequence referred to in the main text. As a result, Methods and Results are interlaced and marked appropriately in the respective headings.

**Contents**

(A) Methods: Sample preparation ... 3
(B) Methods: Sample conditions ... 3
(C) Methods: NMR relaxation measurements ... 4
(D) Methods: Molecular dynamics simulations ... 6
(E) Results: NMR RD profiles ... 6
(F) Results: Difference between relaxation from exchange versus relaxation due to molecular tumbling ... 10
(G) Results: Comparison of NMR RD profiles at 1.2 GHz versus 950 MHz ... 11
(I) Methods: (a) Exchange related relaxation: $R_{2,ex}$ ... 13
(J) Methods: (b) Tumbling related relaxation: $R_{2,0}$ ... 20
(K) Methods: Calculation of SDFs and derived NMR observables $\eta_{xy}$, $\tau_c$, $R_1$, $R_2$, and NOEs ... 20
(L) Results: Comparison of measured with calculated NMR observables $\eta_{xy}$, $\tau_c$, $R_1$, $R_2$, and NOEs ... 22

### (A) Methods: Sample preparation

Recombinant p53-TAD (1-73) was expressed as a fusion protein with an N-terminal $Z_2$ domain using a modified pET28a vector (*57*). Perdeuterated $^{15}N$-labeled p53-TAD (1-73) samples were expressed at 25 °C in *E. coli* adapted to 100% $D_2O$ minimal medium supplemented with $D_7$-glucose as the carbon source and $^{15}N$-$NH_4Cl$ as the nitrogen source. Protein expression was induced with 1 mM IPTG. The expression culture was harvested 12 hours after induction. Recombinant p53-TAD was purified using immobilized metal affinity chromatography on Ni-NTA resin (Macherey-Nagel, Germany) followed by tabac mosaic virus (TEV) protease cleavage at room temperature. The cleaved protein was reloaded onto Ni-NTA resin to remove the $Z_2$ domain and TEV protease. Gel filtration on a Superdex 75 16/60 HiLoad column (GE Healthcare) was performed to further purify the p53-TAD (1-73) fragment. Due to cloning the WT fragment included an additional N-terminal Gly-Ser-extension and the P27A mutant a Gly-Ser-His-Met-extension. The fractions containing the purified protein were combined, concentrated with a 10 MWCO concentrator (Vivascience).

### (B) Methods: Sample conditions

Nuclear Magnetic Resonance (NMR) experiments were performed on a 1.5 mM p53-TAD sample in 50 mM sodium acetate buffer at pH 6.3, containing 50 mM sodium chloride and 0.03% sodium azide. Backbone amide $^1H$ Off-resonance relaxation dispersion ($R_{1\rho}$-RD) experiments at temperatures ranging from 262 K to 265 K were carried out in glass capillary tubes to produce super-cooled conditions below the freezing point of water. Each capillary of 1 mm outer diameter (Wilmad, Buena, New Jersey) contained 25 µl of the p53-TAD sample, and 12 such capillaries were placed inside a 5 mm NMR sample tube. The sample was purposefully not labeled with $^{13}C$ nuclei to avoid the necessity of $^{13}C$ decoupling, which could be an extra source for RF heating. In addition, the heteronuclear J coupling of $C_\alpha$ and carbonyl carbon to the nearby amide proton and nitrogen nuclei can be a source of artifacts in relaxation dispersion profiles.

All NMR sub-$\tau_c$ relaxation measurements and $^1H_N$ extreme power Carr-Purcell-Meiboom-Gill (CPMG) RD measurements were performed on uniformly perdeuterated, $^{15}N$ labeled proteins (both p53-TAD WT and P27A mutant) back exchanged with 100% $H_2O$. The samples were finally buffer exchanged to 50 mM sodium acetate buffer at pH 6.3 containing 50 mM NaCl, 5% (vol/vol) $D_2O$, and 0.02% sodium azide. The final p53-TAD protein concentrations of WT and P27A samples were 1.0 mM and 0.7 mM, respectively. Each sample was transferred to a 2 mm capillary and was placed within the magnet using a Bruker Match insert assembly.

**(C) Methods: NMR relaxation measurements**

All NMR spectra were collected with a Bruker Avance III HD spectrometer operating at 950 MHz equipped with a TCI 5 mm cryo-probe and a Bruker Neo spectrometer operating at 1.2 GHz $^{1}$H field strengths, equipped with a TCI 3 mm cryo-probe. Sample temperature was controlled with dry $N_2$ gas using Bruker BCU-II VT units with medium chiller strength and 670 liter/hour gas flow for all experiments. Temperatures over the 263–298 K range were calibrated using a 3 mm Greisinger GMH 3750 thermometer equipped with a thermocouple. All spectra were referenced with respect to the water peak.

2D-$^{15}$N-$^{1}$H spectra, collected with a FAST-HSQC (*58*) pulse sequence at each temperature, were used to validate sample conditions. Assignments were transferred from previously published sources (*59*) at 298 K and propagated to spectra collected at 277 K and 263 K by recording a series of 2D-HSQC spectra at 5 K temperature intervals. All 2D-NMR data were processed within the UNIX software environment NMRPipe (*60*) and were further analyzed and visualized using the software package nmrfam-sparky (*61*).

The $^{15}$N $R_1$, $R_{1\rho}$ experiments were recorded at 298 K, 277 K, and 263 K under 950 MHz $^{1}$H field strength using standard protocols (*62*) and an eight-point measurement scheme (with two repeat points). $^{15}$N-$R_{1\rho}$ relaxation rates were measured using a spin lock field of 2 kHz and were subsequently converted to $R_2$ values following the standard methodology (*63*). Heteronuclear $^{15}$N nuclear Overhauser effect (NOE) measurements were recorded with 5 s mixing time using standard protocols (*62*) at 298 K and 277 K under 950 MHz $^{1}$H field strength.

Site-specific $^{15}$N-transverse cross-correlated relaxation (CCR) rates ($\eta_{xy}$) were measured via 2D $^{15}$N TRACT (TROSY for rotational correlation times) (*64, 65*) experiments performed at 298 K, 277 K, and 263 K under 950 MHz and 1.2 GHz field strengths. The software NMRPipe was used to process all pseudo-3D spectra as well as to extract relaxation rates. Uncertainties in measured relaxation rates were estimated using error propagation from spectral RMS noise. Measured $\eta_{xy}$ values were converted (*66, 67*) to approximate site-specific rotational correlation times ($\tau_c$) and further converted to approximate chemical exchange free intrinsic $^{15}$N and $^{1}$H transverse auto-relaxation rates ($R_{2,0}$) as published (*68*) using Python scripts. For all calculations, standard values (*66*) for $^{15}$N-$^{1}$H bond length 1.02 Å, $\theta_{xy}$ = 17°, $^{15}$N CSA = -160 ppm, and $^{1}H_N$ CSA = 10 ppm (*69*) were used. The effect of varying these parameters in the calculation of $\tau_c$ has been described in detail elsewhere (*66*). The relaxation data was visualized using OriginPRO software.

One set of $^{1}H_N$ Carr–Purcell–Meiboom–Gill (CPMG) experiments using extreme power were recorded at 263 K and 298 K with the published pseudo-4D IP/AP scheme pulse sequence (*70*) with modifications as published in (*36*), required to run the experiment at extreme CPMG frequencies (*71*). The experiments (3, *72, 71, 73, 74, 75*) were performed with the power on the $^1$H channel set to 18 W ($^1$H 90° pulse length ~ 8.1 μs) on the 950 MHz spectrometer and ~ 16.3 W ($^1$H 90° pulse length ~ 6.25 μs) on the 1.2 GHz spectrometer. A recycle delay of 3 s for all experiments was used to ensure minimal sample heating and a low-duty cycle. All experiments were recorded with 128 initial equilibration scans to equilibrate the spin system and sample temperature before data acquisition. For the 1.2 GHz experiments, the constant CPMG duration ($T_{CP}$) was set to 20 or 40 ms, and 28 points were sampled in the CPMG frequency dimension (including the reference plane and two repeat points) ranging from 100 Hz up to 40 kHz. For each experiment, 100–120 (indirect dimension) and 1536 (direct dimension) complex points were recorded with 16 scans. The experiments were recorded with a 2 ppm bandwidth E-BURP refocusing pulse (centered at the middle of the $^1H_N$ region ~ 8.0 ppm) at the center of the CPMG duration. The total experiment time was ~ 90 h at each temperature for each sample. The experiments at 950 MHz were acquired with 30 points (including the reference plane and two repeat points) in the CPMG frequencies dimension, ranging from 100 Hz up to 30.7 kHz, and with a hard pulse at the center of the CPMG duration. For each experiment, 120 (indirect dimension) and 1024 (direct dimension) complex points were recorded with 4 scans, totaling ~ 24 h experimental time per sample at each temperature.

For all experiments, spectra were processed, and relaxation rates were extracted separately for the IP and AP sets of spectra using the UNIX software environment NMRPipe, followed by averaging for subsequent analysis. The differences between values from the IP and AP datasets were minimal. Site-specific solvent exchange contributions to measured $^1H_N$ $R_2$ values at 298 K were estimated at 950 MHz, using differences of site-specific $^1$H $R_1$ values acquired from two sets of inversion recovery pulse sequences recorded with standard parameters and recycle delays of > 10 $s^{-1}$. In the first experiment, water magnetization was kept along the Z axis, whereas in the second experiment, it was completely dephased with a low gradient (*76*).

The 2D [$^1$H-$^1$H]-NOESY experiment was collected on a Bruker Avance III HD spectrometer operating at 900 MHz equipped with a TCI 5 mm cryo-probe at 298 K. The NOESY experiment was performed with 120 ms mixing time and 1024 and 512 complex points along $t_1$ and $t_2$ dimensions, respectively. The NOESY data were processed using NMRPipe (*60*) and analyzed with nmrDraw and CARA (*77*).

#### (D) Methods: Molecular dynamics simulations

All simulations were performed using the molecular dynamics (MD) simulation software package GROMACS Version 2019.3 (*78*). Starting structures for each of the 30 trajectories were generated by first collapsing a fully extended P53-TAD (1-73) molecule performing a short generalized Born implicit solvent (GBSA) simulation (*79*). From these trajectories structures were selected which were not fully collapsed (radius of gyration > 3.5 nm) and did not contain any secondary structure elements. The p53-TAD (residues 1-73) WT protein and the P27A mutant were placed within dodecahedral boxes with an initial volume of 1222 $nm^3$ (edge lengths: 12.0, 12.0, 8.485 nm) solvated in water and 100 mM NaCl. Virtual sites were used to allow for a time step of 4 fs (*80*). The LINCS algorithm (*81*), applying a sixth-order iterative restraint on the bond distances, was used. The Particle Mesh Ewald (PME) algorithm (*82*) was used for electrostatic interactions with a cut-off of 1.0 nm. A reciprocal grid of 96 x 96 x 96 cells was used with fourth order B-spline interpolation. A single cut-off of 1.215 nm was used for the Van der Waals interactions. Neighbor searching was performed every 60 steps. Temperature was controlled by the velocity-rescale algorithm (*83*) with a 298 K target temperature and a 0.1 ps coupling constant; for pressure coupling the Parrinello-Rahman algorithm (*84*) was used with 1 bar target pressure and a coupling constant of 20.0 ps. All fully hydrated systems were allowed to equilibrate for 10 ns before the final production runs were started. Protein coordinates were recorded every 100 ps. All WT simulations were 20 µs long and all P27A simulations were 60 µs long to achieve better convergence for the expected slower P27A timescales. The Amber99sbws (*85*) force field was used in combination with the TIP4P2005s (*86*) water model.

#### (E) Results: NMR RD profiles

Figure S1 shows NMR RD profiles measured at 263 K and 1.2 GHz for WT (green dots) and P27A mutant (orange dots) of p53-TAD. The gray lines show fits of the CPMG equation (Eq. 17) to these measured data points using a Bayesian approach as described below (subsection ‘Fitting Procedure’). The Bayesian posterior was estimated using standard Monte Carlo sampling and provides an estimated uncertainty of the fit. 4000 samples were taken for the posterior distribution of the fitted parameters and used to calculate plausible CPMG profiles. These sample profiles were added to the figure as transparent lines, which appear as a shaded area around the mean fitted curve (darker opaque line). Relaxation times $\tau$ were obtained from these fits and are shown in table S1, together with their uncertainties also estimated from the posterior.

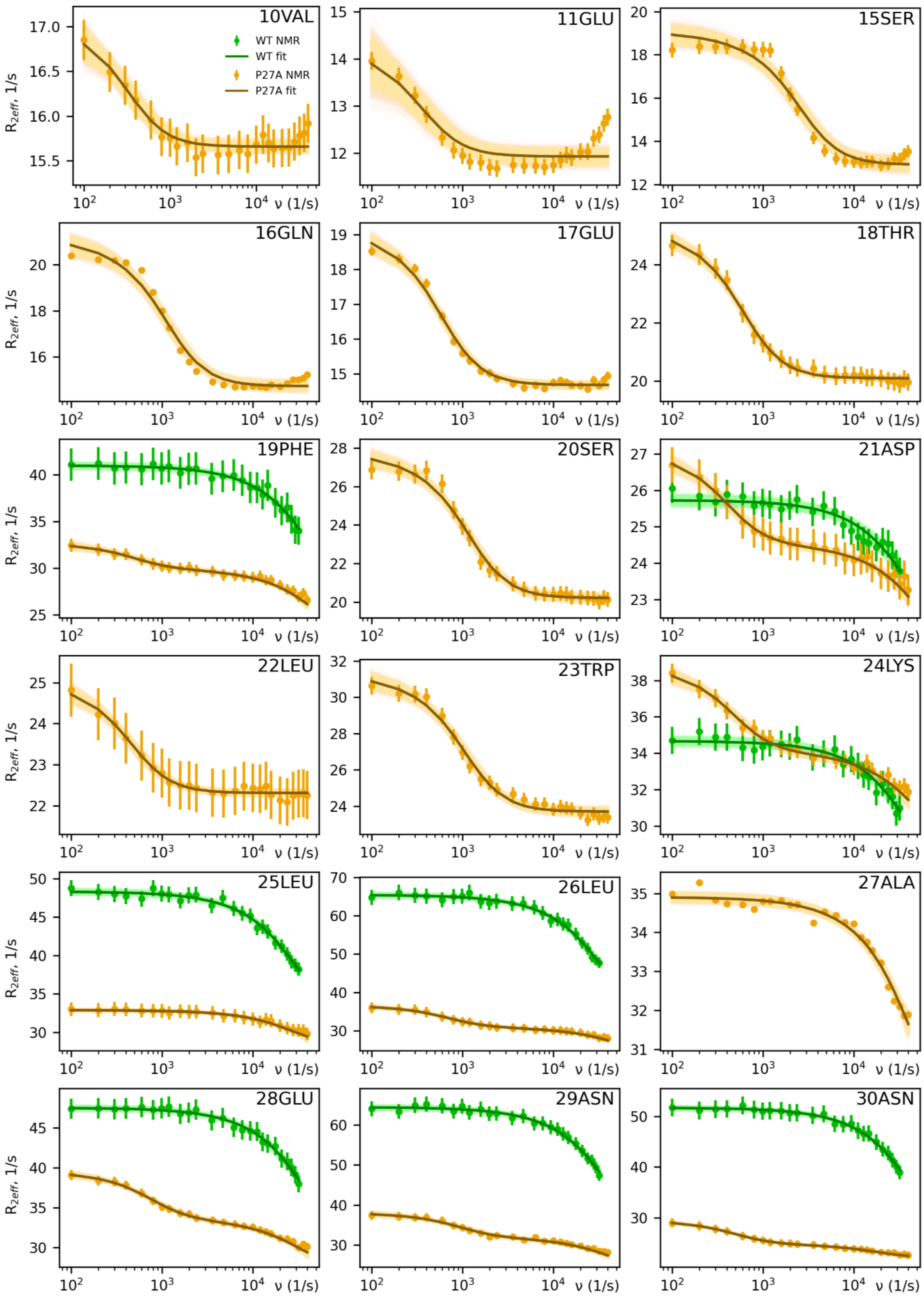
10VAL
11GLU
15SER
16GLN
17GLU
18THR
19PHE
20SER
21ASP
22LEU
23TRP
24LYS
25LEU
26LEU
27ALA
28GLU
29ASN
30ASN
WT NMR
WT fit
P27A NMR
P27A fit
$R_{2eff}$, 1/s
ν (1/s)

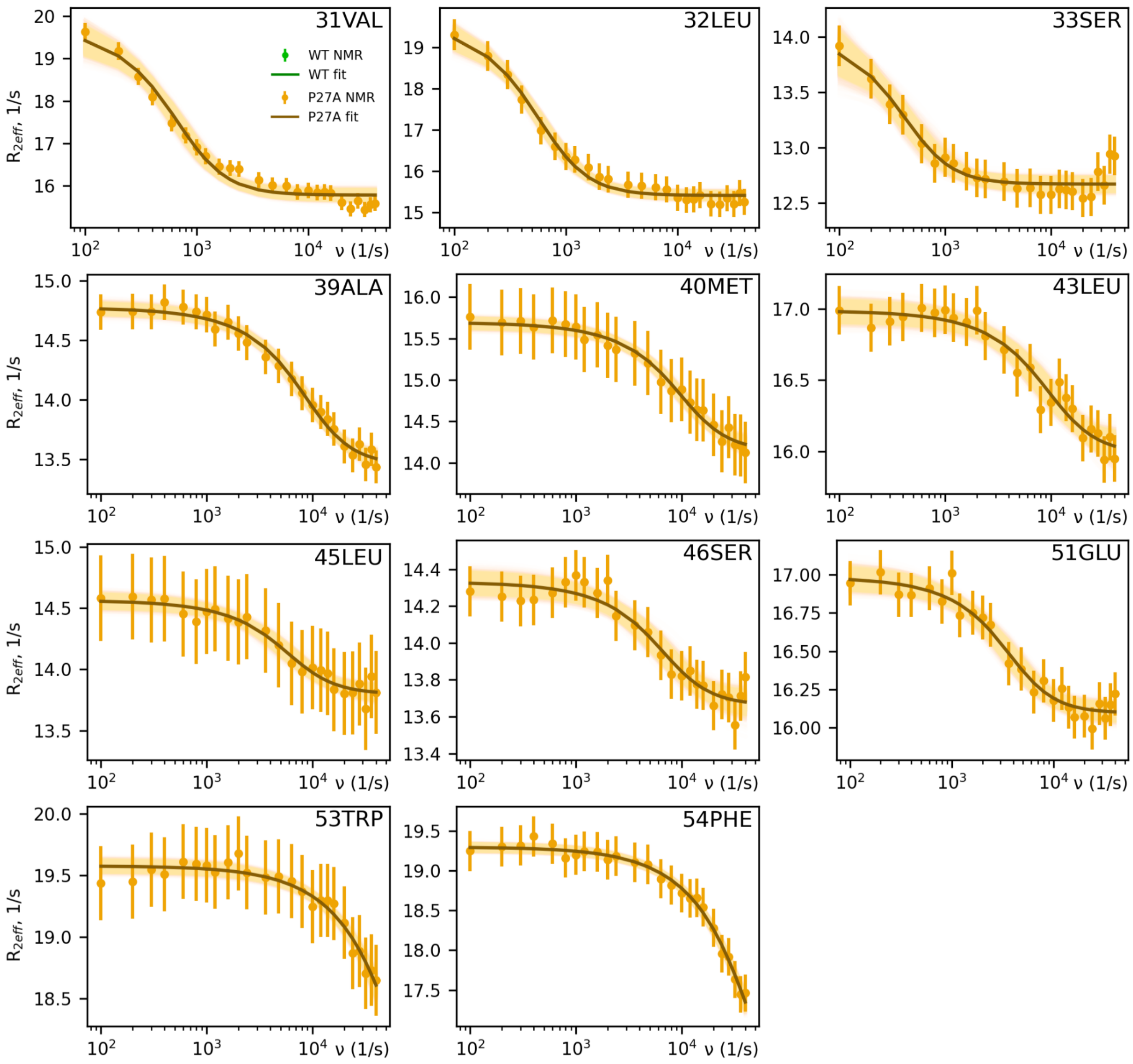

**Fig. S1. Complete set of p53-TAD NMR RD profiles for all measured residues**. Profiles were recorded at 263 K and at 1.2 GHz for WT (green) and for P27A (orange); vertical bars indicate measurement uncertainties. Superimposed are CPMG fits to the measured data points (gray lines) and uncertainties of the fits (shaded areas). Relaxation times $\tau$ obtained from these fits are listed in table S1.

| Residue | WT $\tau$ (μs) | P27A $\tau$ (μs) | P27A $\tau_2$ (μs) |
|---|---|---|---|
| Val10 | | 471.02 ± 70.65 | |
| Glu11 | | 435.06 ± 102.38 | |
| Ser15 | | 60.87 ± 5.77 | |
| Gln16 | | 136.03 ± 8.79 | |
| Glu17 | | 258.09 ± 13.37 | |
| Thr18 | | 249.52 ± 10.42 | |
| Phe19 | 3.92 ± 0.75 | 314.03 ± 45.23 | 3.22 ± 0.73 |
| Ser20 | | 130.34 ± 7.07 | |
| Asp21 | 4.06 ± 1.93 | 379.3 ± 30.99 | 2.74 ± 0.97 |
| Leu22 | | 332.77 ± 25.94 | |
| Trp23 | | 144.22 ± 9.39 | |
| Lys24 | 4.08 ± 1.13 | 320 ± 30.68 | 3.36 ± 1.06 |
| Leu25 | 5.65 ± 0.83 | 5.76 ± 1.34 | |
| Leu26 | 4.97 ± 0.64 | 220.37 ± 13.34 | 3.18 ± 0.76 |
| Ala27 | | 3.86 ± 0.79 | |
| Glu28 | 4.38 ± 0.73 | 202.48 ± 11.9 | 4.97 ± 1.29 |
| Asn29 | 4.62 ± 0.64 | 163.37 ± 13.91 | 3.74 ± 0.96 |
| Asn30 | 3.98 ± 0.52 | 310.69 ± 12.62 | 8.41 ± 1.05 |
| Val31 | | 228.47 ± 24.1 | |
| Leu32 | | 264.88 ± 20.84 | |
| Ser33 | | 367.26 ± 62.89 | |
| Ala39 | | 17.75 ± 1.46 | |
| Met40 | | 15.64 ± 1.61 | |
| Leu43 | | 16.56 ± 3.21 | |
| Leu45 | | 26.24 ± 3.71 | |
| Ser46 | | 23.38 ± 3.84 | |
| Glu51 | | 42.09 ± 5.47 | |
| Trp53 | | 2.28 ± 0.8 | |
| Phe54 | | 3.4 ± 0.79 | |

**Table S1. Lifetimes $\tau$ obtained from the CPMG fits to the measured NMR RD profiles shown in fig. S1.** P27A shows additional, drastically increased relaxation times. For certain residues at helix 1, a second timescale was detected in the same timescale range as for the WT. The amplitude of the faster relaxation process is close to the detection limit and, therefore, was not observed for most residues.

### (F) Results: Difference between relaxation from exchange versus relaxation due to molecular tumbling

CPMG measurements resolve exchange processes for relaxation frequencies slower than a few µs. To further probe the existence of tier 1 kinetics faster than these timescales, we also determined the fast individual rotational correlation times $\tau_c$, via TRACT (TROSY for rotational correlation times) measurements (crosses in Fig. 1G). This measurement allowed for an independent approach to detect dynamics faster than the timescale accessible to RD measurements, by comparing the relaxation amplitude at the highest available RD (CPMG) frequency with that observed for and explained by molecular tumbling. Because at each particular frequency the measured $R_{2,\mathrm{eff}}$ value contains contributions by all faster relaxation processes, the profiles shown in fig. S1 also contain information about these faster conformational dynamics, even though these are far outside the measurement frequency window between the correlation time and about 4 µs.

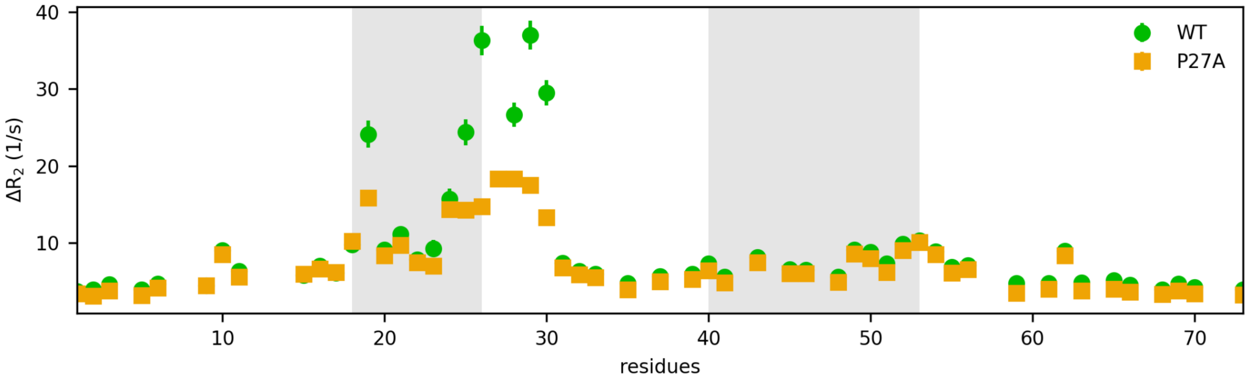

**Fig. S2. Fast supra-$\tau_c$ conformational p53-TAD dynamics.** Shown is the relaxation amplitude difference $\Delta R_2$ between the highest available CPMG frequency and that of relaxation explained by molecular tumbling, as measured via TRACT experiments. $\Delta R_2$ differences are shown for the WT (green dots) and the P27A mutant (orange squares). All measurements were performed at 263 K at a magnetic field strength of 1.2 GHz. Gray shaded areas indicate helix 1 (18-26) and helix 2 (40-54) residues.

Also contributing to the $R_{2,\mathrm{eff}}$ value are relaxation processes unrelated to conformational dynamics, however, in particular those arising from fast molecular tumbling. These are quantified by our TRACT measurements, and the difference $\Delta R_2$ between the $R_{2,\mathrm{eff}}$ value at fastest measured CPMG frequency and the TRACT relaxation value therefore measures the presence and amount of additional supra-$\tau_c$ dynamics faster than maximum E-CPMG frequency (fig. S2). To assess the relaxation contribution of fast exchange above the RD measurement frequencies, the relaxation contribution from dipolar coupling of $^{1}H$-$^{15}N$ interactions and proton chemical shift anisotropy were therefore subtracted from RD measurements (1.2 GHz). Indeed, residues within and next to helix 1 (residues 18-30) show markedly larger relaxation amplitudes, which must thus arise from dynamics slower than tumbling but faster than that captured at RD frequencies, establishing supra-$\tau_c$ tier 1 dynamics within this otherwise “blind spot” of NMR measurements. These dynamics are also unaffected by the mutation, which provides additional and independent evidence that the 215 μs tier 0 dynamics seen for P27A are a new process, but leaves the origin of tier 1 dynamics unresolved.

#### (G) Results: Comparison of NMR RD profiles at 1.2 GHz versus 950 MHz

The detection of the 4 μs motion in the P27A mutant was possible at 1.2 GHz only but not at 950 MHz (fig. S3). For such fast motions, the amplitude of the RD profile is very small indeed; however, because it increases quadratically with the field strength, the amplitude is 60% larger at 1.2 GHz compared to 950 MHz, thus enabling detection of these dynamics.

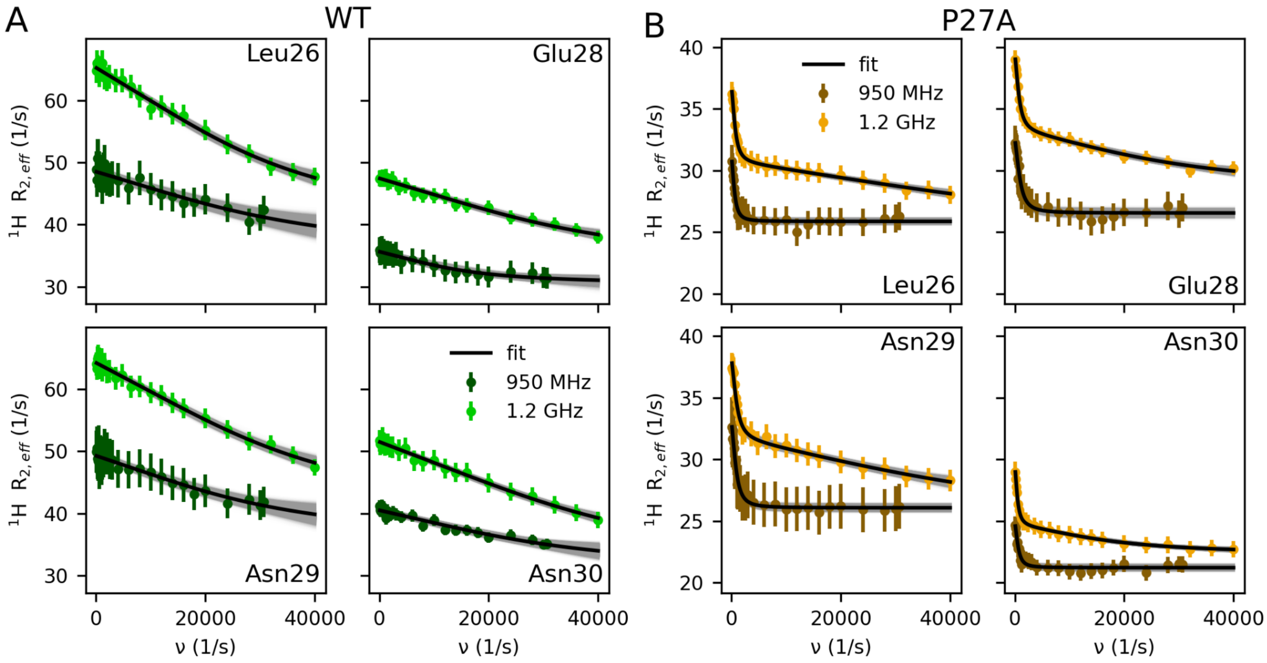

**Fig. S3. Field-dependent detection of fast dynamics in p53-TAD using high-power $^1H^N$ CPMG RD, highlighting the importance of 1.2 GHz $B_0$ field used in the study.** Comparison of RD profiles measured at 263 K and at two magnetic field strengths: 950 MHz (dark colors) and 1.2 GHz (light colors), for both WT (**A**, green/dark green dots with error bars) and P27A (**B**, orange/brown dots with error bars). RD profiles of WT and P27A at 950 MHz were fitted by a single CPMG equation (mean fit: black line, posterior distribution of the mean: shaded gray area); the RD profiles for P27A measured at 1.2 GHz were fitted by the sum of two CPMG functions, yielding two different timescales. Relaxation times $\tau$ obtained from these fits are listed in table S2.

| Residue | WT $\tau$ (µs) | | P27A $\tau$ slow (µs) | | P27A $\tau$ fast (µs) |
|---|---|---|---|---|---|
| | **950 MHz** | **1.2 GHz** | **950 MHz** | **1.2 GHz** | **1.2 GHz** |
| Leu26 | 5.02 ± 1.22 | 4.97 ± 0.64 | 262.23 ± 24.13 | 220.37 ± 13.34 | 3.18 ± 0.76 |
| Glu28 | 6.31 ± 2.52 | 4.38 ± 0.73 | 153.27 ± 18.61 | 202.48 ± 11.9 | 4.97 ± 1.29 |
| Asn29 | 5.62 ± 1.27 | 4.62 ± 0.64 | 171.99 ± 14.19 | 163.37 ± 13.91 | 3.74 ± 0.96 |
| Asn30 | 4.64 ± 1.13 | 3.98 ± 0.52 | 241.5 ± 36.12 | 310.69 ± 12.62 | 8.41 ± 1.05 |

**Table S2. Relaxation times ($\tau$) obtained from the fits to the NMR RD profiles.** Profiles of WT and P27A were measured at 263 K at 950 MHz and 1.2 GHz, as shown in fig. S3.

### (H) Results: Comparison of NMR and MD RD profiles

Figures S4 and S5 compare RD profiles derived from NMR measurements with those calculated from our MD simulations, summarized in Fig. 2B of the main text. Two major components contribute to the exchange contribution to relaxation, (a) relaxation due to exchange between states (e.g., folding and unfolding) with different chemical shifts $R_{2,ex}$; (b) relaxation caused by molecular tumbling and sub-$\tau_c$ internal motion summarized in $R_{2,0}$. The effective relaxation $R_{2,eff}$ is the sum of these two components $R_{2,ex}$ and $R_{2,0}$, which therefore were calculated separately from the trajectories as described below, figs. S4 and S5 show their sum.

As can be seen, for most residues quantitative agreement is achieved. In particular, the dynamic profile showing higher amplitudes for the helical regions is very similar for both measured and calculated amplitudes. Deviations are seen mainly for the N-terminal region up to residue Val10, likely caused by the presence of 2-4 additional residues in the experiments, which were required for cloning. These were omitted in the MD simulations, which aimed at a most accurate modelling of the WT p53-TAD (for validation, see SM (N) and fig. S9). The deviation seen for Asn28 is presumably due to the absence of proline isomers in the MD simulations, supported by the fact that much better agreement is seen for the P27A mutant (Fig. 2B, orange).

#### *Uncertainties and controls*

We note that the comparison (Fig. 2, B and E to H) of measured RD profiles with those calculated from MD simulations involves three sources of uncertainty. First, IDPs are known to be particularly sensitive to force field inaccuracies (*87-89*); we have therefore carried out test simulations using different force fields and obtained similar profiles for the NMR observables (fig. S18). The best agreement was seen for the Amber99sbws force field, which we therefore chose for further analysis. Second, despite extensive millisecond sampling, the MD ensemble likely does not fully cover the full structure ensemble probed in the experiments. Although most residues show converged RD profiles (fig. S19), those of residues Asp49, Phe54, Glu56, Glu62, Ala63, and Met66 indicate relaxation times similar to the simulation length. For these residues, therefore, relaxation times cannot be reliably determined and were omitted from Fig. 2B. Third, the accuracy of calculated RD profiles relies on that of our chemical shift calculations using SPARTA+ (*90*), which yielded very similar results as SHIFTX (*91*). Notably, because the orientational correlations times were directly calculated from the orientational autocorrelation of the backbone NH bond vector without recourse to chemical shifts, their comparison with the measured TRACT relaxation times also provides an independent test of the accuracy of the simulations. Indeed, for all residues except the N-terminal ones, very good agreement is seen (Fig. 2B). Here, too, this deviation is mainly due to the additional residues required for sample preparation, as evidenced by the markedly improved agreement obtained for additional control simulations which included these residues (fig. S9). Finally, deviations between experiment and MD simulations might arise from the very slow isomerization dynamics of prolines (*92*), which are not described by our simulations. However, additional control simulations comparing, e.g., WT cis-Pro8 with trans-Pro8 showed no significant differences (fig. S6).

**(I) Methods: (a) Exchange related relaxation: $R_{2,ex}$**

Conformational dynamics such as folding and unfolding of an intrinsically disordered protein (IDP) induce changes in the chemical shift of the involved nuclei and were probed by CPMG RD experiments as described above. By fitting to the CPMG curve (Eq. 18), the characteristic exchange timescale between the folded and unfolded states was determined for those residues for which the exchange rate falls within the detectable frequency range set by the maximum applicable RF power and where the chemical shift difference is sufficiently large. RD profiles were calculated from our MD simulations via the power spectrum of the combined chemical shift time traces $CS_{HN}(t)$, calculated for all available trajectories of the respective nuclei. Chemical shifts of the p53-TAD backbone protons were predicted using the software SPARTA+ (*90*) from simulation frames separated 100 ps in time from 20 µs trajectories for the WT and 60 µs trajectories for the P27A system. As shown by Xue et al. earlier, the RD profile is derived from the chemical shift autocorrelation functions (*93, 94*), which we have calculated from $CS_{HN}(t)$ via the properly normalized power spectrum, i.e., the absolute-squared Fourier transform

$$R_{2,ex}(\nu)\frac{1}{2}\mathrm{N}\Delta t = \left|\mathcal{F}\big(CS_{NH}(t)\big)\right|^2 \quad (1)$$

using the Wiener-Khinchin theorem (*95*). Here, $\nu$ is the CPMG frequency, *N* is the number of simulation frames, $\Delta t$ is the time step between frames (100 ps). Note that the unit of $CS_{HN}(t)$ is rad/s. RD profiles were calculated separately from each MD trajectory and then averaged.

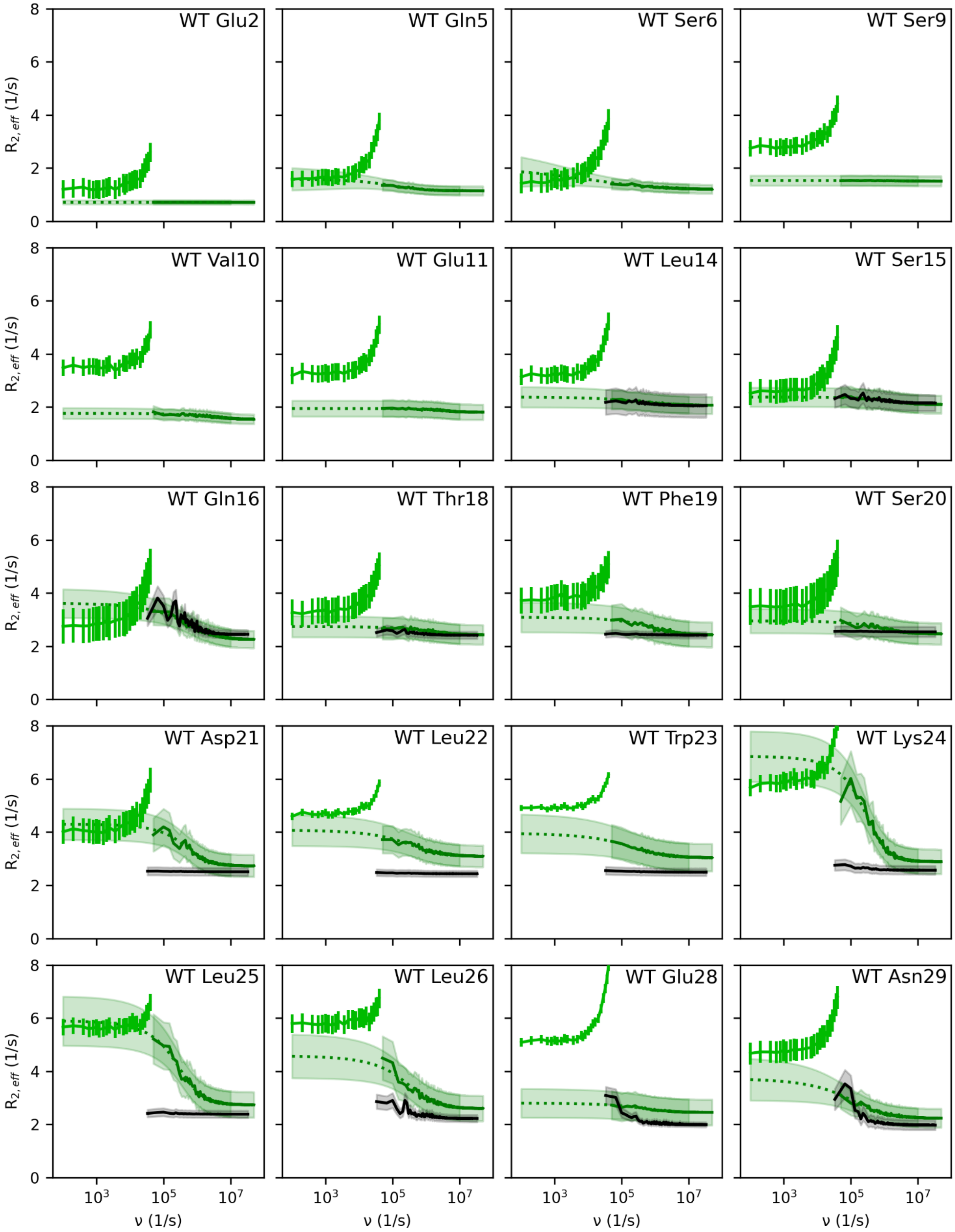
WT Glu2
WT Gln5
WT Ser6
WT Ser9
WT Val10
WT Glu11
WT Leu14
WT Ser15
WT Gln16
WT Thr18
WT Phe19
WT Ser20
WT Asp21
WT Leu22
WT Trp23
WT Lys24
WT Leu25
WT Leu26
WT Glu28
WT Asn29
$R_{2,eff}$ (1/s)
ν (1/s)
0
2
4
6
8
$10^3$
$10^5$
$10^7$

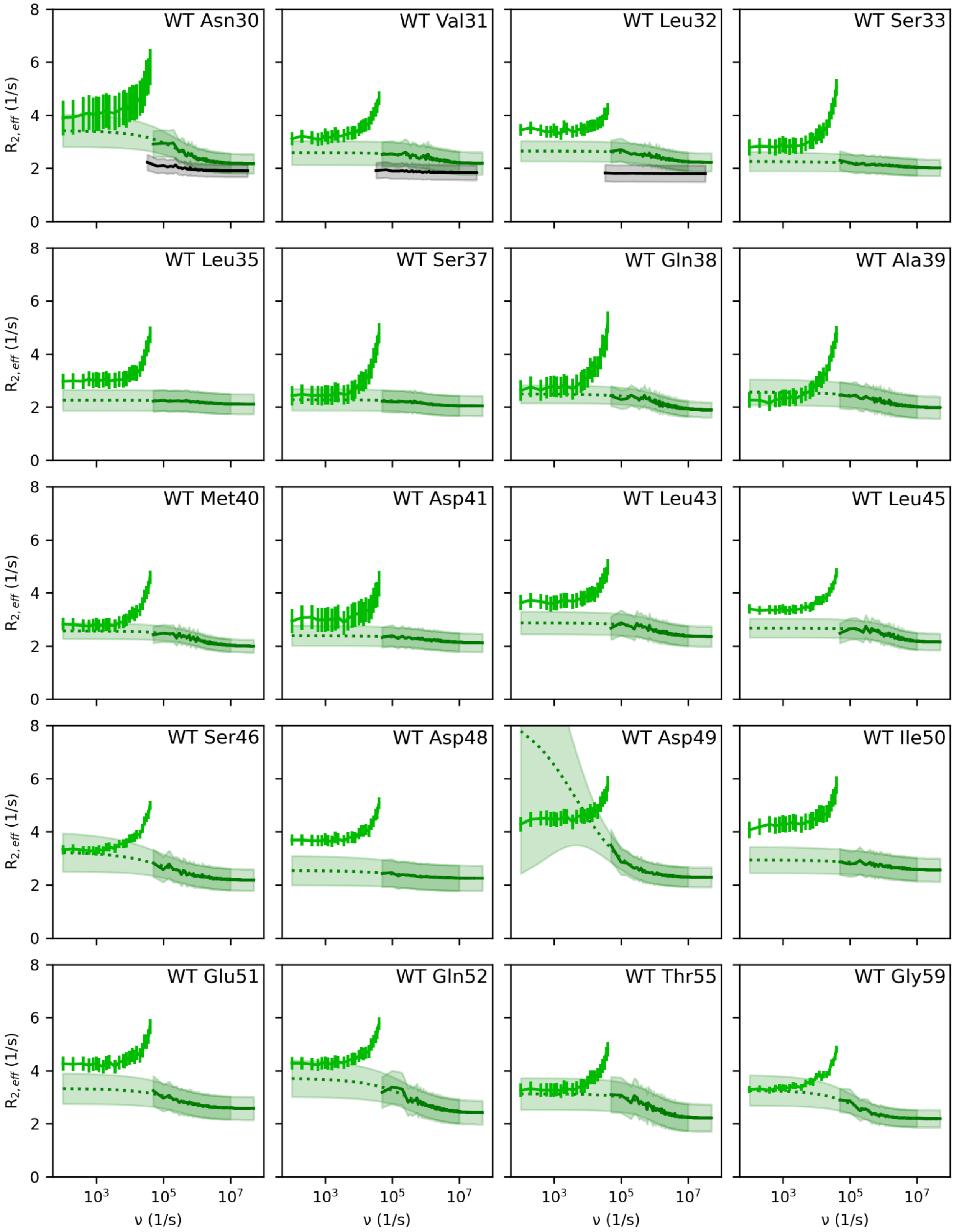
WT Asn30
WT Val31
WT Leu32
WT Ser33
WT Leu35
WT Ser37
WT Gln38
WT Ala39
WT Met40
WT Asp41
WT Leu43
WT Leu45
WT Ser46
WT Asp48
WT Asp49
WT Ile50
WT Glu51
WT Gln52
WT Thr55
WT Gly59
$R_{2,eff}$ (1/s)
ν (1/s)
0
2
4
6
8
$10^3$
$10^5$
$10^7$

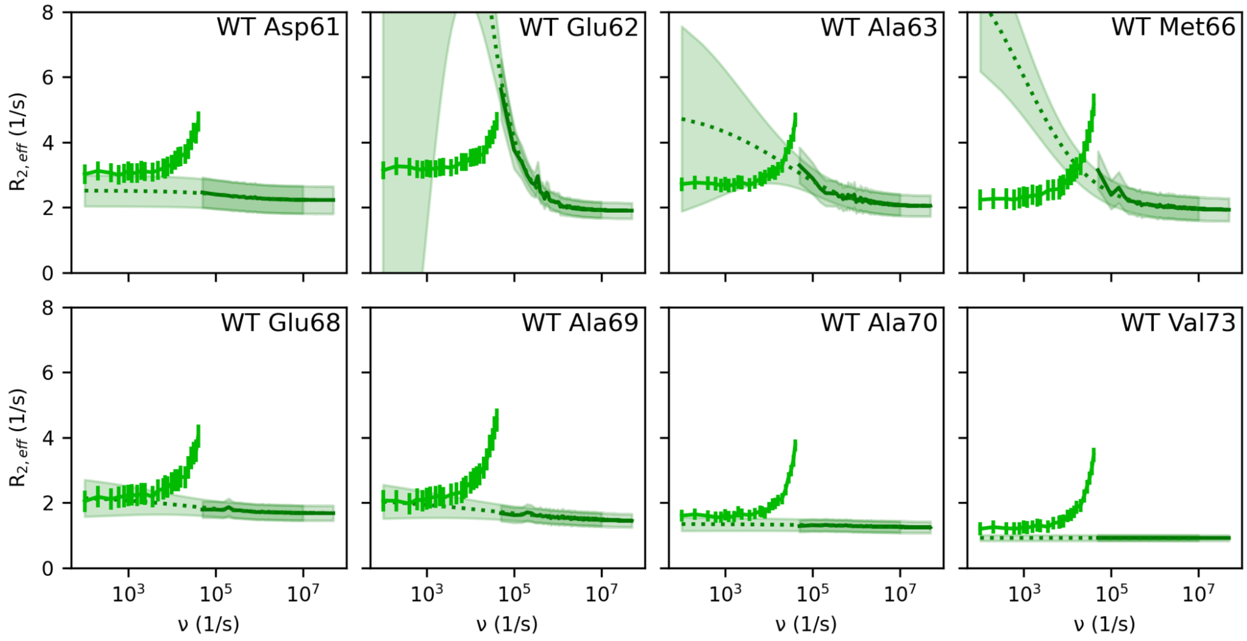

**Fig. S4. Measured vs. calculated p53-TAD WT RD profiles.** For each residue (panels), profiles derived from NMR measurements are shown in light green; profiles calculated from MD simulations are shown in dark green. To facilitate comparison, stretched CPMG-fits to the MD profiles are provided (dotted lines). Errors of the NMR measurements (vertical bars) as well as the statistical uncertainty of the MD profiles (shaded area) were estimated from the standard deviations of repeated measurements or trajectories. The uncertainty of the fits (shaded areas) was estimated from the standard deviations of a posterior sample obtained via Bayes fitting. For residues 14-32, which involve α-helix 1, additional simulations with a fixed ideal α-helix (black lines) were carried out using dihedral restraints, and RD profiles were calculated (black lines and shaded areas).

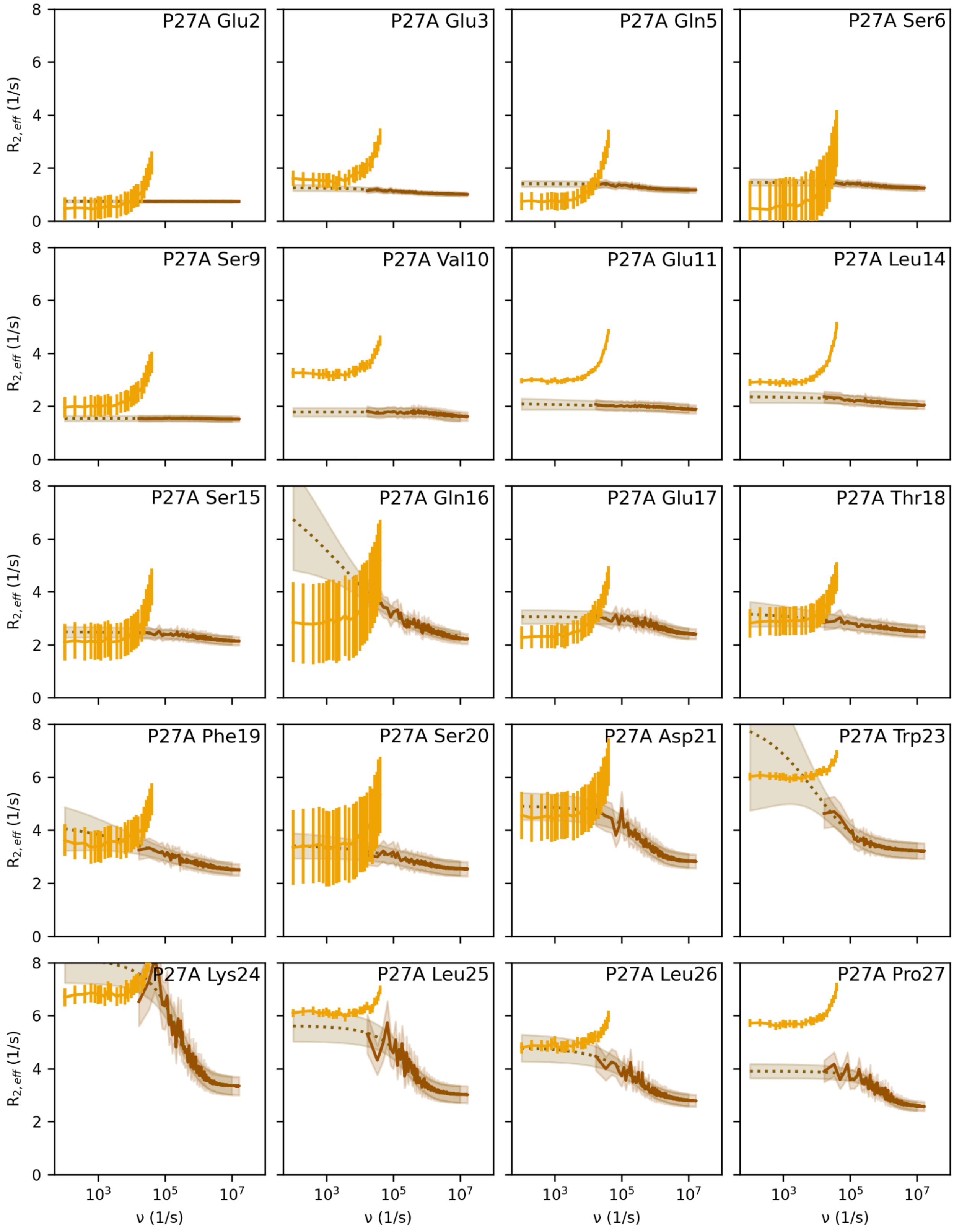

P27A Glu2
P27A Glu3
P27A Gln5
P27A Ser6
P27A Ser9
P27A Val10
P27A Glu11
P27A Leu14
P27A Ser15
P27A Gln16
P27A Glu17
P27A Thr18
P27A Phe19
P27A Ser20
P27A Asp21
P27A Trp23
P27A Lys24
P27A Leu25
P27A Leu26
P27A Pro27
$R_{2,eff}$ (1/s)
ν (1/s)

364

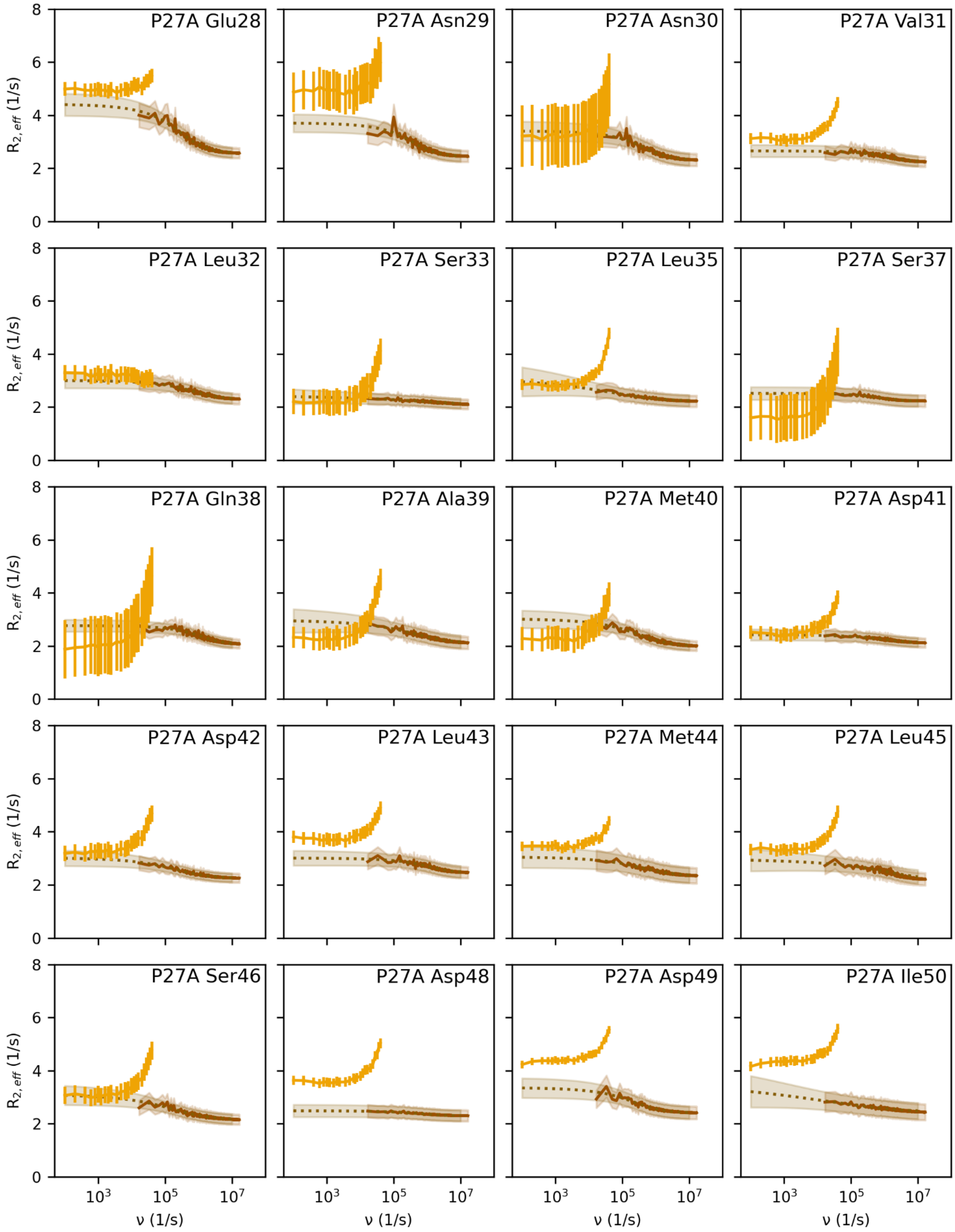
P27A Glu28
P27A Asn29
P27A Asn30
P27A Val31
P27A Leu32
P27A Ser33
P27A Leu35
P27A Ser37
P27A Gln38
P27A Ala39
P27A Met40
P27A Asp41
P27A Asp42
P27A Leu43
P27A Met44
P27A Leu45
P27A Ser46
P27A Asp48
P27A Asp49
P27A Ile50
$R_{2,eff}$ (1/s)
ν (1/s)

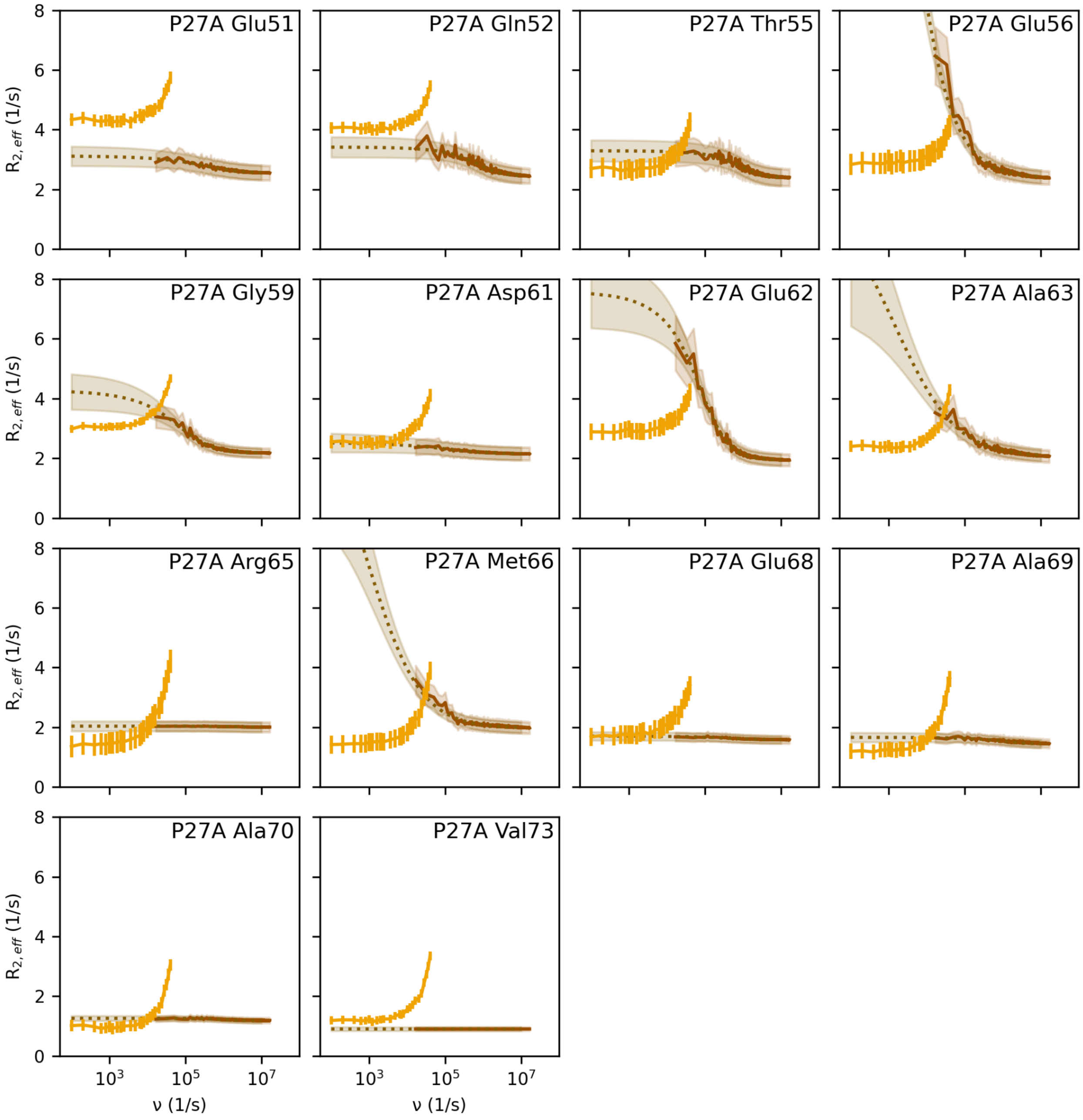

**Fig. S5. Measured vs. calculated p53-TAD P27A mutant RD profiles.** For description, see caption to fig. S4.

**(J) Methods: (b) Tumbling related relaxation: $R_{2,0}$**

To estimate $R_{2,0}$, the spectral density function (SDF) was calculated for each residue as described subsequently. From the SDF of each residue, $R_{2,0}$ was calculated via Eq. 13 and used as the offset for the calculated RD profiles.

**(K) Methods: Calculation of SDFs and derived NMR observables $\eta_{xy}$, $\tau_c$, $R_1$, $R_2$, and NOEs**

SDFs were calculated for each residue from the Fourier transformation

$$SDF(\nu) = \frac{1}{5} t_{max} Re\left(\mathcal{F}\big(C(\tau)\big)\right) , \quad (2)$$

of the backbone H-N-bond rotation-autocorrelation function $C(\tau)$ (*49, 96*)

$$C(\tau) = \langle P_2(\boldsymbol{\mu}(t) \cdot \boldsymbol{\mu}(t+\tau)) \rangle , \quad (3)$$

where $\tau$ is the lag time, $\boldsymbol{\mu}$(t) is the unit vector of the covalent bond, $P_2$ is the second Legendre polynomial, $t_{max}$ = 100 ns is the longest lag time considered, and ⟨ ⟩ denotes the time average along the trajectories. Due to the symmetry of the autocorrelation function, the real part of the Fourier transform is used for this calculation.

By combining the values of the SDF at specific frequencies, the transverse cross-correlation rate constant $\eta_{xy}$ and tumbling timescale $\tau_c$ were calculated using Eqs. (4-8), respectively, as described by Robson et al. (*66*),

$$\eta_{xy} = p\delta_N(4J(0) + 3J(\omega_N))(3cos^2\theta - 1) , \quad (4)$$

where

$$p = \frac{\mu_0 \, \gamma_H \, \gamma_N \, h}{16\pi^2 \sqrt{2}\, r^3} , \quad (5)$$

$$\delta_N = \frac{\gamma_N \, B_0 \, \Delta\delta_N}{3\sqrt{2}} , \quad (6)$$

$$J(\omega) = \frac{2\tau_c}{5[1+(\tau_c\omega)^2]} , \quad (7)$$

and $\mu_0$ = 1.27×10$^{-6}$ H m$^{-1}$, $\gamma_H$ = 267.52 rad s$^{-1}$ T$^{-1}$, and $\gamma_N$ = -27.12 rad s$^{-1}$ T$^{-1}$ are the gyromagnetic ratios of proton and nitrogen nuclei, respectively; $h$ = 6.63×10$^{-34}$ Js is Planck's constant, $\Delta\delta_N$ = 160 ppm, $r_{NH}$ = 1.02 Å, $B_0$ = 28.19 T, and $\theta$ = 17°.

Similarly (*66*),

$$\tau_c = \frac{5c}{24} - \frac{336\omega_N^2 - 25c_1^2\omega_N^4}{24\omega_N^2\left(1800c_1\omega_N^4 + 125c_1^3\omega_N^6 + 24\sqrt{3}\sqrt{21952\omega_N^6 - 3025c_1^2\omega_N^8 + 625c_1^4\omega_N^{10}}\right)^{\frac{1}{3}}} +$$

$$\frac{\left(1800c_1\omega_N^4 + 125c_1^3\omega_N^6 + 24\sqrt{3}\sqrt{21952\omega_N^6 - 3025c_1^2\omega_N^8 + 625c_1^4\omega_N^{10}}\right)^{\frac{1}{3}}}{24\omega_N^2}\ , \quad (8)$$

where

$$c_1 = \frac{\eta_{xy}}{p\delta_N(3\cos^2\theta - 1)}\ . \quad (9)$$

Next, the rate constants $R_1$ and $R_2$ were calculated via Eqs. (10-15) as described by Palmer III. (*20*),

$$R_1 = \frac{d_2^2}{4}(3J(\omega_N) + J(\omega_m) + 6J(\omega_p)) + c_2^2 J(\omega_N)\ , \quad (10)$$

where

$$c_2 = \frac{\omega_N \Delta\delta_N}{\sqrt{3}}\ , \quad (11)$$

$$d_2 = \frac{\mu_0 \gamma_H \gamma_N \hbar}{8\pi^2 r^3}\ , \quad (12)$$

and $\omega_m = \omega_H - \omega_N$ and $\omega_p = \omega_H + \omega_N$ ,

as well as

$$R_2 = \frac{d_3}{8}\left(4J(0) + 3J(\omega_H) + J(\omega_m) + 6J(\omega_N) + 6J(\omega_p)\right) + \frac{1}{6}c_3\omega_H^2\left(4J(0) + 3J(\omega_H)\right)\ , \quad (13)$$

where

$$c_3 = \frac{\Delta\delta H^2}{3}\ , \quad (14)$$

$$d_3 = \left(\frac{\mu_0}{4\pi}\right)^2\left(\frac{h}{2\pi}\right)^2 \gamma_H^2\gamma_N^2 r^{-6}\ , \quad (15)$$

and $\Delta\delta H = 10^{-5}$ is the difference in the axially symmetric proton chemical shift tensor.

Finally, NOEs were calculated as described by Farrow et al. (*97*),

$$NOE = 1 + \frac{d_3^2\gamma_H}{4\gamma_N}\left(6J(\omega_p) - J(\omega_m)\right)\frac{1}{R_1}\ , \quad (16)$$

where $d_3$ is defined in Eq. (15) and $R_1$ in Eq. (10).

The code used for all the above calculations, as well as sample calculations and sample data have been deposited at the public repository GitHub https://github.com/dszollosi/p53_TAD_dynamics.

### (L) Results: Comparison of measured with calculated NMR observables $\eta_{xy}$, $\tau_c$, $R_1$, $R_2$, and NOEs

Figure S6 compares, per residue, calculated with measured NMR parameters $\eta_{xy}$, $\tau_c$, rate constants $R_1$, $R_2$, and NOE values. Both $R_1$ and NOEs were measured at a 950 MHz magnetic field strength, whereas $\eta_{xy}$, $\tau_c$, and $R_2$ were measured at 1.2 GHz. Accordingly, these field strengths were also used for the corresponding values calculated from the MD simulations.

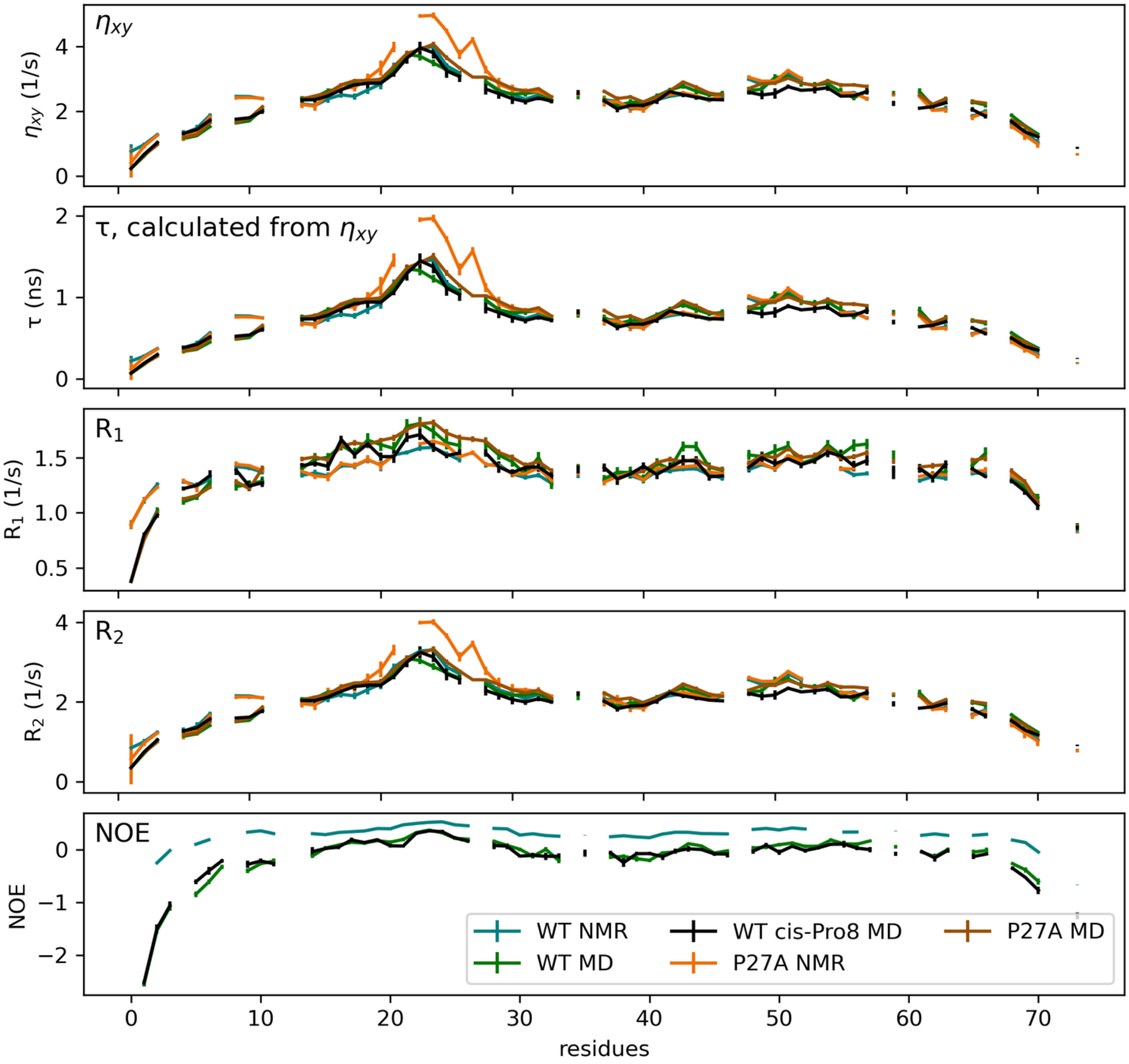

**Fig. S6. Comparison of measured with calculated NMR observables.** Comparison of $\eta_{xy}$, $\tau_c$, the rate constants $R_1$, $R_2$, and NOE values derived from NMR measurements with values calculated from MD simulations for the p53-TAD WT (turquoise and green lines) and the P27A mutant (orange and brown lines). A cis-isomer configuration of Pro8 (black line) shows no significant difference to the trans isomer of the WT (green line). $R_1$ and NOE values were measured at a magnetic field strength of 950 MHz; $\eta_{xy}$, $\tau_c$, and $R_2$ at 1.2 GHz. The values calculated from our MD simulations used corresponding magnetic field strengths. Lines show the mean of the trajectory-wise calculated observables with error bars (vertical bars). Missing data points are due to proline residues without backbone amide protons, for which therefore no spectra were obtained.

**(M) Results: SDFs calculated from MD simulations; comparison with NMR measurements and a simple tumbling model**

The considerable length of our MD trajectories enabled us to calculate the spectra density function (SDF) $J(\nu)$, which is not directly accessible to NMR measurements. For each residue over a broad frequency range covering more than four orders of magnitude (figs. S7, green lines) $J(\nu)$ was estimated. For comparison to the experiment, and using multiple NMR measurements at various conditions, for each residue we calculated values of the SDF $J(0)$, $J(\omega_N)$, and $J(\varepsilon\omega_H)$ at three distinct frequencies, as described earlier by Farrow et al. (*97*). The horizontal black line indicates the SDF value for $\nu$ = 0; the dots depict values at frequencies $\nu_1 = \omega_N/2\pi$ = 96.3 MHz and $\nu_2 = \varepsilon\omega_H/2\pi$ = 826.5 MHz, respectively.

Note that, following the convention in the literature, frequencies $\nu$ are given in units 1/s, whereas frequencies $\omega$ are in units rad/s. To facilitate easier comparison with low-frequency spectra, e.g., from CMPG experiments, and unfortunately contrary to common usage, we show high frequency spectra such as the SDF also as a function of $\nu$ and in units of 1/s.

In fig. S7 we also compared the SDF calculated from our simulations with an analytical SDF (blue), calculated for a simple model (Eq. (7)) assuming the underlying dynamics are described by a single tumbling timescale $\tau_c$. To this end, $\tau_c$ was measured by the TRACT experiment described above. Notably, the SDF calculated from our MD simulations are stretched relative to the Lorentzian functions of the simple model, as evident from the shallower slope at higher frequencies. This result is in line with a similar observation for RD profiles (fig. S10), which strongly suggests that also the tier 2 reorientation dynamics (which determine the SDF) comprise not one but many timescales.

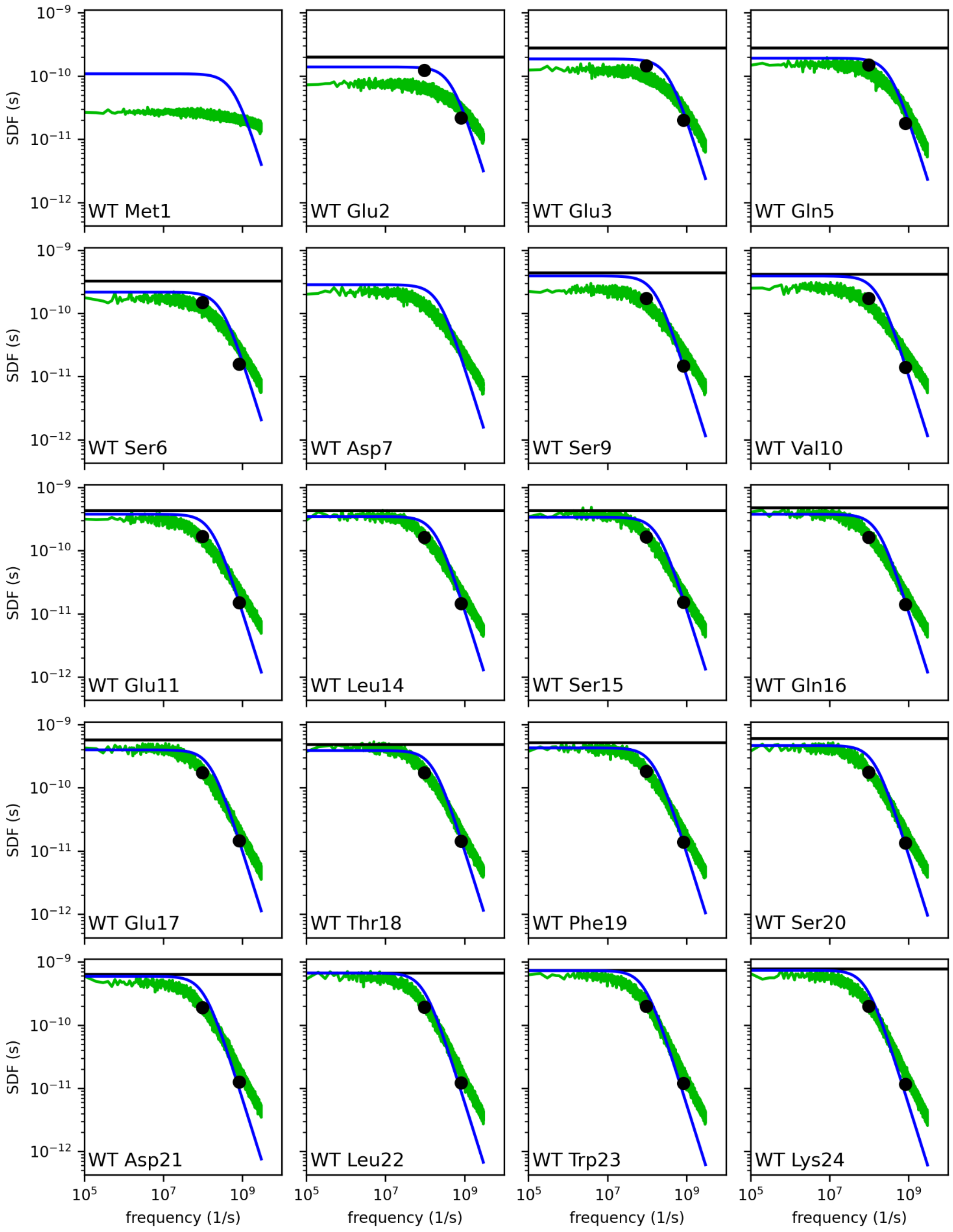

SDF (s)
frequency (1/s)
WT Met1
WT Glu2
WT Glu3
WT Gln5
WT Ser6
WT Asp7
WT Ser9
WT Val10
WT Glu11
WT Leu14
WT Ser15
WT Gln16
WT Glu17
WT Thr18
WT Phe19
WT Ser20
WT Asp21
WT Leu22
WT Trp23
WT Lys24

477

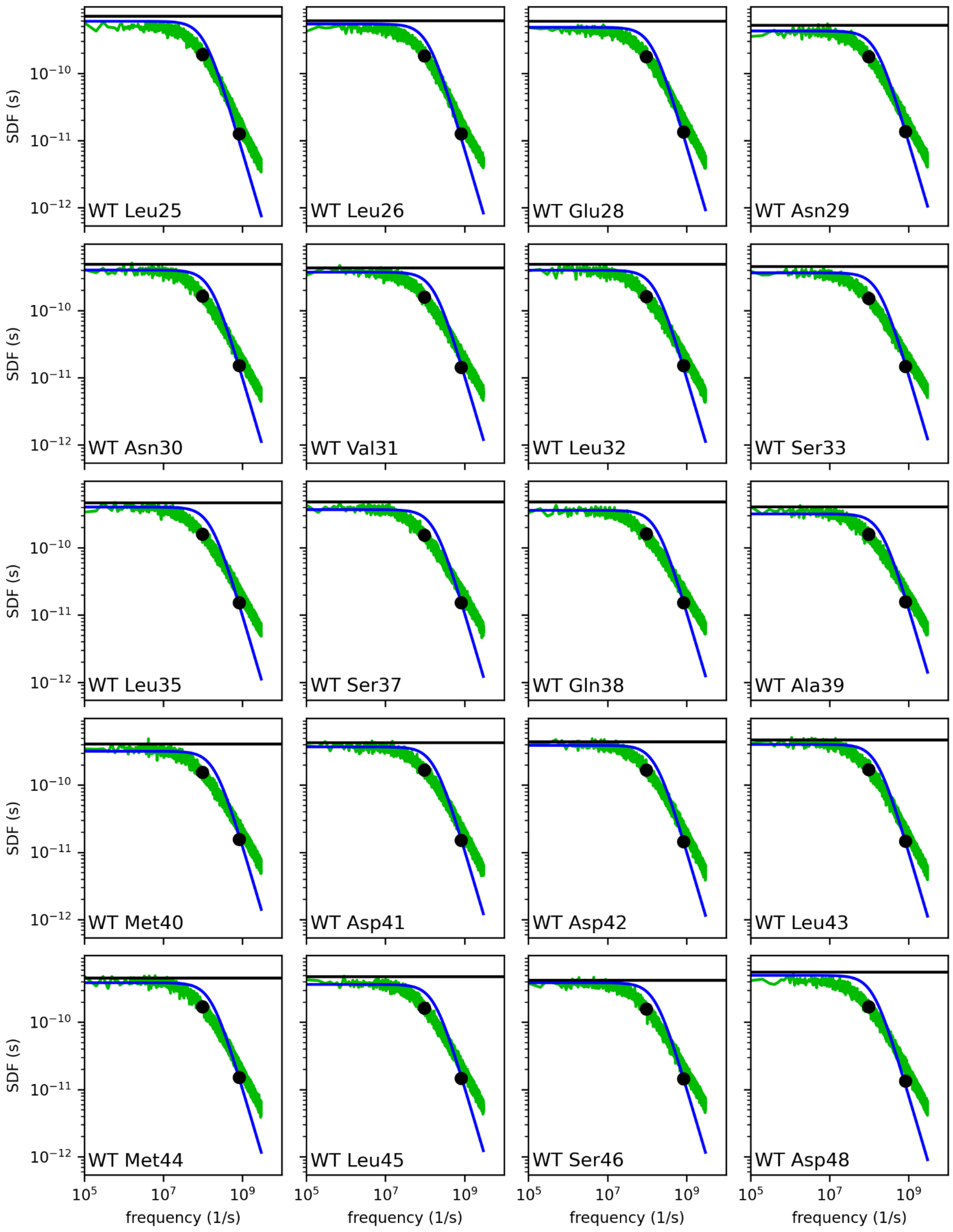
WT Leu25
WT Leu26
WT Glu28
WT Asn29
WT Asn30
WT Val31
WT Leu32
WT Ser33
WT Leu35
WT Ser37
WT Gln38
WT Ala39
WT Met40
WT Asp41
WT Asp42
WT Leu43
WT Met44
WT Leu45
WT Ser46
WT Asp48
SDF (s)
$10^{-10}$
$10^{-11}$
$10^{-12}$
$10^{5}$
$10^{7}$
$10^{9}$
frequency (1/s)

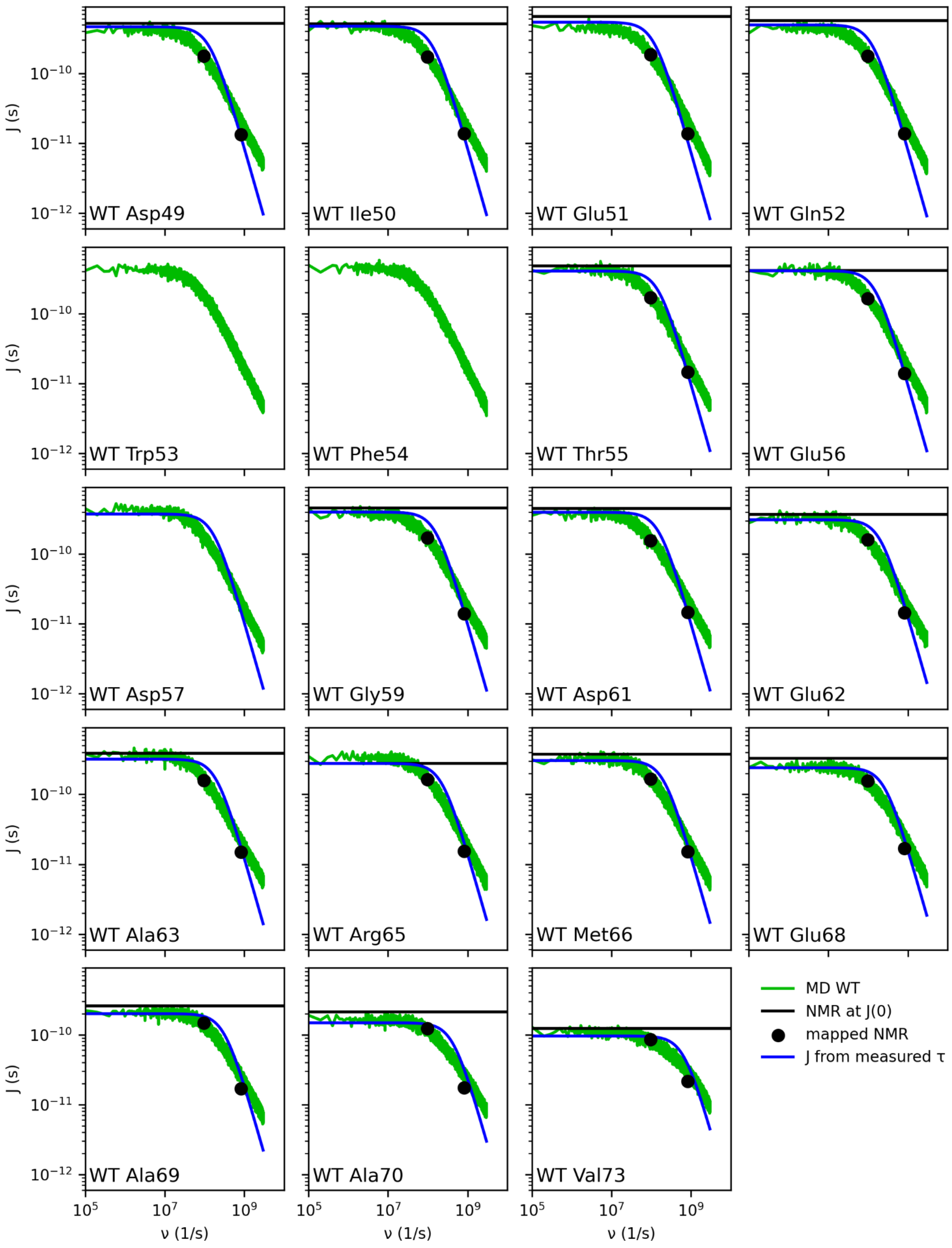

**Fig. S7. Calculated SDFs vs. NMR experiment.** Shown are SDFs $J(\nu)$ calculated from our MD simulations (green lines) and measured values $J(\nu)$ at frequency 0 (horizontal black lines) as well as at frequencies $\nu_1 = \omega_N/2\pi = 96.3$ MHz and $\nu_2 = \varepsilon\omega_H/2\pi = 826.5$ MHz, respectively (black dots) for each residue of the p53-TAD WT, as indicated in each panel. Blue lines show SDFs calculated from a simple state model, for which the relaxation time $\tau_c$ determined from TRACT experiments was used. Note the different ranges on the y-axes.

To facilitate easier quantitative comparison between the measured and calculated SDFs, fig. S8 shows the same data as fig. S7 for the three frequencies 0, $\nu_1 = \omega_N/2\pi$ = 96.3 MHz, and $\nu_2 = \varepsilon\omega_H/2\pi$ = 826.5 MHz, using the same color code as in fig. S7. At $\nu$ = 0 (fig. S8A), the $J(0)$ calculated from our MD simulation (green) agrees well with the measured ones (black), and both also agree well with the simple model (blue). Notably, at $\nu_1$ = 96.3 MHz (fig. S8B), calculated and measured values agree very well, the simple model (blue line) consistently predicts larger values, which, again results from the different slopes of the high-frequency parts, underscoring the multi-timescale reorientational dynamics of P53-TAD within tier 2. At the highest frequency ($\nu$ = 826.5 MHz, fig. S8C), the simple model predictions agree well with the measured values, which, however, is largely accidental, due to the fact that this frequency is often close to the intersection point of the two spectra (cf. fig. S7).

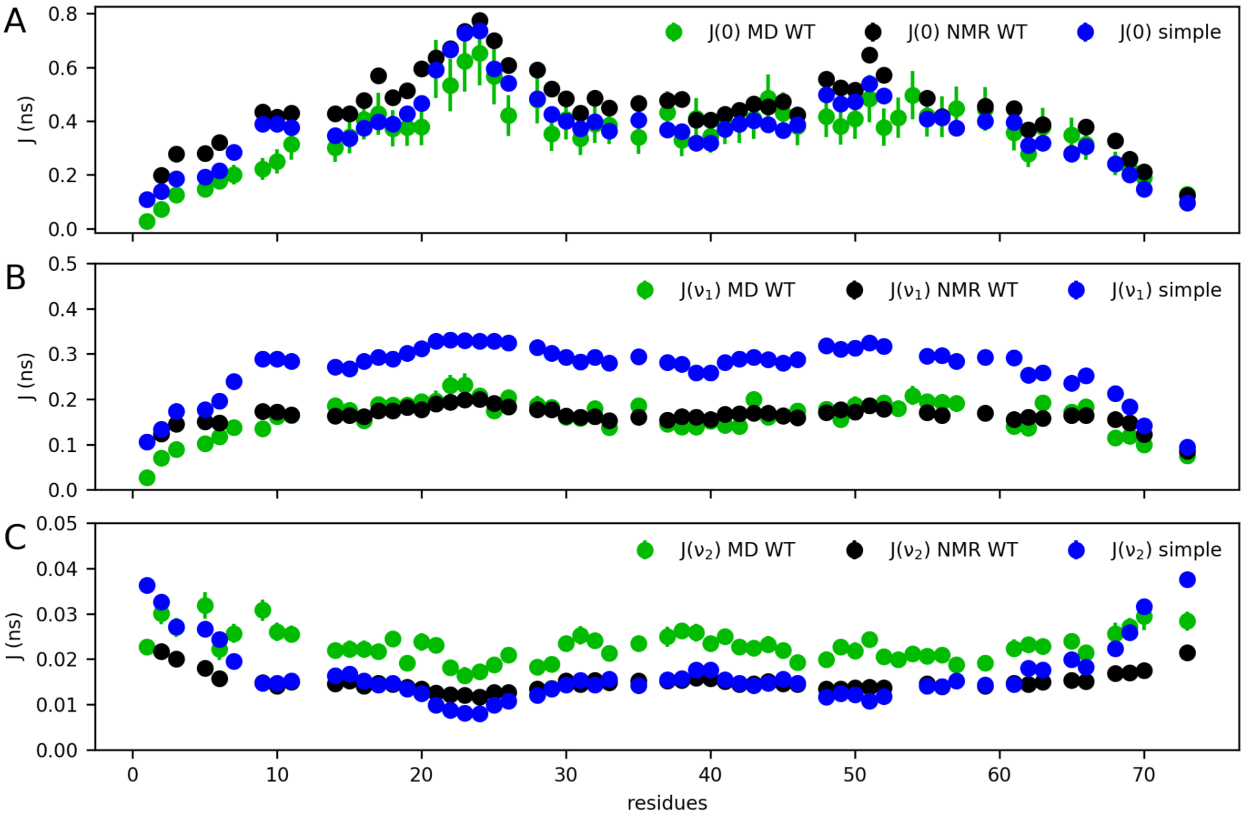

**Fig. S8. Summary of calculated SDFs vs. NMR experiment.** Shown are SDF values $J(0)$ (**A**), $J(\nu_2)$ (**B**), and $J(\nu_2)$ (**C**) at those frequencies (cf. fig. S7) that were experimentally accessible. Each panel compares SDF values calculated from our MD simulations (green) with measured ones (black) and those predicted from a simple model (blue) using measured values for $\tau_c$ from NMR. Errors are indicated as vertical bars and are otherwise smaller than the symbols.

The predictions calculated from our MD simulations yield a very similar profile as the measured ones, but are consistently too large. The reason for this discrepancy remains unclear; we speculate that at fast nanoseconds timescales, the used Amber99sbws force field is known to produce dynamics that are slightly too slow at least in helix formation (cf. Table 2 in Sorin & Pande 2005 (*98*)) and, according to equipartition, exhibits higher amplitudes at these fast frequencies. In addition to this systematic deviation, larger differences are seen for the N-terminal residues up to Val10, likely due to the few residues added during cloning, which are absent in the simulations.

**(N) Results: Effects of additional residues at the N-terminus**

Our study aims to understand the hierarchical dynamics of the WT p53-TAD in its native form, and to compare these dynamics to the P27A mutant. Therefore, in our simulations we have constructed the respective peptides. In contrast, for technical reasons during the cloning process, the peptides used in the experiments included additional amino acids at the N-terminus, namely a Gly-Ser-extension to the WT sequence and a Gly-Ser-His-Met-extension to the P27A mutant. To assess the effect of these additional residues on the p53-TAD dynamics, we performed additional MD simulations of 5 μs each of the extended P27A construct. Specifically, five simulations each were performed for the two most likely protonation states of the histidine, singly protonated (neutral) and doubly protonated (charge +1). Simulation parameters and the analysis were as for the non-extended sequences. Due to shorter simulation lengths of the extended constructs, we compared the SDF at the frequency $\omega_N$, which showed faster convergence and for the WT where measurements were available.

As can be seen in fig. S9, SDF values measured by NMR agree very well with those calculated from both simulations of the (identical) extended construct, now also for all N-terminal residues. Whereas the values obtained from the MD simulations that lack the extension also agree well with both the measured ones as well as the other simulations for the whole peptide up to Val10, differences for the N-terminal residues are clearly visible. We conclude that most of the N-terminal deviations reported in the main text are due to the presence of these 2-4 additional residues in the measurements. Notably, the SDF values calculated for both possible protonation states are very similar to each other (as well as to the measured NMR values) even close to the histidine at the N-terminus, which suggests that these do not markedly affect the dynamics of the peptide.

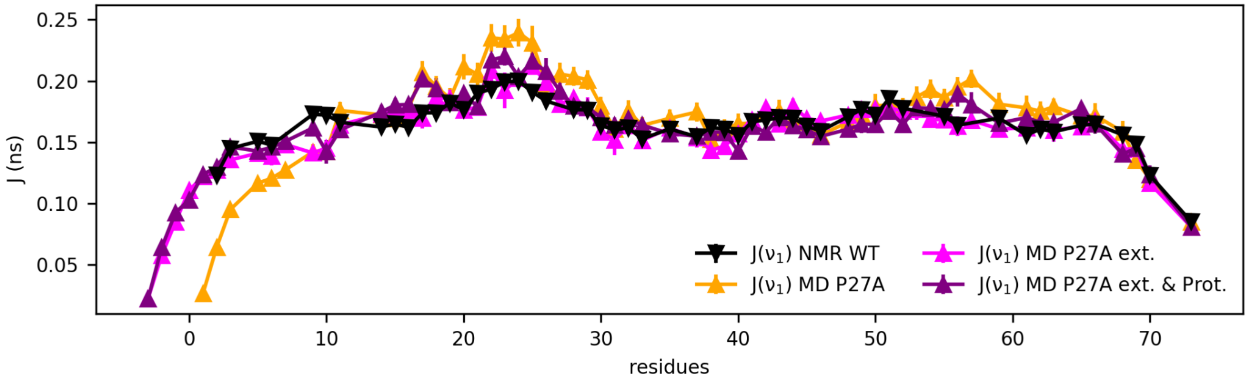

**Fig. S9. Comparison of the extended vs. non-extended p53-TAD construct.** Residue numbering is according to the original p53-TAD WT sequence. Shown are SDF values at frequency $\omega_N$ for the non-extended p53-TAD WT measured by NMR (black), compared to SDF values obtained from our MD simulations for the non-extended P27A (orange), for the P27A extended (magenta) and for the P27A extended protonated (purple) constructs. Errors are indicated as vertical bars and are otherwise smaller than the symbols.

### (O) Methods: Fitting of 2-state CPMG model and stretched CPMG model to RD profiles calculated from MD simulations

To compare our NMR CPMG measurements with our MD simulations not only in terms of the raw spectra (see above), but also in terms of the characteristic exchange dynamics timescales $\tau$, the analytical form of CPMG profiles for a two-state model (Eq. 17) (*43, 99*)

$$R_{2,MD}(\nu) = \phi\tau\left(1 - 4\nu\tau\, tanh\frac{1}{4\nu\tau}\right), \quad (17)$$

was fitted both to the measured NMR RD spectra as well as to those calculated from our MD simulations. Here, $\nu$ is the CPMG frequency, $\Phi = p_A\, p_B\, \Delta CS^2$ is the population weighted chemical shift variance, $p_A$ and $p_B$ are the populations of the two states A and B of the model, respectively, and $\Delta CS$ is the chemical shift difference between these two states.

Inspired by the shape of the RD profiles calculated from the simulations (cf. Fig. S4), which in the logarithmic plot appears to be 'stretched' in the frequency domain relative to the above CPMG two-state model, a generalized model in terms of a 'stretched' function was considered,

$$R_{2,MD}(\nu) = \phi\tau\left(1 - (4\nu\tau)^{\gamma}\tanh\frac{1}{(4\nu\tau)^{\gamma}}\right), \quad (18)$$

similar in spirit to the 'stretched exponential functions' used by Frauenfelder to describe the multi-tier dynamics of folded proteins (*5, 100, 101*). Similar to what has been shown by (*102*), Eq. (18) can be interpreted as a weighted superposition of many two-state CPMG profiles, Eq. (17), with a broad range of characteristic timescales, which is described by the additional fitting parameter $\gamma$. Accordingly, $\gamma$ also characterizes the distribution width of barrier heights governing p53-TAD dynamics, namely $\gamma$ = 1 indicate a two-state process and the smaller the value the broader the timescale distribution is. Fits of Eqs. (17) and (18) to either measured or calculated RD profiles were performed using a Bayesian approach (*103*) using the pymc5 Python package (*104*), with $\tau$ and $\Phi$ (and, additionally, $\gamma$ for Eq. (18)) as free parameters to be determined by the fits.

The fact that all RD profiles were calculated by averaging over 30 individual RD profiles, each calculated from an absolute-squared Fourier transform of chemical shift trajectories as described above, required particular attention. Notably, Fourier transforms calculated numerically from finite time series are notoriously noisy, and the distribution of each individual (absolute squared) Fourier coefficient $R_k := R_2(\nu_k)$ for realizations with a uniform distribution of random phases follows an exponential function (*105*),

$$p(R_k) = \frac{1}{R_k^0} e^{-\frac{R_k}{R_k^0}} ,$$

where $p(R_k^0)$ is the true coefficient from which the realizations were drawn. Hence, the probability distribution of the mean $\bar{R}_k$ of $N$=30 Fourier transforms of realizations of the same process (the phases of which are assumed to be statistically independent and uniformly distributed) follows a gamma distribution

$$p(\bar{R}_k) = \frac{\beta^N \bar{R}_k^{N-1} e^{-\beta \bar{R}_k}}{\Gamma(N)} , \quad (19)$$

where $\beta = \frac{N}{\bar{R}_k}$ .

Accordingly, this probability distribution was used (instead of a Gaussian distribution) for the Bayesian fit of each Fourier coefficient, with the fitting target $\bar{R}_k$. A log-uniform prior distribution between $10^3$-$10^9$ Hz$^2$ was used for $\Phi$, a log-uniform prior between 1 ns-100 μs for $\tau$, and a uniform prior between 0-2 for $\gamma$. An additional jupyter notebook describes in detail all fitting steps, available at the public repository GitHub https://github.com/dszollosi/p53_TAD_dynamics.

This Bayes approach was also used for the fits to measured NMR relaxation dispersion profiles, except that a Gaussian function was used as an error model instead of the above gamma function.

For the residues 49ASP, 54PHE, 56GLU, 57ASP, 62GLU, 63ALA and 66MET, the RD profile does not reach a plateau at low frequencies, indicating very slow dynamics (cf. the respective panels in figs. S4 and S5) which are also apparent from the Bayes posteriors. For these residues, therefore, no relaxation timescales were derived from the fits, which are therefore also not shown in Fig. 2B.

**(P) Results: Fitting CPMG model to RD profiles calculated from MD simulations**

As an illustrative example, figs. S10 A-D show the RD profile calculated from MD simulations for the Leu25 residue (green) fitted with a non-stretched (A and C, cyan and blue, Eq. 17) and a stretched (B and D, purple, Eq. 18) CPMG function, respectively. For the non-stretched CPMG fit, two attempts were made, one with equal weights on all data points of the calculated profile (cyan), and a second one with weights inversely proportional to the frequency (blue). For better visual assessment of the fits both at low and high frequencies, they are shown on a linear (fig. S10 A-B) as well as on a logarithmic (fig. S10 C-D) scale.

The former fit agrees with the high-frequency part of the profile, but fails for the low-frequency part which contains fewer data points and, hence, carries little weight for the fit. The latter fit, vice versa, approaches the low-frequency part, but fails for the high-frequency part of the calculated profile. In contrast, the stretched function approximates the calculated profile well for the whole frequency range. Figure S10E shows the mean stretch parameter $\gamma$ for all p53-TAD WT residues, as obtained from similar stretched CPMG fits. Missing data points are due to proline residues without backbone amide protons, for which therefore no spectra could be recorded.

Figure S10F illustrates a plausible interpretation of why the RD profile is best described by a stretched CPMG function, in terms of conformational dynamics between many conformational states, which are indeed seen in our trajectories. Generalizing the well-known analytical results for a two-state Markov process, such multi-state conformational dynamics can be described by a multi-state Markov process (*106*). Accordingly, and as has been shown previously (*101, 102, 107*), the RD profile (power spectrum) of such a Markov process with $M$ states is a weighted superposition of $M$-1 Lorentzian (or here, CPMG) functions, with the $M$-1 non-zero eigenvalues of the Markov matrix representing the respective relaxation times $\tau$.

As an example, fig. S10F shows a superposition of 11 non-stretched CPMG functions (blue to green lines) with 11 relaxation times $\tau_i$ ($i$ = 1…11) log-uniformly distributed between 60 ns and 2.5 µs and weights 1/$i$. As can be seen, the sum of these 11 CPMG functions (bold green line) is well approximated by a stretched CPMG function (purple) with a typical stretch parameter $\gamma$ = 0.65, taken from the above fits. Incidentally, this time range overlaps largely with the characteristic times extracted from the Markov model shown in Fig. 4A-D.

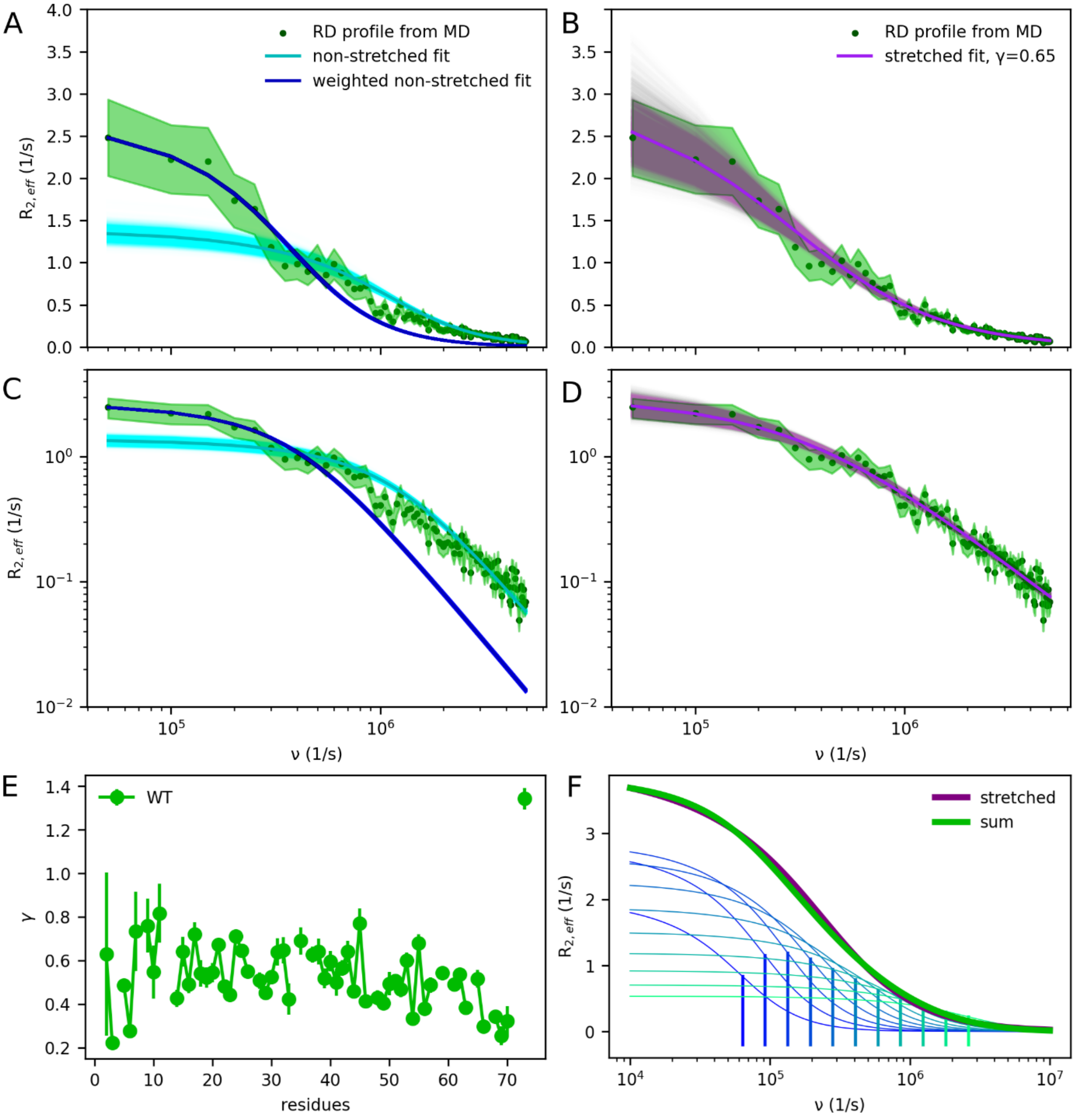

**Fig. S10. Stretched CPMG fits to RD profiles reflect multi-state multi-timescale conformational dynamics.** The p53-TAD WT RD profile of the Leu25 residue (green), calculated from 30 MD trajectories, was fitted with a non-stretched CPMG function with equal weights for each data point (solid cyan line) and with weights proportional to $1/\nu$ (solid blue line) (**A**, **C**). The same RD profile was also fitted by a stretched CPMG function (solid purple line) (**B**, **D**). For better visual inspection, profiles and fits are shown on a linear (**A**, **B**) as well as on a logarithmic scale (**C**, **D**). The statistical uncertainty of the calculated profile is shown as a green area around the mean (green dots). Transparent cyan, blue and purple regions show the fitting uncertainty, represented by Bayesian posterior ensembles; the respective best (average) fit is shown as an opaque cyan, blue and purple line. (**E**) Mean stretch parameter $\gamma$ for all p53-TAD WT residues, obtained from similar fits to profiles calculated for each residue from our MD simulations; error bars indicate the standard deviation of each parameter. (**F**) A superposition (green line) of 11 non-stretched CPMG functions (blue to green lines, see text) with log-uniformly distributed relaxation times (vertical bars) is well approximated by a stretched CPMG function (purple line).

**(Q) Results: Accuracy assessment of the unbiased MD ensemble**

In addition to the above comparisons to NMR measurements, we assessed the accuracy of the ensemble derived from our unbiased MD simulations by comparing it to data derived from further independent calculations and experiments collected from the literature. These were (a) chemical shift predictions, (b) ensemble averaged size determined by size exclusion chromatography (SEC), small-angle X-ray scattering (SAXS), and dynamic light scattering (DLS), (c) distances determined by fluorescence resonance energy transfer (FRET), (d) photoinduced electron transfer fluorescence correlation spectroscopy (PET-FCS), and (e) paramagnetic relaxation enhancement (PRE).

***(a) Predicted chemical shifts***

Chemical shifts were calculated from our p53-TAD WT trajectories using SPARTA+ (*90*). To assess the accuracy of these empirical predictions, we also calculated chemical shifts using SHIFTX2 (*108*) and obtained almost identical chemical shifts, deviating by ca. 2.5% on average. For a small fraction of the structures, and for unknown reason, chemical shifts below 5 ppm were predicted incorrectly by SHIFTX2, as indicated by a discontinuous chemical shift distribution, which therefore were excluded from the comparison. All further analyses and comparisons with experiments were carried out with chemical shifts calculated by SPARTA+.

Next, calculated chemical shifts, averaged over our MD trajectories and compared to own NMR measurements as well as to chemical shifts by Wong et al. (*109*) (fig. S11). For the $C_\alpha$ chemical shifts, very good agreement is seen, with Pearson correlations of 0.996 and a mean absolute error (MAE) of 0.26 ppm and 0.24 ppm (fig. S11, A and C), respectively. Proton chemical shifts, from which RD profiles were calculated, show a Pearson correlation of over 0.76 and a MAE of 0.13 ppm (fig. S11, B and D). We note that the amplitudes of the RD spectra are sensitive to the somewhat larger uncertainty of the predicted proton chemical shifts and, therefore, the comparison of calculated to measured RD profiles provides an independent accuracy assessment. In contrast, relaxation times derived from calculated RD spectra are insensitive to such inaccuracies.

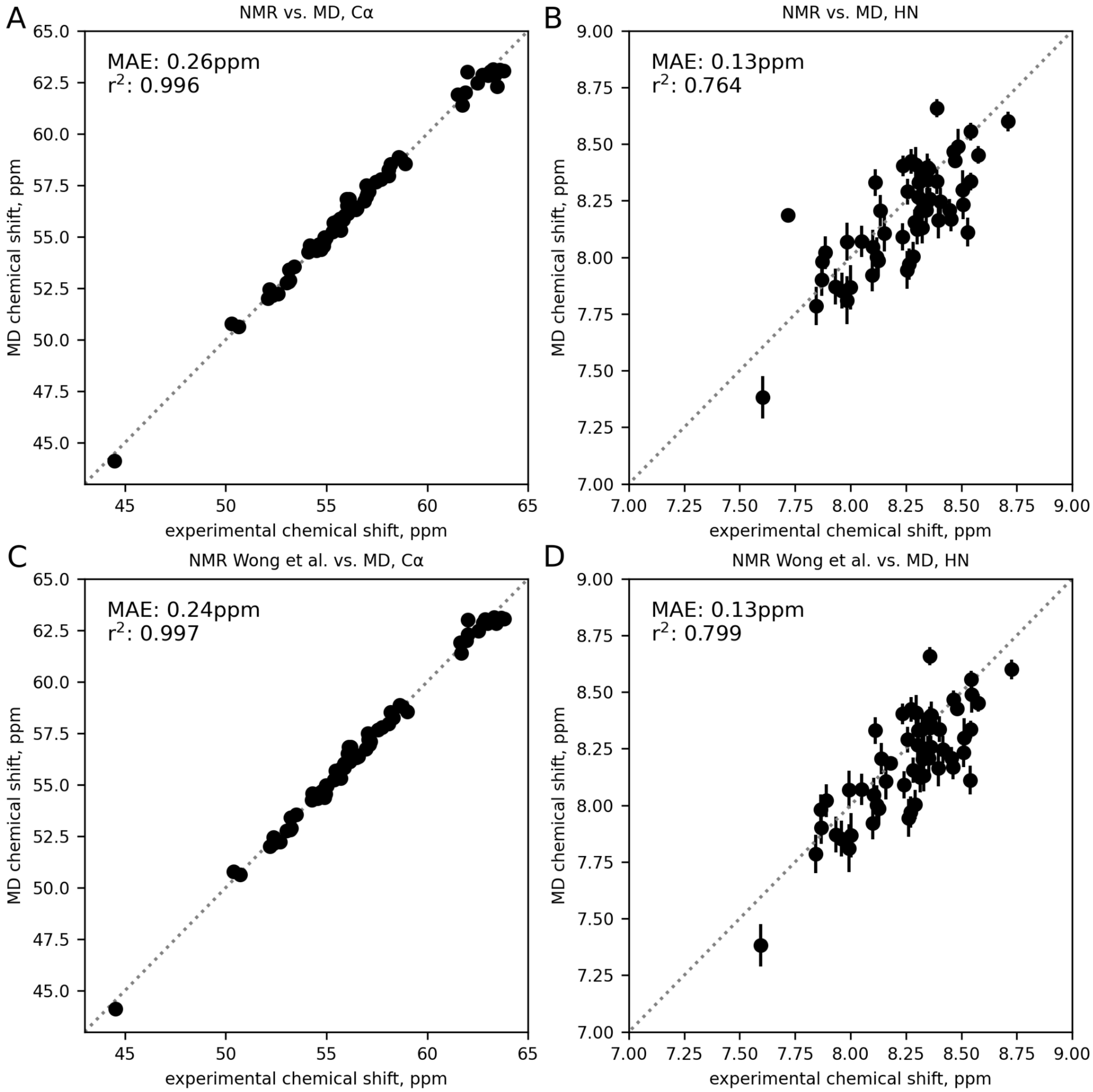

**Fig. S11. Comparison of predicted with measured p53-TAD chemical shifts.** $C_\alpha$ (**A**, **C**) and HN (**B**, **D**) chemical shifts in ppm (circles) derived from MD simulations compared to chemical shifts from two independent NMR measurements, our own (**A**, **B**) and published by Wong et al. (*109*) (**C**, **D**). Pearson correlation coefficients and mean absolute errors (MAE) are shown in each panel; a linear fit (dotted line) is shown for comparison. Error bars show, for each nucleus, the standard error of the mean of the chemical shifts calculated for all structures of the MD ensemble. For the $C_\alpha$ chemical shifts, the error bar is smaller than the symbol size.

***(b) Ensemble averaged radius of gyration and hydrodynamic radius***

To further assess the accuracy of the p53-TAD (1-73) ensemble generated from our MD simulations, we calculated ensemble averaged (and distributions of) hydrodynamic radii and radii of gyration and compared these with three independent experiments, namely hydrodynamic radii derived from published SEC and DLS experiments, and radii of gyration derived from SAXS measurements.

First, the hydrodynamic radius $R_h$ was determined from the radius of gyration $R_g$ computed from the obtained structural ensemble by adopting the approach of Nygaard et al. (*110*),

$$\frac{R_g}{R_h}(N, R_g) = \frac{\alpha_1(R_g - \alpha_2 N^{0.33})}{N^{0.6} - N^{0.33}} + \alpha_3 \quad , \quad (20)$$

where N = 73 is the number of p53-TAD residues, 0.33 (folded proteins) and 0.6 (disordered proteins) are the fitting parameters as previously published by Nygaard et al., and $\alpha_1$ = 0.216 Å$^{-1}$, $\alpha_2$ = 4.06 Å, and $\alpha_3$ = 0.821 Å.

Second, $R_h$ was determined from $C_\alpha$-$C_\alpha$ distances obtained from simulation trajectories by applying the Kirkwood equation (*111*), corrected for the 19% underestimation of $R_h$ due to the missing hydration shell according to Nygaard et al. (*110*).

Third, $R_h$ was also determined from MD trajectories by first calculating the diffusion coefficient obtained using the HYDROPRO program (*112*) with default parameters and a viscosity value of $\eta$ = 0.9 mPa·s. The diffusion coefficient was converted to $R_h$ using the Stokes-Einstein equation (*113*).

Figure S12A compares $R_h$ distributions estimated from our MD simulations to hydrodynamic radii derived from SEC and DLS experiments (vertical lines). Considering some variation depending on the chosen calculation method, the mean of the MD distribution is larger by ca. 0.15 to 0.3 nm or 7% to 13%. Figure S12B provides a more direct comparison of the $R_g$ distribution calculated from our MD simulation ensembles to radii of gyration between 2.4 and 3.0 nm, determined from SAXS measurements at different p53-TAD concentrations (*114*). Independent SAXS measurements by Daughdrill et al. (*115*) yield $R_g$ values of 2.2 nm and 2.8 nm at protein concentrations of 10 and 4 mg/mL, respectively, albeit with a buffer with higher ionic strength (~70 mM vs. ~120 mM). In particular, because our simulation system contains only one isolated monomer and thus best describes a highly diluted solution, the mean radii of 2.98 nm for the WT and 2.94 nm for the P27A calculated from the simulations agree well with the measured $R_g$ values at lower protein concentrations.

Figure S12C shows a more direct comparison of calculated vs. measured SAXS spectra. SAXS spectra were computed from our MD ensembles using the program CrySol (part of ATSAS) (*116, 117*) after intensity normalization. Here, too, the SAXS spectra measured at lower intensities (1.6 and 3.2 mg/mL) agree well with the calculated spectra, which is also the case for the SAXS spectra measured by Daughdrill et al. (*115*) at 4.0 mg/mL. In contrast, deviations are seen for the spectra measured at higher concentrations of 6.4 mg/mL particularly at low wave numbers, likely due to intermolecular interactions within the sample.

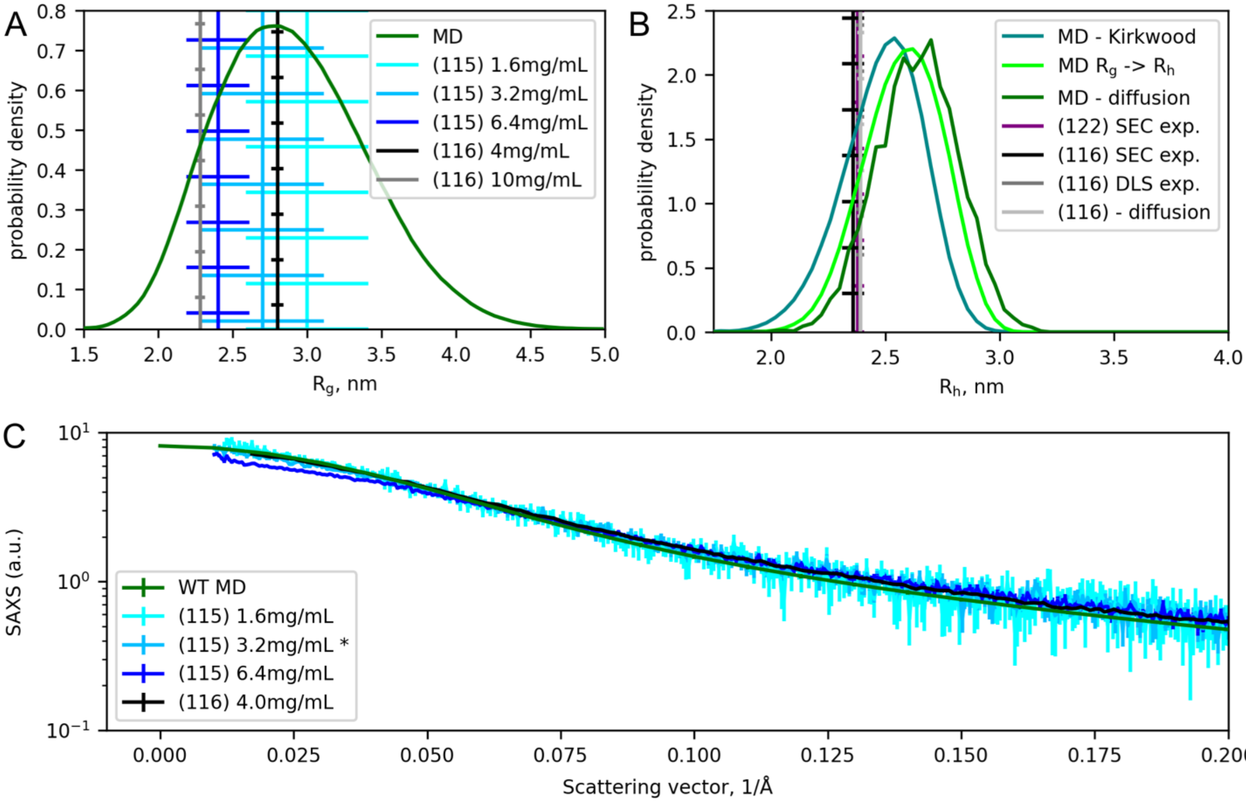

**Fig. S12. Calculated p53-TAD radii and SAXS spectra vs. SEC, DLS, and SAXS measurements.** (**A**) Comparison of $R_h$ derived from SEC and DLS experiments (vertical lines, black, shades of blue and gray) with $R_h$ distributions calculated from MD simulations (green). (**B**) Comparison of $R_g$ calculated from SAXS experiments (vertical lines) at varying protein concentrations with distributions calculated from MD simulations for p53-TAD WT (shades of green). (**C**) Direct comparison of measured SAXS spectra (black, shades of blue) with spectra calculated from MD simulations (green).

***(c) FRET distance distributions***

Fluorescence Resonance Energy Transfer (FRET) is used to measure intramolecular distances between 2-10 nm within a molecule (*118*). In these experiments, two dyes are attached to different positions of interest, and the efficiency of the energy transfer, which depends on the donor-acceptor distance, is measured. For p53-TAD, Huang et al. (*50*) used single-molecule FRET (SM-FRET) spectroscopy to measure the distance between dyes attached to residues 10 and 56 (fig. S13A, black line) of a larger p53-TAD construct comprising residues 1-91. The same authors also used time-resolved FRET (TR-FRET) spectroscopy for different, shorter peptide constructs labeled at their N- and C-terminus, specifically, segments with residues 1-17 and 14-30 (fig. S13B and C). Independently, Moses et al. (*119*) reported FRET efficiencies of a p53-TAD segment comprising residues 1-61, labeled also at the N- and C-terminus (fig. S13D). From these FRET efficiencies, we calculated the mean distance between the given residue pair using the reordered equation reported by Moses et al. (*119*).

Fig. S13 compares these measured distance distributions of the above residue pairs with distance distributions calculated from the respective $C_\alpha$ positions taken from our MD simulations. To facilitate better comparison, all distance distributions were normalized to their maxima. For all mean distances, the distances calculated from the simulations agree very well with the measured ones, except for residue pair 1-17 (fig. S13B), for which the measured average distance is ca. 2 nm smaller. Here, the missing upstream residues in the sequence of the shorter peptide used for the TR-FRET experiments might affect the detected distances compared to the full-length p53-TAD protein used for the other measurements. Also, a potential small population of cis-isomer conformations of the four proline residues within this particular sequence, which is not accounted for in the MD simulations, might cause additional shortening of this particular distance. The widths of the respective distance distributions to the measurement error are not expected to agree, nor do they, because those derived from the simulations indicate the actual distance distributions of the ensemble, whereas those reported from the FRET measurements involve milliseconds time averages as well as shot noise of the recorded FRET efficiencies; for this reason, the ensemble measurement in fig. S13D shows a much narrower distribution than those determined by single-molecule FRET. Finally, fig. S13 compares calculated $C_\alpha$-$C_\alpha$ distances to measured dye-dye distances, for which anisotropic dye-orientation distributions are known to cause deviations. Here, due to the mostly disordered nature of the structure ensemble, we assume dye-orientation anisotropy to be small, and thus also this deviation. Only minor differences are seen between the WT and the mutant P27A. Overall, given these differences between experiment and simulations, the agreement is remarkably good.

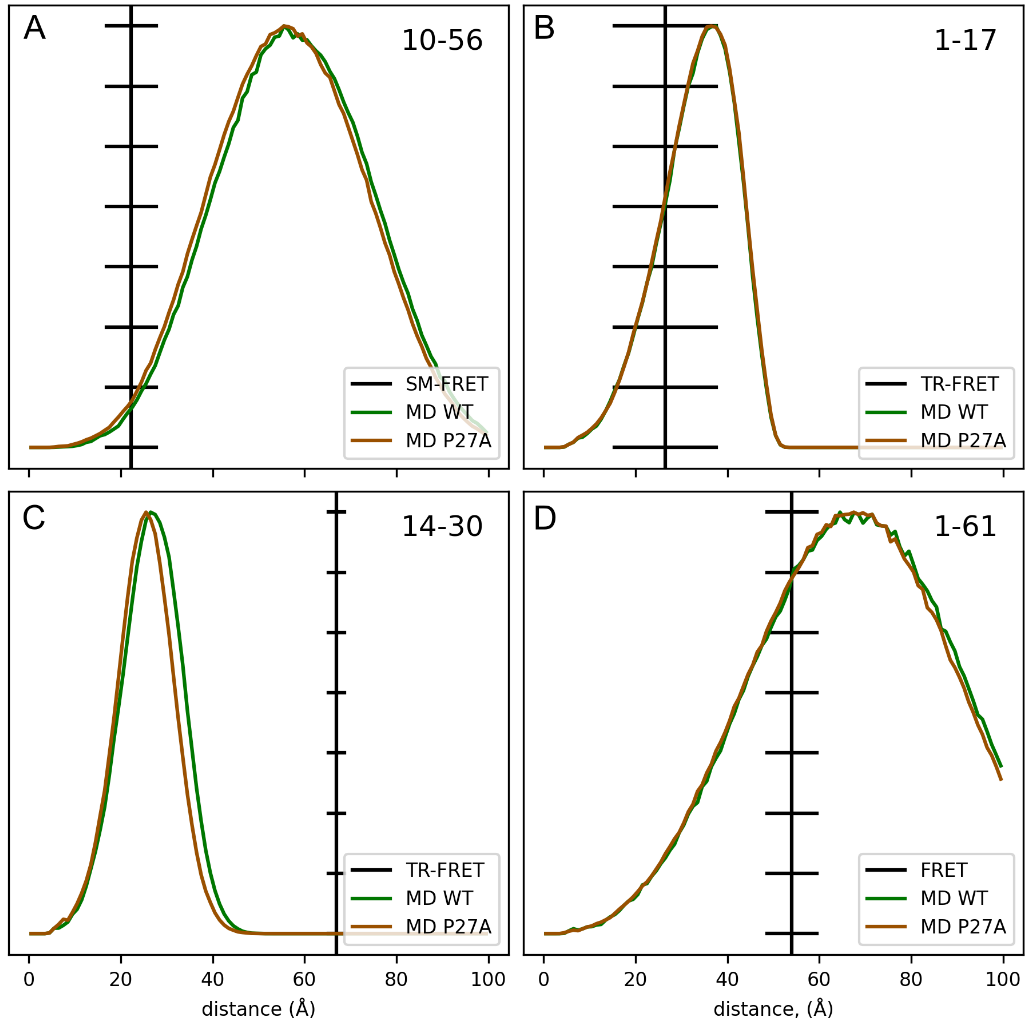

**Fig. S13. Comparison of measured FRET distances with distances calculated from our MD ensemble.** Fluorophore distances derived from SM-FRET (*50*), TR-FRET (*50*), and FRET (*119*) experiments (black solid lines) and $C_\alpha$-$C_\alpha$ distances calculated from MD simulations are shown for residue pairs (**A**) 10-56, (**B**) 1-17, (**C**) 14-30, and (**D**) 1-61 for p53-TAD WT (green) and P27A (orange).

***(d) Photoinduced Electron Transfer Fluorescence Correlation Spectroscopy (PET-FCS)***

Independent experimental information about the dynamics of p53-TAD WT (residues 1-93) was obtained by Lum et al. (*38*) using Photoinduced Electron Transfer Fluorescence Correlation Spectroscopy (PET-FCS). Briefly, these experiments rest on quenching of a fluorescence dye, attached to a specific residue, via photoinduced electron transfer upon contact with a tryptophane residue at a different position. The authors obtained a total four different constructs, which enabled them to measure the autocorrelation function (ACF) of dye-tryptophane contact formation (e.g., loop closure kinetics) for four p53-TAD segments via the fluorophore-quencher pairs at 13-23, 23-31, 31-53, and 53-60 (*38*).

To compare our atomistic simulations to these measurements, we calculated respective dye/quencher contact formation ACF curves from our MD trajectories, using the $C_\alpha$ distances between the same residue pairs as in the experiments. Having been unable to obtain the original data by Lum et al., we recalculated the measured autocorrelation functions from the published loop closure coefficients and parameters obtained from the published figures. Because the absolute scaling of the ACF is unrelated to the timescales of interest, its amplitude was manually scaled to match the calculated ACF. As can be seen in fig. S14, the ACF calculated from the simulations (green) agrees well with the measured ACF (black) for all four dye/quencher pairs and for nearly all lag times longer than 100 ns. For shorter lag times, deviations are seen, which are due to limited time resolution (> ca. 40 ns) of the experiment, a limitation that does not apply to our simulations. Indeed, when including a hypothetical 20 ns process within the ACF derived from the available experimental data, the resulting ACF (cyan) is still fully consistent with the published data (as assessed by graphical comparison of the published figures), and now also agrees with the ACF calculated from the simulations also for all lag times shorter than 100 ns.

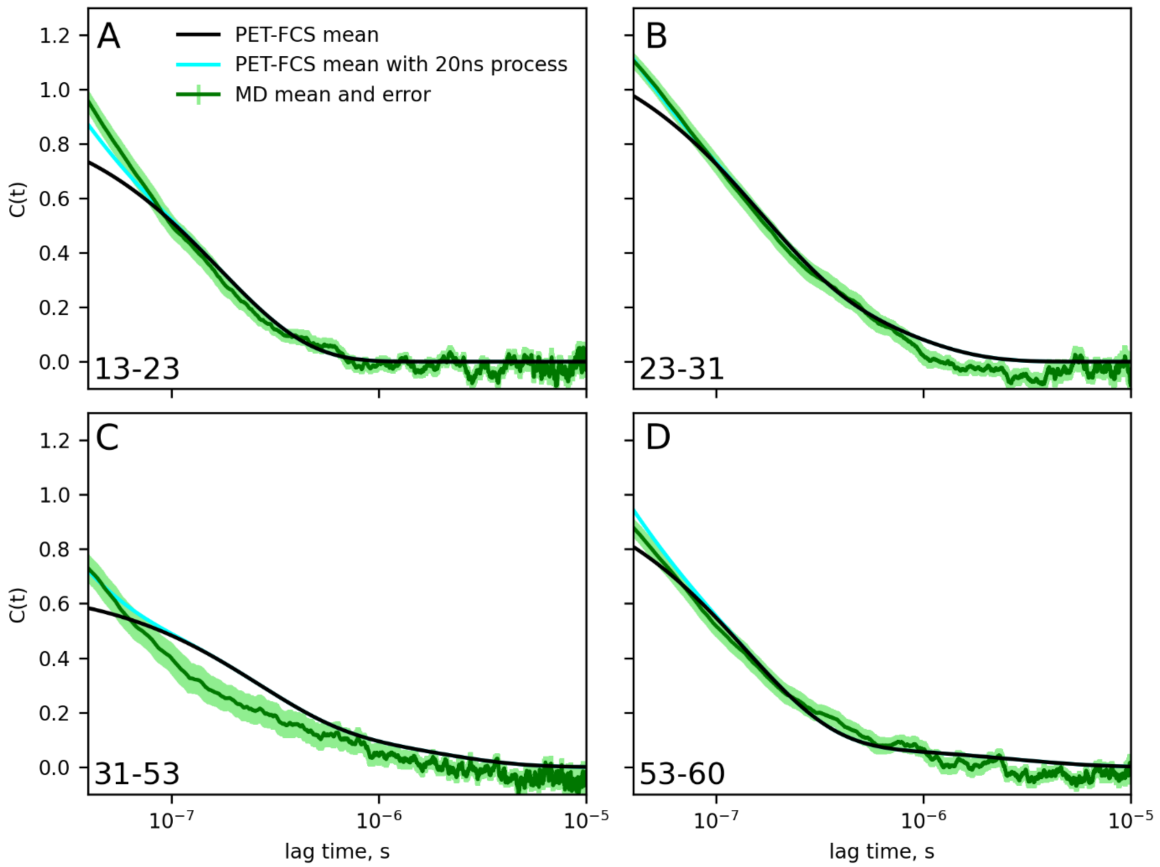

**Fig. S14. Comparison of Photoinduced Electron Transfer Fluorescence Correlation Spectroscopy (PET-FCS) measurements with our atomistic simulations.** Shown are fluorescence autocorrelation functions reproduced from published experiments in the absence of the faster dynamics observed in the simulations but inaccessible to experiment (black), experimental fluorescence autocorrelation functions including a hypothetical 20 ns dynamics component included to compensate for lack of experimental resolution (cyan), and fluorescence autocorrelation functions calculated from our MD trajectories (green) with error estimates (light green shared areas) for dye/quencher pairs at residues (**A**)13-23, (**B**) 23-31, (**C**) 31-53, and (**D**) 53-60.

#### *(e) Paramagnetic Relaxation Enhancement (PRE)*

Residue distances were also derived from Paramagnetic Relaxation Enhancement (PRE) experiments (*120*), in which the unpaired electron spin of a paramagnetic spin label enhances NMR relaxation rates of nearby nuclei, thereby providing long-range distance information. In the experiments by Lowry et al. (*121*), cysteine residues were introduced at four different positions of a p53-TAD (residues 1-73) construct, namely D7C, E28C, A39C and D61C, and the paramagnetic spin label MTSL was attached to all four mutants. $^{15}$N heteronuclear single quantum coherence (HSQC) spectra were recorded, from which resonance intensity ratios $I_{ox}/I_{red}$ were calculated (*121*) (fig. S15, black dots). We calculated corresponding intensity ratios for these four residues (fig. S15, green lines) from the $C_\alpha$-$C_\alpha$ distances taken from our MD ensemble as described by Liu et al. (*122*) and using the reported proton linewidth $R_2$ and correlation times $\tau_c$ (*121*).

Overall, the calculated ratios agree rather well with the measured ones, with a root mean squared deviation (RMSD) of 0.19, 0.33, 0.21, and 0.17 for D7C, E28C, A39C, and D61C, respectively. These RMSD values are similar or smaller than those reported by Liu et al. (*122*). The only notable discrepancy is seen for E28C between residues 35 and 55, indicating somewhat shorter distances to E28C in the experiment. This finding is in line with the above comparisons of molecular size ($R_h$ and $R_g$), which also point to a slightly more extended MD ensemble.

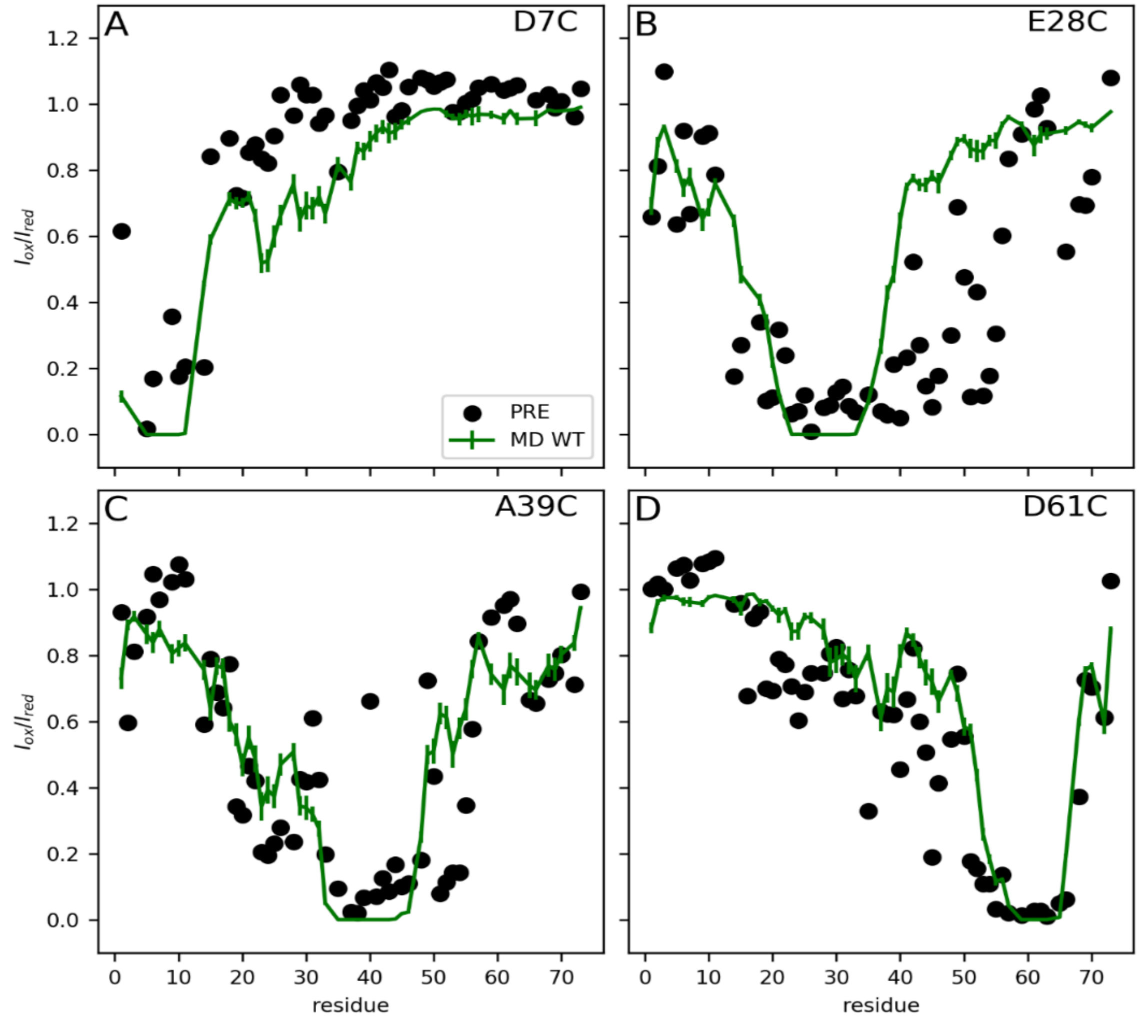

**Fig. S15. Comparison of measured PRE intensity ratios with those calculated from MD ensemble.** Resonance intensity ratios $I_{ox}/I_{red}$ determined from PRE experiments (*121*) (black dots) and intensity ratios calculated from $C_\alpha$-$C_\alpha$ distances obtained from MD simulation trajectories (green lines) for label positions (**A**) D7C, (**B**) E28C, (**C**) A39C, and (**D**) D61C. Error bars indicate errors of the mean over 30 trajectories.

**(R) Methods: Markov state model analysis**

The structural dynamics of the p53-TAD WT helix 1 and helix 2 comprising residues 18-26 and 40-53 respectively, were analyzed by building a Markov state model from a dimension-reduced representation of the 600,000 available structure snapshots using the Python library "Deeptime" (*123*). To extract the Markov states from the MD simulations, clustering was performed on the combined 600 µs trajectory set using the WT backbone dihedral angles φ and ψ of all helix 1 residues and snapshots from the MD trajectories recorded every nanosecond. For the clustering, sine and cosine values of these angles were used to avoid periodicity discontinuities, such that each residue is described/characterized by sin(φ), cos(φ), sin(ψ), and cos(ψ). Using these internal coordinates, super-positioning of snapshots of this IDP was avoided, which would have been challenging.

Dimension reduction of the resulting structure vectors was performed using time-lagged independent component analysis (TICA) (*46*), with a lag time of 10 ns. The first three independent components were used as input for subsequent KMeans clustering (*124*) (using the Deeptime implementation requesting 150 clusters, init_strategy was 'kmeans++', max_iter = 500 and fixed_seed = 13). This clustering served to assign all simulation frames to microstates, which served as input for building a hidden Markov model requesting 15 initial macrostates. These 15 initial macrostates were merged into a final number of eight macrostates, chosen empirically by inspecting spatial proximity of average structures in the TICA projection, to extract structurally and chemically unique and distinct conformations (table S3). Hydrogen bonds stabilizing an α-helix or a $3_{10}$-helix were defined by donor-acceptor distance $d$ (in Å) according to the empirical Espinosa hydrogen bonds energy estimate (in KJ/mol) $E_{HB} = -25300e^{-3.6d}$ with a cut-off hydrogen bond strength of 1 $k_B T$ (*125*). As a result, hydrogen bonds were counted for all donor-acceptor pairs closer than $d$ = 0.256 nm.

| Initial state | Final state |
|---|---|
| 4,9,10,14 | 1 |
| 3,11 | 2 |
| 1,15 | 3 |
| 8 | 4 |
| 13 | 5 |
| 2,6 | 6 |
| 5,7 | 7 |
| 12 | 8 |

**Table S3. Combining initial into final Markov states.** A total of 15 initial macrostates (left column) were combined into eight final macrostates (right column) for p53-TAD WT helix 1.

**(S) Results: Markov state model analysis of helix 2 folding dynamics**

The Markov model for p53-TAD WT helix 2 comprising residues 40-53 (fig. S16) was derived from the same MD ensemble in a similar way as described for helix 1 above, except that here five initial macrostates were requested, and after inspection as described above, no further merger was deemed necessary.

For both Markov models of helix 1 and helix 2, and for each final Markov state, a representative structure was determined at the highest density (most likely) region in the space spanned by the first two TICA components. These representative structures are depicted in Fig. 3C (helix 1) and fig. S16 (helix 2). As can be seen from the estimated free energies of the individual states in fig. S16C, helix 2 is much less stable than helix 1 (Fig. 3C) and also exhibits more diverse folding dynamics.

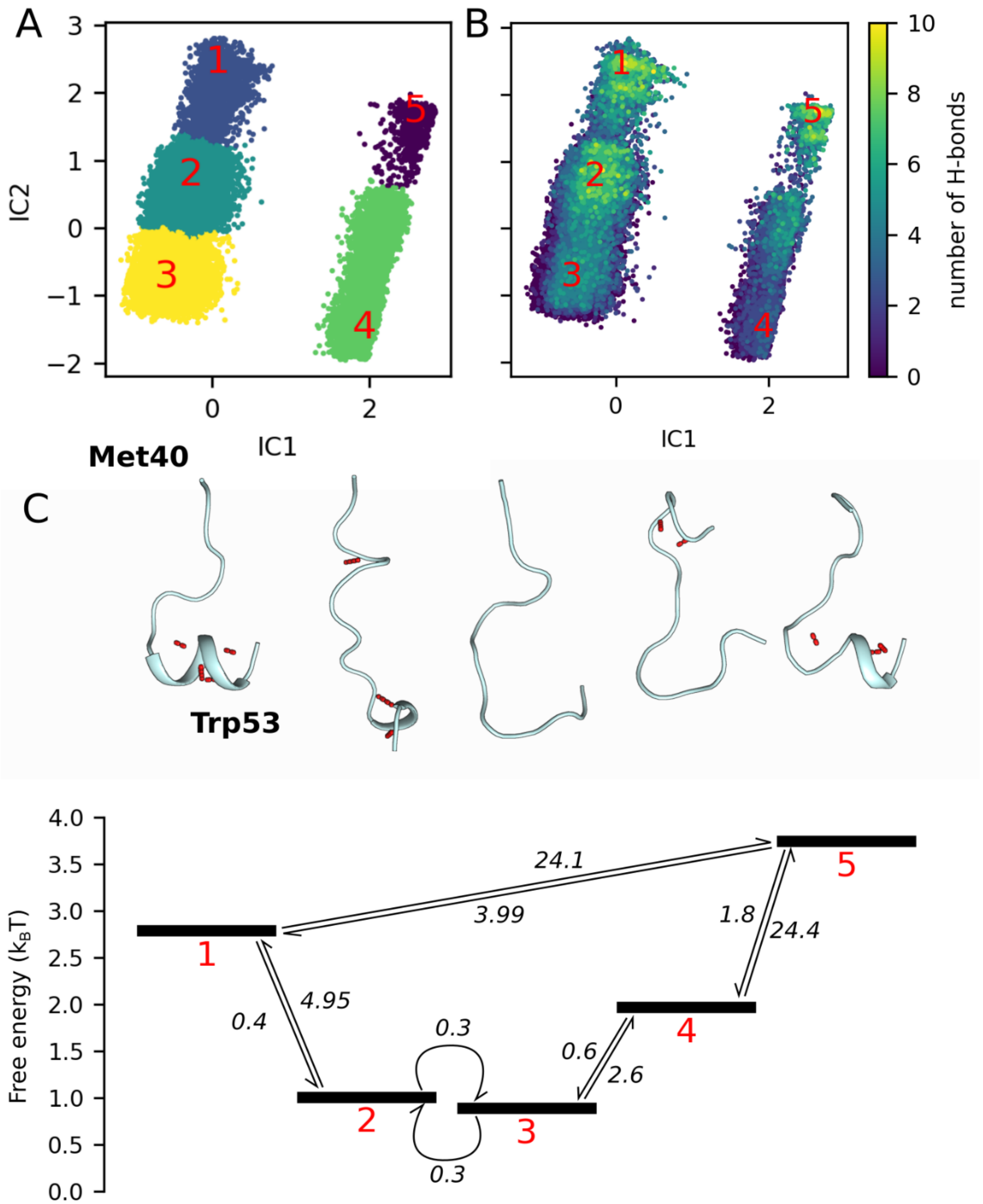

**Fig. S16. Markov state model reveals multi-timescale folding/unfolding dynamics of helix 2.** (**A**) Projection of the conformational ensemble of helix 2 (residues 40-53) onto the TICA space defined by the two collective coordinates IC 1 and IC 2 that contribute most to the intra-helical dynamics of the WT trajectories. Colors and red numbers indicate five Markov states. (**B**) The same projection as in (A), but colored by the number of intra-helical hydrogen bonds. (**C**) Representative structures of the five Markov states (intra-helical hydrogen bonds shown in red dots, top) and their free energies (black bars, below); mean first passage times of transitions between the states (arrows) are shown in μs.

**(T) Methods: Identification of metastable tertiary structure elements**

p53-TAD transiently forms tertiary structures more complex than single α-helices or β-sheets. To detect those tertiary structures that last longer than 1 μs and therefore contribute to the overall structure ensemble, we performed a $C_\alpha$-$C_\alpha$ distance fluctuation analysis on our MD trajectories (both for the WT and for the P27A mutant), which uses internal coordinates and therefore does not require orientational fitting. Specifically, for every trajectory frame, intramolecular distances between all pairs of $C_\alpha$ atoms were calculated. Residue pairs separated by no or only one residue along the sequence were excluded from this analysis, because their mutual $C_\alpha$-$C_\alpha$ distances showed only small fluctuations and thus do not provide much information on metastable tertiary structures.

For all trajectories, the size of the distance fluctuations was quantified over time and for each residue pair via standard deviations, calculated by averaging over a sliding window of 100 ns width (containing 1000 frames). For each window position in time, and for each residue, the standard deviation of the distances to all other residues was calculated and then sorted. From the obtained list of standard deviations, the $10^{th}$ smallest one, indicating the $10^{th}$ least fluctuating distance, was used as a tertiary structure indicator. Here, the $10^{th}$ distance was chosen empirically, to best reflect tertiary structures larger than a typical two turn α-helix or a smaller β-sheet. Of note, this indicator was also chosen because it does not require the involved residues to be on a continuous sequence segment and, therefore, it also identifies folds formed by distant residues.

Figure 3E shows an example of the resulting fluctuation map, showing averaged distance fluctuations between 0.1 nm (yellow) and 0.5 nm (dark blue), plotted for every residue (y-axis) over time (x-axis). In this plot, regions extending vertically over several residues and horizontally over more than 1 μs are easily identified, e.g., the two large yellow regions involving residues 15 to 53 lasting from 2 to 6 μs, and the one involving residues 47 to 66 between 20 and 24 μs. This analysis also recovers the faster tier 1 folding dynamics of the two helices of p53-TAD, which are sufficiently stable to show up as rapidly fluctuating yellow bands between residues 18-26 (helix 1) and residues 40-53 (helix 2). For an automated scan of all MD trajectories, a fluctuation cutoff of 0.15 nm was chosen, and 93 transient tertiary structure elements were identified lasting longer than 1 μs.

One might expect the transient tertiary structures indicated by the MD simulations and tier 0 dynamics to be detected on a NOESY spectrum. The fact that not one but many different transient tertiary structures are seen also explains why these are not seen as peaks in the NOESY spectra. Indeed, inspection of the expected positions in measured NOESY spectra (see Methods C) did not reveal any signals beyond the noise level. Although the population of tertiary structures, taken together, is large enough to evoke a low-frequency component in the RD profiles at least for P27A, the population of each of the many different structures is below 0.3%, whereas an estimated population of at least 1% would be required to generate a visible NOESY cross-peak.

#### (U) Methods: Analyses and comparison to polymer models

The extensive sampling provided by our atomistic simulations of the p53-TAD also allowed us to characterize the local fast (tier 2) dynamics of this IDP from a polymer model perspective, e.g., in terms of persistent lengths and end-to-end distances, which for a polymer chain – and depending on the polymer model used – are connected (*126*). For, e.g., a wormlike chain model, the (average squared) end-to-end distance $R_e$ reads

$$\langle R_e^2 \rangle = 2 l_p L \left( 1 - \frac{l_p}{L} \left( 1 - e^{-\frac{L}{l_p}} \right) \right) \text{ , (21)}$$

where $l_p$ is the persistence length, $L$ = $bN$ is the contour length, i.e., the length of one polymer unit ($b$ = 0.38 nm) multiplied by their number $N$. From our MD trajectories, $\langle R_e^2 \rangle$ was calculated by time- and trajectory averaging over all distances between all pairs of $C_\alpha$ atoms separated by given contour length (i.e., number of residues). Fitting Eq. (21) to the resulting curve (fig. S17A) yielded a persistence length of $l_p$ = 1.0 nm.

Residue-specific persistence lengths were obtained similarly, except that averaging over different sequence positions was omitted, such that $\langle R_e^2 \rangle$ was obtained as a function of 'start' residue. Similar fits to Eq. (21) as above provided residue-position-resolved persistence length, characterizing deviations from a simple homopolymer (fig. S17B). As can be seen, shorter persistence lengths $l_p$ are obtained for proline rich segments, e.g., Pro12-Pro13.

We also characterized the relation between persistence length $l_\mathrm{p}$ and radius of gyration $R_\mathrm{g}$, which is given by the polymer scaling law (*52*)

$$\langle R_g \rangle = \sqrt{\frac{2 l_p b}{(2\nu+1)(2\nu+2)}} N^{\nu} \text{ ,}$$

where $N$ is the number of residues in the different tested segments, $l_\mathrm{p}$ and $b$ are as defined above, and $\nu$ is the scaling exponent. A value of $\nu$ = 0.5 indicates a 'Flory random coil' (*127*), smaller values a compact and larger values a more extended ensemble. Similarly, as for $\langle R_e^2 \rangle$, $R_\mathrm{g}$ was calculated for all protein segments of length between $N$ = 6 and 30 residues, averaging over time, trajectories, and all possible segment positions in the peptide. From the fit shown in fig. S17C a markedly shorter persistence length of 0.31 nm and a scaling exponent of 0.66 was obtained; the latter indicating an expanded coil state

(*52*), similar to an excluded volume chain model (0.588) (*128*). For comparison, $R_g$ of the full protein was calculated similarly, but was not used for the above fit.

Alternatively, the persistence length was also estimated (fig. S17D) by fitting single and double exponential functions, respectively, to the normalized orientation autocorrelation function of the vectors connecting $C_\alpha$ atoms separated by N>2 residues (*126*).

$$C(N) = Ae^{-\frac{Nb}{l_{p,1}}} + (1-A)e^{-\frac{Nb}{l_{p,2}}},$$

where *A* is a prefactor, *N* is the number of polymer units (residues) separating the vectors, with the same unit length *b* = 0.38 nm as above. Persistence length ($l_p$) from the single exponential fit is 1.19 ± 0.01 nm, whereas from the double exponential 0.84 ± 0.01 nm and 5.06 ± 0.26 nm, for $l_{p,1}$ and $l_{p,2}$, respectively.

The fast reorientational tier 2 dynamics were also characterized by the running average root mean squared deviation (RMSD)

$$RMSD(\tau) = \sum_{i=1}^{N_{atoms}} \sqrt{\frac{1}{N_{atoms}} \left(x_i(t) - x_i(t+\tau)\right)^2},$$

averaged over all trajectories (fig. S17E). With increasing lag time, and for ca. 10 ns over nearly two timescale decades, the structural deviation increase closely follows a power law with an exponent of 0.39; thereafter the RMSD saturates. This reorganization time of ca 10 ns agrees with that measured for other polymers by single-molecule FRET (*14*).

Complementing this analysis of structural dynamics at a rather detailed level, we finally calculated the autocorrelation times of the radius of gyration $R_g$ as well as of the end-to-end distance of the full p53-TAD chain. Fitting a double exponential function

$$C(\tau) = Ae^{-\frac{\tau}{\tau_1}} + (1-A)e^{-\frac{\tau}{\tau_2}}$$

to the respective autocorrelation functions that were calculated similarly as above, two timescale values ($\tau_1$ and $\tau_2$) were obtained for each of the two observables (fig. S17F), namely 22.0 and 406.8 ns for $R_g$ and 18.5 and 349.8 ns for the end-to-end distance.

### (V) Results: Analyses and comparison to polymer models

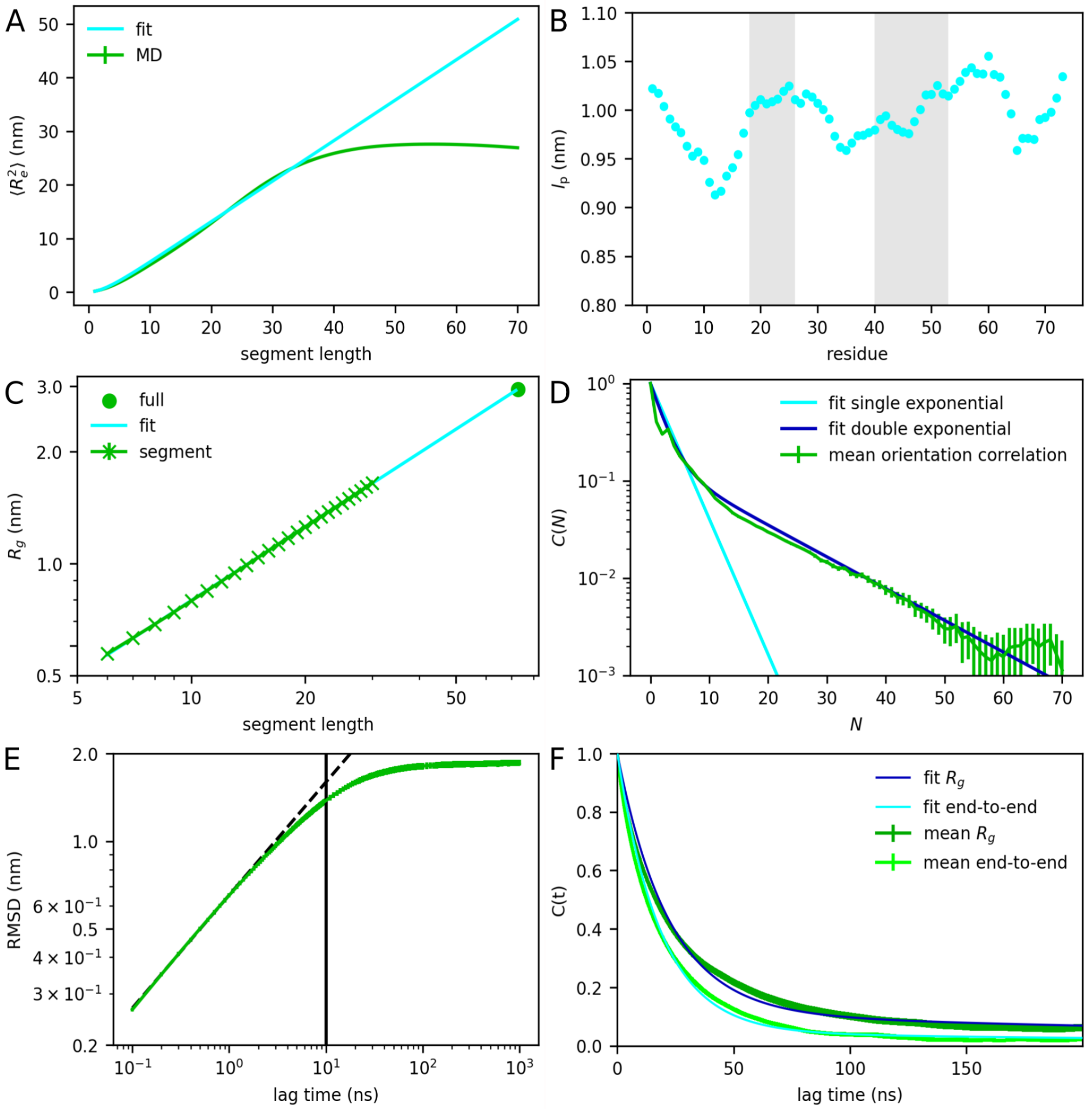

**Fig. S17. Analysis of p53-TAD from a polymer physics perspective.** (**A**) Mean squared end-to-end distance for increasing peptide segment length scanned over the protein (green) and a fitted scaling law function (cyan); the estimated error of the mean is smaller than the line width. (**B**) Persistence length (cyan) derived from the mean squared end-to-end distance of a fixed segment length, starting from each residue; gray areas indicate helix 1 (18-26) and helix 2 (40-54) residues. (**C**) Radius of gyration calculated for increasing protein segment lengths (green crosses) and for the full peptide (green circle); the cyan line shows a scaling law fit; the estimated standard error is smaller than the line width. (**D**) Orientation autocorrelation function of vectors of consecutive $C_\alpha$ atoms with increasing separation (*N*); error bars show the error of the mean; a single (cyan) and a double (blue) exponential function was fitted to obtain the persistence length. (**E**) Average increase of root mean squared deviation (RMSD) over increasing lag time; error bars show the error of the mean; the vertical black line indicates the estimated beginning of the RMSD saturation. (**F**) Time autocorrelation functions of the full p53-TAD radius of gyration (dark green) and end-to-end distance (light green); error bars show the standard error; dark blue and cyan lines show double exponential function fits.

**(W) Results: Comparison of different force fields**

Although force fields for explicit water MD simulations of IDP have been compared and assessed before (*87, 89, 88*), no consensus has emerged regarding which force field describes the structure ensemble and dynamics of IDPs most accurately. One reason is that the achieved accuracy seems to depend also on the studied IDPs.

We have therefore determined specifically for the p53-TAD IDP the accuracy of combinations of the two force fields that ranked highest in previous assessments with three different water models. In particular, 10 MD simulations covering a total length of 10 μs each were performed using (a) Amber99sbws (*85*) with the TIP4P2005s (*86*) water model, (b) CHARMM36m (*88*) with the default TIP3P water model and (c) CHARMM36m with the OPC water model (*129*), which recently was found to be similarly accurate for several systems (unpublished data).

For each trajectory and each residue, the SDF was calculated via Eqs. 2 and 3, from which the transverse cross-correlation rate constants $\eta_{xy}$ and tumbling timescales $\tau_c$ were calculated using Eqs. 4-9 as described above, as well as rate constants $R_1$ and $R_2$ from Eqs. 10-15 and NOE values from Eq. 16. These calculated observables were averaged, errors of the mean were estimated from their standard deviations, and the resulting values for each residue were compared to NMR measurements (fig. S18).

As can be seen, for all five observables the best accuracy is achieved by the Amber99sbws + TIP4P2005s force field (green lines), which is generally closest to the measured values and best reproduces the peaks between residues 20 and 26 (helix 1). In contrast, the CHARMM36m + TIP3P (red lines) combination results in too fast dynamics, as reflected by short tumbling times $\tau_c$, while those for CHARMM36m + OPC (purple lines) are too long, indicating too slow dynamics particularly of the highly flexible residues between p53-TAD helix 1 and helix 2.

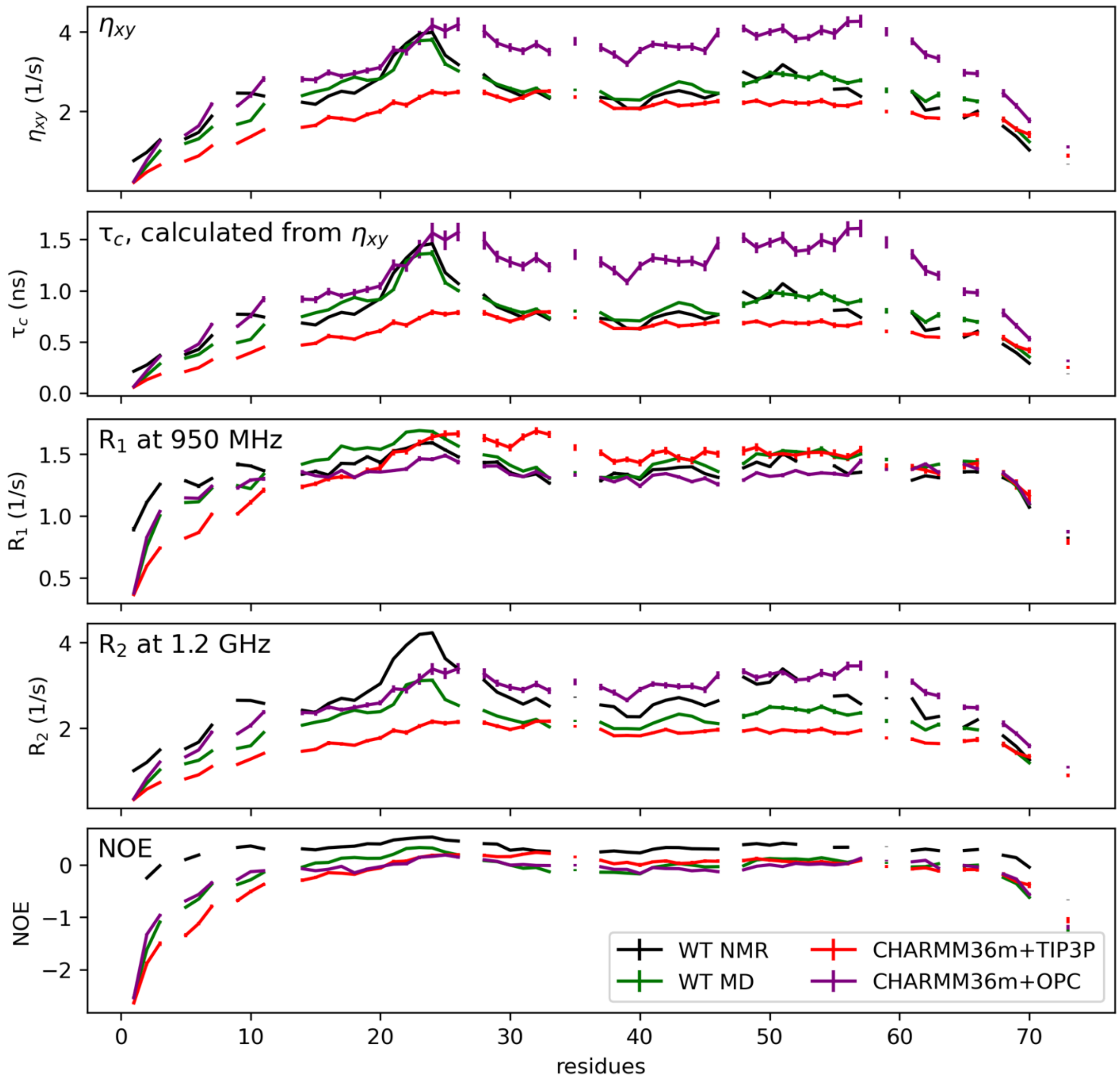

**Fig. S18. Assessment of force field accuracies.** The five panels show NMR measurements (black lines) of (from top to bottom) transverse cross-correlation rate constants $\eta_{xy}$, tumbling timescale $\tau_c$, rate constants $R_1$ (at a magnetic field of 950 MHz), $R_2$ (at 1.2 GHz), and NOE values. In each panel and for each residue, these are compared to respective values calculated from MD simulations using the three different force field-water combinations Amber99sbws + TIP4P2005s (green), CHARMM36m + TIP3P (red), and CHARMM36m + OPC (purple). Error bars indicate standard error of the mean estimated from standard deviations, and adjacent residues are connected by lines to guide the eye.

**(X) Results: Convergence analysis**

Convergence of the MD simulations was assessed by comparing observables calculated separately from three consecutive 20 µs blocks of the 60 µs long P27A trajectories. Chemical shift-based RD profiles were calculated for the full trajectories as described above and were averaged over 30 independent trajectories using the first, second and third 20 µs block of each trajectory. Figure S19 compares these averaged RD spectra calculated for each of the blocks (lines) and errors of the mean (shaded areas) calculated from the respective standard deviations. As can be seen, almost all averaged RD profiles are within their respective error ranges except for very few cases such as Trp23, indicating sufficient convergence.

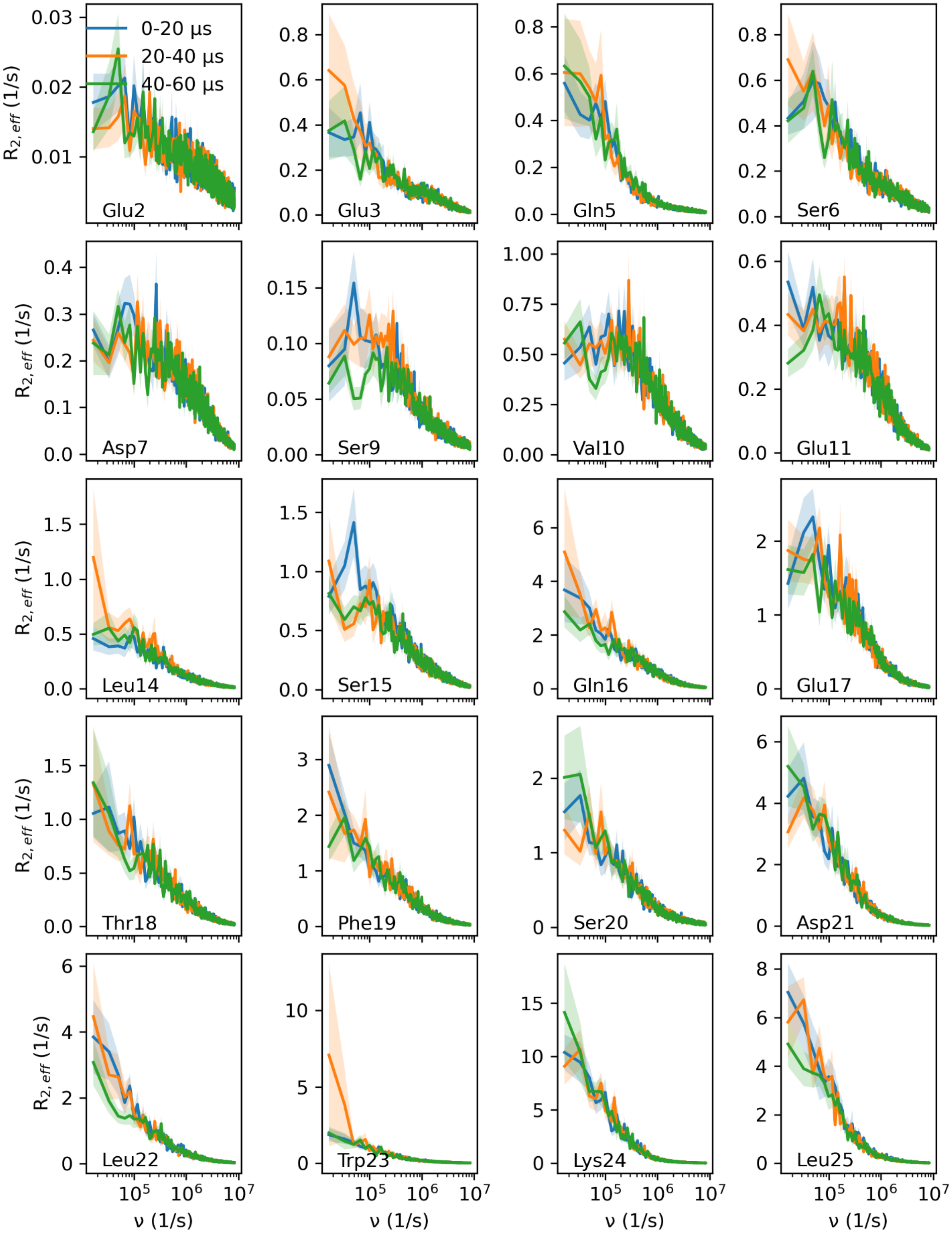
0-20 μs
20-40 μs
40-60 μs
$R_{2,eff}$ (1/s)
ν (1/s)
Glu2
Glu3
Gln5
Ser6
Asp7
Ser9
Val10
Glu11
Leu14
Ser15
Gln16
Glu17
Thr18
Phe19
Ser20
Asp21
Leu22
Trp23
Lys24
Leu25

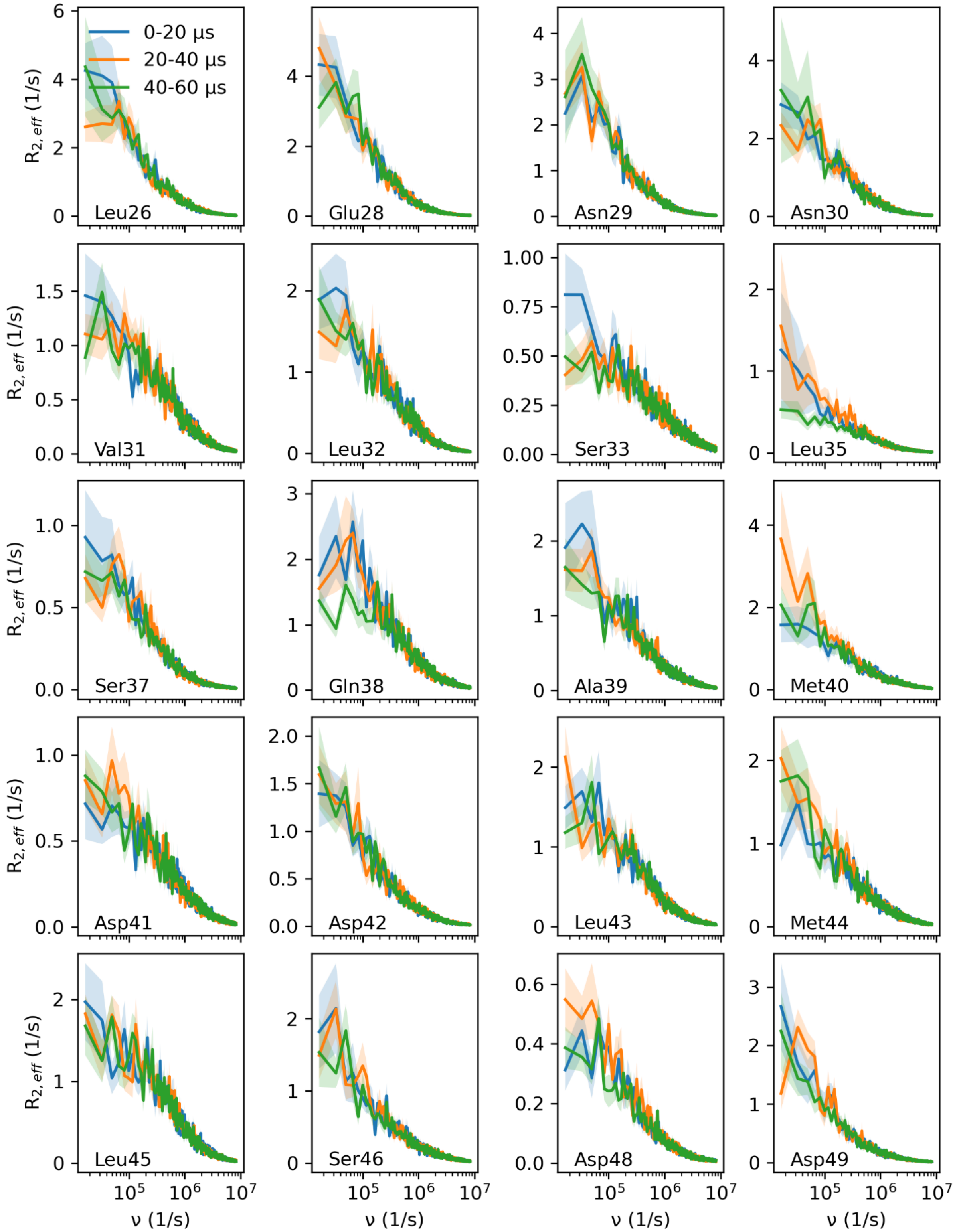
0-20 µs
20-40 µs
40-60 µs
$R_{2,eff}$ (1/s)
ν (1/s)
Leu26
Glu28
Asn29
Asn30
Val31
Leu32
Ser33
Leu35
Ser37
Gln38
Ala39
Met40
Asp41
Asp42
Leu43
Met44
Leu45
Ser46
Asp48
Asp49

1116

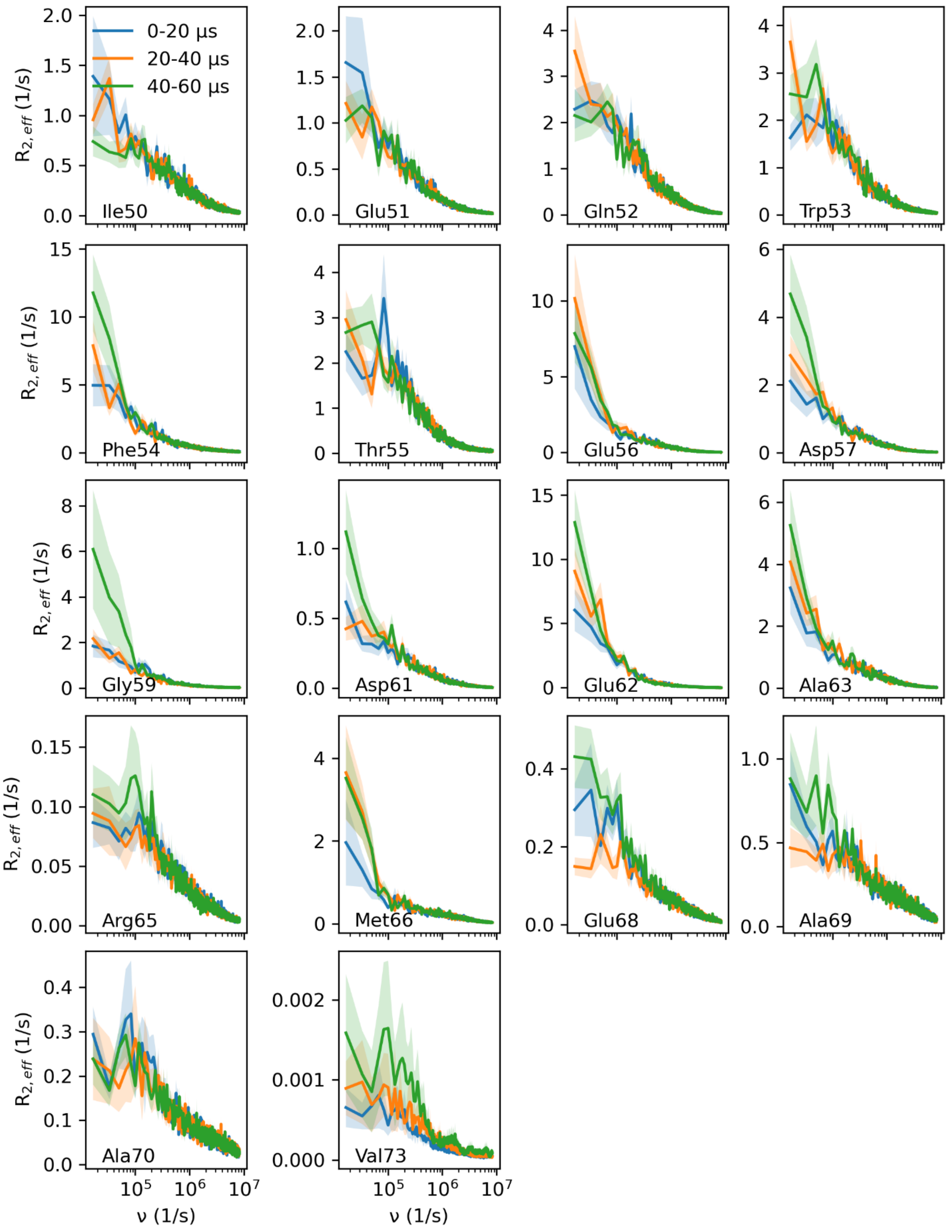

**Fig. S19. MD simulations convergence analysis.** Shown are, for each residue, averaged RD profiles (lines) calculated from three non-overlapping 20 µs blocks of the 30 x 60 µs long trajectories of the P27A mutant. Errors of the mean are shown as shaded areas.

### (Y) Results: RD profiles for the Measles $N_{TAIL}$ peptide

To test if the broad spectrum of time scales observed for p53-TAD is unique to this IDP or, rather, is a more general feature of IDPs, we have additionally calculated and analyzed RD profiles from MD simulations of the measles virus peptide $N_{TAIL}$ (residues 399-525). Compared to p53-TAD, $N_{TAIL}$ has a quite different charge distribution and no sequential similarity to p53-TAD. The details of these simulations are described in (*130*). Briefly, the AMBER99SB-disp force field and the TIP4P water model (*89, 131*) were used with $Na^+$ and $Cl^-$ ions corresponding to an ion concentration of 150 mM. Six MD simulations of 10 µs each were used.

The RD profiles of $N_{TAIL}$ were calculated and a stretched CPMG equation (Eq. 18) was fitted using the same procedure as described above for p53-TAD. Figure S20 compares the resulting stretching parameters $\gamma$ for each residue, which describes how much the profiles are stretched with respect to the analytical two-state CPMG profile (Eq. 17). Here, $\gamma$ = 1 corresponds to an unmodified version of the CPMG equation, and increasingly smaller values indicate more extended RD profiles and, hence, superpositions of more exchange processes with timescales distributed over an increasingly broader frequency range. As can be seen, on average both IDPs show similar $\gamma$-parameters of about 0.6, scattering over similar ranges between ca. 0.25 and 0.95 for the individual residues, with by far the most being markedly smaller than 1. This result suggests that the observed multi-timescale dynamics are rather independent of the sequence, length, and physico-chemical properties of a particular IDP, and rather a more general phenomenon. Clearly extensive MD simulations of other IDPs will be helpful to further establish this finding.

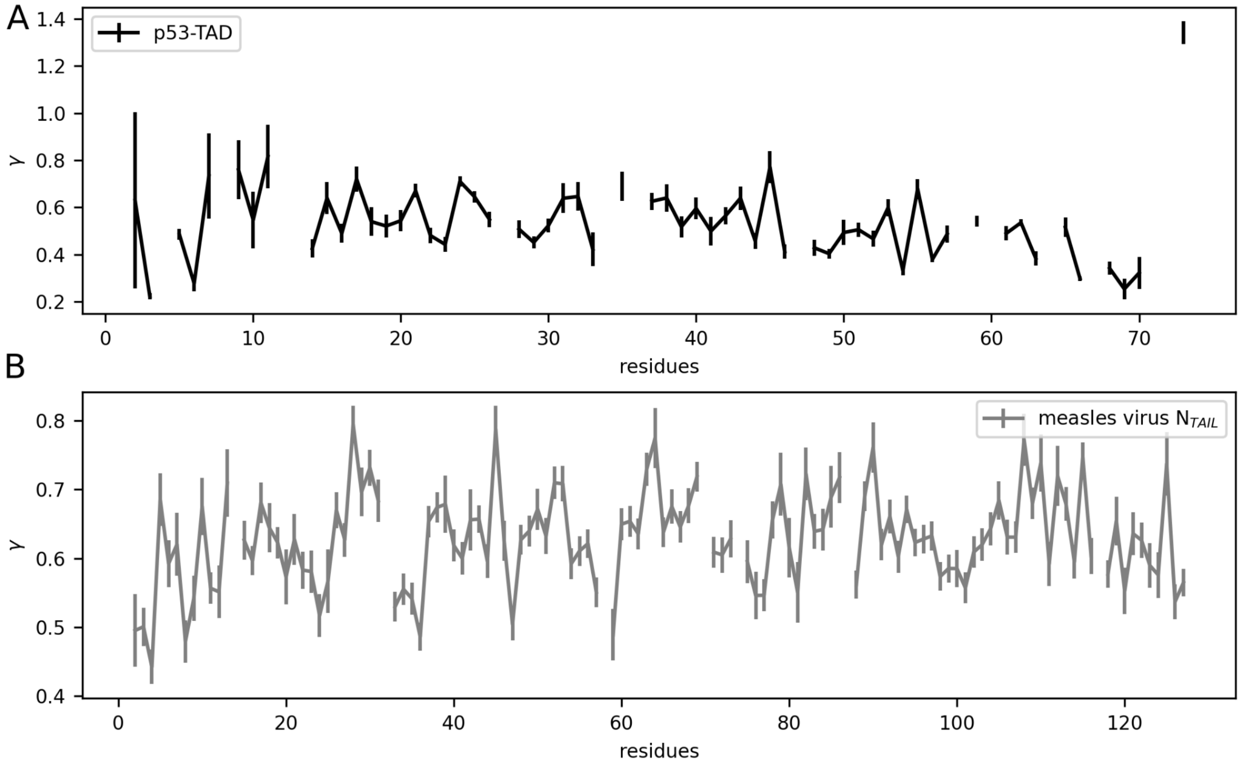

**Fig. S20. Different IDPs show similar multi-timescale dynamics.** Shown are, for each residue, stretching parameters $\gamma$ obtained from fits of the stretched CPMG equation (Eq. 18) to RD profiles calculated from a set of MD trajectories of (**A**) p53-TAD and (**B**) the Measles virus $N_{TAIL}$. The symbols show the mean of the Bayesian posterior distributions and error bars indicate its standard deviation.

**(Z) p53-TAD tertiary structures resembling protein structure elements**

To test if the metastable tier 0 tertiary structures observed in our atomistic simulations also occur in protein structures, we searched the complete RCSB Protein Data Bank (*54*). Specifically, we searched for protein structure fragments, the shape of which is most similar to these tier 0 tertiary structures. To this end, all 12 (WT) and 52 (P27A) stable tertiary structures detected by $C_\alpha$ distance fluctuations (see SM (T)) were used as a search query in “Structure Similarity Search” mode with parameters "strict_shape_match" and scoring_strategy = "structure", which identifies structures that are similar in shape (i.e., electron density overlap (*132*)), irrespective of sequence or specific atom types. Both complete tertiary structures as well as 11 residue long subsets sliding along the sequence (WT: 126; P27A: 460) were used for the search. For the complete structures, the best 10 search results (based on their score) were recorded; for the 11 residue long subsets, only the best matching result was recorded. Structure fragments consisting of a helix only were discarded. All identified protein structure fragments were compared to the respective metastable tier 0 tertiary structure using the software PyMOL (*133*). Structure similarity was quantified using root mean squared distances (RMSDs) calculated using the PyMOL "super" method, which also accepts sequentially unrelated structures for comparison.

For the complete WT tertiary structures, four similar structure fragments were identified with RMSD values of approximately 0.3 nm; fig. S21A shows an example. For the complete P27A tertiary structures, three fragments were identified (fig S21B). Using the 11 residue long sliding sub-structures, two structure fragments resembling tertiary structures seen for the WT were identified (e.g., fig. S21C), and six different structure fragments seen for the P27A mutant. Fig. S21D shows an example of the latter with an RMSD of 0.08 nm.

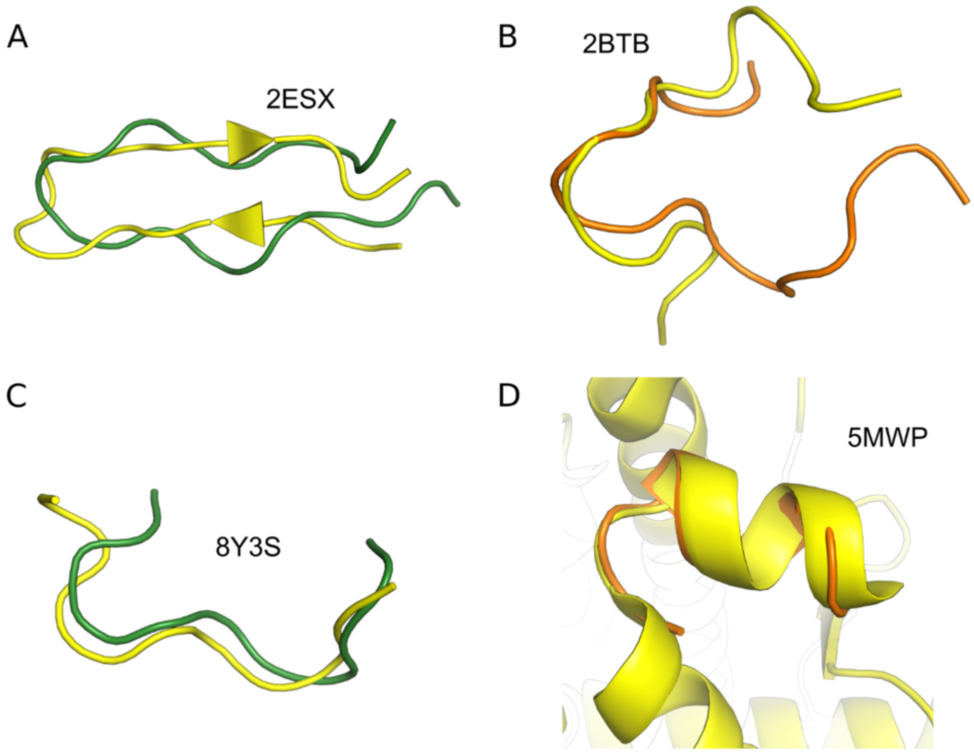

**Fig. S21. Examples of transient tertiary tier 0 structures compared to similar structure fragments identified in the RCSB Protein Data Bank.** (**A**) Cartoon representation of a tertiary structure observed for the p53-TAD WT (green) and structure fragment (yellow) of the V3 region of gp120 of the JR-FL HIV-1 strain with PDB-ID indicated in the figure; (**B**) P27A tertiary structure (orange) and N-terminal residues 1-15 of the human band 3 peptide (yellow); (**C**) p53-TAD WT tertiary structure (green) and residues 53-63 of the human keratin 19 head domain (yellow); (**D**) P27A tertiary structure (orange) and residues 22-32 of the human mineralocorticoid receptor (yellow).

### Description of Movies S1, S2, S3

**Movie S1: Super slow motion movie of a sample MD simulation trajectory of the p53 P27A mutant.** Shown is the trace of the backbone (grey), with helix1 highlighted in cyan, helix 2 in orange, and the N-terminal in yellow. A total of 100 ns is shown (2.5 ns/second).

**Movie S2: Slow motion movie of a sample MD simulation trajectory of the p53 P27A mutant.** Coloring as in S1; a total of 3 μs is shown (25 ns/second).

**Movie S3: Movie of a sample MD simulation trajectory of the p53 P27A mutant.** Coloring as in S1; a total of 10 μs is shown in slow motion (250 ns/second).
